# Specific decorations of 17-hydroxygeranyllinalool diterpene glycosides solve the autotoxicity problem of chemical defense in *Nicotiana attenuata*

**DOI:** 10.1101/2020.08.26.267690

**Authors:** Sven Heiling, Lucas Cortes Llorca, Jiancai Li, Klaus Gase, Axel Schmidt, Martin Schäfer, Bernd Schneider, Rayko Halitschke, Emmanuel Gaquerel, Ian Thomas Baldwin

**Affiliations:** Max Planck Institute for Chemical Ecology, Department of Molecular Ecology, Hans Knöll Straße 8, 07745 Jena, Germany; Max Planck Institute for Chemical Ecology, Department of Biochemistry, Hans Knöll Straße 8, 07745 Jena, Germany; Max Planck Institute for Chemical Ecology, Research Group Biosynthesis/NMR, Hans Knöll Straße 8, 07745 Jena, Germany; Centre for Organismal Studies Heidelberg, Im Neuenheimer Feld 360, 69120 Heidelberg, Germany; Institut de Biologie Moléculaire des Plantes, CNRS UPR 2357 Université de Strasbourg, 12 rue du Général Zimmer, 67084 Strasbourg, France

## Abstract

17-hydroxygeranyllinalool diterpene glycosides (HGL-DTGs) are abundant and potent anti-herbivore defense metabolites in *Nicotiana attenuata* whose glycosylation and malonylation biosynthetic steps are regulated by jasmonate signaling. To characterize the biosynthetic pathway of HGL-DTGs, we conducted a genome-wide analysis of uridine diphosphate glycosyltransferases (UGTs) and identified 107 members of family-1 UGTs. Tissue-specific time-course transcriptional profiling revealed that the transcripts of three UGTs were highly correlated with two HGL-DTG key biosynthetic genes: geranylgeranyl diphosphate synthase (Na*GGPPS)* and geranyllinalool synthase (Na*GLS*). NaGLS’s role in HGL-DTG biosynthesis was confirmed by virus-induced gene-silencing. Silencing the UDP-rhamnosyltransferase, *UGT91T1,* indicated its role in the rhamnosylation of HGL-DTGs. *In vitro* enzyme assays revealed that UGT74P3 and UGT74P4 use UDP-glucose for the glucosylation of 17-hydroxygeranyllinalool (17-HGL) to lyciumoside I. *UGT74P3* and *UGT74P5 stably* silenced plants were severely developmentally deformed, suggesting a phytotoxic effect of 17-HGL. Applications of synthetic 17-HGL and silencing of these UGTs in HGL-DTG-free plants confirmed the phytotoxic effect of 17-HGL. Feeding assays with *Manduca sexta* larvae revealed the defensive functions of the glucosylation and rhamnosylation steps in HGL-DTG biosynthesis. Glucosylation is a critical step that contributes to the metabolites’ defensive function and solves the autotoxicity problem of this potent chemical defense.

## Introduction

In the course of evolution, plants developed versatile defense strategies against biotic stresses, such as herbivores and pathogens. Among these defensive strategies are physical and chemical barriers such as trichomes (Fordyce and Agrawal, 2001), volatiles that attract predators (Kessler and Baldwin, 2001; Halitschke et al., 2008) and a wide array of defensive specialized metabolites. Many small molecules produced as part of a plant’s specialized metabolism function as direct defenses by being toxic, repellent, or anti-nutritive for herbivores of different feeding guilds and degrees of specialization; compounds with a broad spectrum of toxicity are likely to be of greater defensive value against a greater diversity of attackers. Noteworthy examples are glucosinolates, alkaloids, and terpenoids. However, these broad-spectrum toxins force the producers to solve the "toxic waste dump problem" of chemical defense, as many direct defense compounds are generally cytotoxic and can damage the tissues of non-adapted producers.

Plants have evolved numerous ways of solving this “toxic waste dump problem” for broad-spectrum chemical defenses. One frequently used solution is glycosylation, which is one of the most prevalent and widespread biochemical modifications contributing to the structural and functional diversity of specialized metabolites in plants. Incorporation of sugar molecules into small lipophilic metabolites can regulate storage/localization of defensive metabolites (Gachon et al., 2005; Yadav et al., 2014) and change their bioactivity by detoxifying phytotoxic intermediates. For example, for the defensive deployment of hydrogen cyanide (Gleadow and Moller, 2014) or steroidal saponins (Mylona et al., 2008), plants store these toxins as glycosides, sometimes in particular compartments, away from lytic enzymes that liberate the active toxins in response to the tissue damage that frequently accompanies herbivore and pathogen attack. Similarly, glucosinolates are compartmentalized away from the myrosinases that rapidly hydrolyze them to toxic isothiocyanates and other biologically active products (Matile, 1980; Halkier and Gershenzon, 2006). Steroidal alkaloids are also safely stored as glycosides and silencing *GAME-1*, a UDP-galactosyltransferase responsible for the glycosylation of these defense compounds in tomato, results in the accumulation of the aglycone, α-tomatidine with severe developmental consequences (Itkin et al., 2011). Glycosylation of the saponin, hederagenin in *Medicago truncatula* (Naoumkina et al., 2010), and of avenacin A-1 in *Avena sativa* (Mylona et al., 2008) are other examples which point to a similar chemical sequestration role exerted by glycosylation.

Diterpene glycosides (DTGs) are a diverse compound class whose members are often associated with phytotoxic activities (Macias et al., 2008) and have potent anti-herbivore resistance/deterrence effects. For example, the abundance of monomers and dimers of capsianosides is correlated with thrips resistance in pepper (Macel et al., 2019) and 17-hydroxygeranyllinalool diterpene glycosides (HGL-DTGs) in *Nicotiana* species have been shown to function in resistance against larvae of the specialist herbivore *Manduca sexta* (Lou and Baldwin, 2003; Jassbi et al., 2008; Heiling et al., 2010) and the generalist herbivore, *Heliothis virescens* (Snook et al., 1997).

HGL-DTGs constitute an abundant and structurally diverse group of specialized metabolites whose function in plants is poorly understood (Heiling et al., 2010). HGL-DTGs occur in the aboveground tissues of the native diploid tobacco, *Nicotiana attenuata* (Heiling et al., 2010) as well as other *Nicotiana* species (Shinozaki et al., 1996; Snook et al., 1997; Lou and Baldwin, 2003; Jassbi et al., 2006; Heiling et al., 2010; Jassbi et al., 2010; Poreddy et al., 2015; Heiling et al., 2016), several other solanaceous genera including *Capsicum* (Izumitani et al., 1990; Hashimoto et al., 1997; Iorizzi et al., 2001; Lee et al., 2006, 2007, 2008; Lee et al., 2009) and *Lycium* (Terauchi et al., 1995; Terauchi et al., 1997b, a; Terauchi et al., 1998b; Terauchi et al., 1998a; Roda et al., 2003), and the Asteraceae, *Blumera lacera* (Akter et al., 2016). HGL-DTGs consist of an acyclic 17-hydroxygeranyllinalool aglycone, which is conjugated at its C-3 and C-17 hydroxyl groups to different sugar moieties, such as glucose and rhamnose. These sugars can be further glycosylated at the C’-2, C’-4 or C’-6 hydroxyl groups and acetylated or malonylated at the C’-6 hydroxyl group of the glucose(s) (Taguchi et al., 2005; Yu et al., 2008, Heiling et al., 2010; Jassbi et al., 2010; Heiling et al., 2016). To date, 45 HGL-DTGs, which differ in their sugar or malonyl decorations, have been putatively annotated or identified in *N. attenuata* (Figure 1) (Heiling et al., 2016). However, it is unclear which of the many different HGL-DTGs or which structural components of HGL-DTGs are responsible for the observed deterrent (Jassbi et al., 2006) and resistance effects against different herbivores (Snook et al., 1997). For example, the geranyllinalool precursor is known to be insecticidal in pine wood and the same compound can be found in the defensive secretions of the termite *Reticulitermes lucifugus* (Baker et al., 1982; Lemaire et al., 1990). Jassbi and colleagues (Jassbi et al., 2010) suggested that the 17-HGL aglycone is responsible for the feeding-deterrent characteristics of HGL-DTGs, but this hypothesis has not been tested.

**Figure 1:**
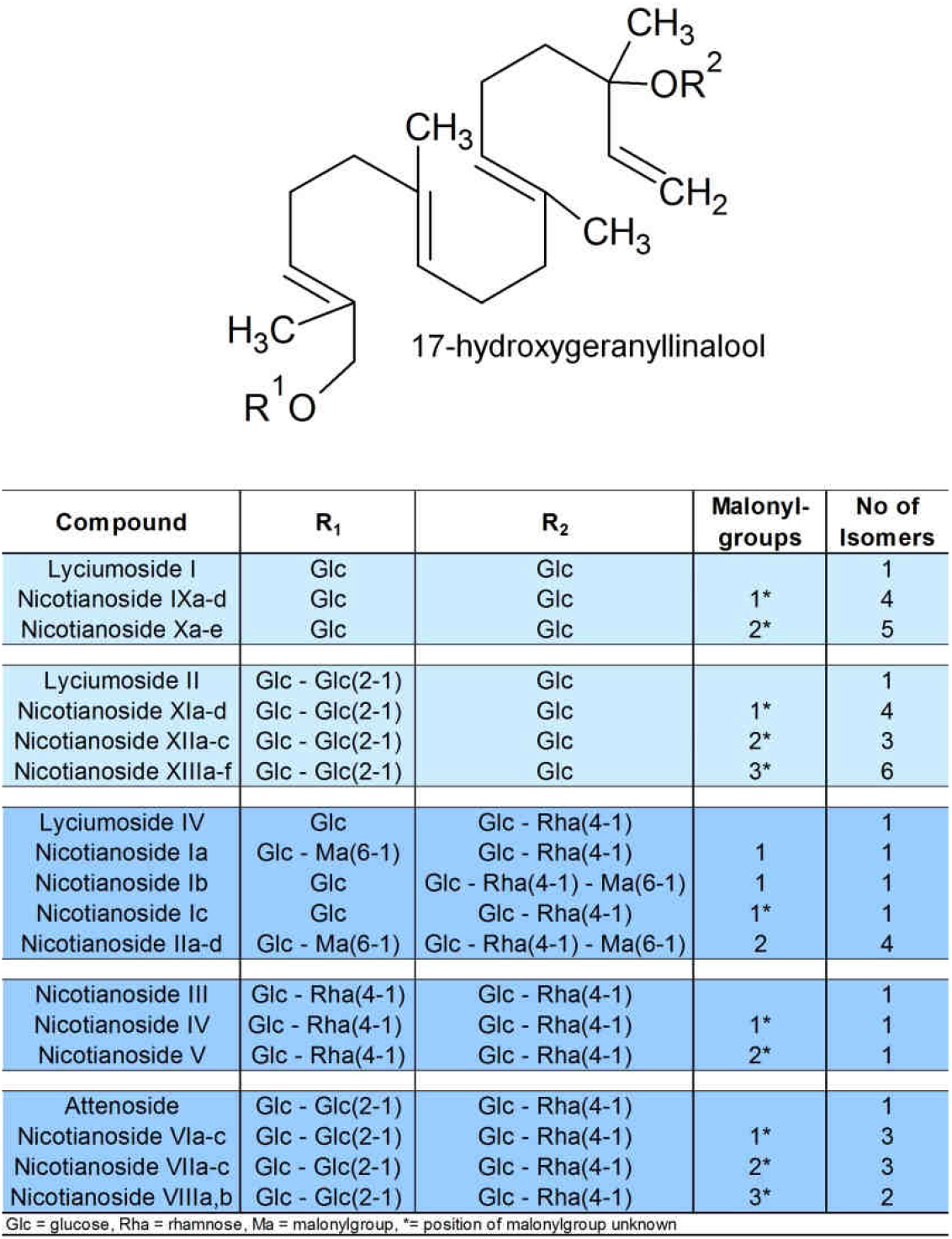
HGL-DTG biosynthetic pathway. Components of the diverse HGL-DTGs structures previously identified and annotated in the leaves of *N. attenuata* that differ with respect to their sugar and malonyl group composition.

While the chemical identity and anti-herbivore effects of HGL-DTGs have been the focus of major investigations, less is known about the possible phytotoxic effects of HGL-DTGs and how producing plants cope with these phytotoxic effects. This is, in part, because we know very little about the enzymes required for the biosynthesis of these metabolites. The 17-HGL aglycone is derived from the condensation of three five-carbon units of isopentenyl-pyrophosphate (IPP) and dimethylallyl-pyrophosphate (DMAPP) to produce the diterpenoid precursor, geranylgeranyl-pyrophosphate (GGPP; (Ohnuma et al., 1998; Dewick, 2002)). This reaction is catalyzed by a plastidial geranylgeranyl pyrophosphate synthase (GGPPS; (Jassbi et al., 2008; Heiling et al., 2010)). Formation of GGPP is followed by its allylic rearrangement by a geranyllinalool synthase (GLS; (Falara et al., 2014)) that produces the tertiary alcohol geranyllinalool. However, the enzymes necessary for further hydroxylation and glycosylation steps, which determine most of the structural diversity of HGL-DTGs, remain to be characterized. Li and colleagues (Li et al., 2018) identified and silenced a malonyltransferase (NaMAT1), which is responsible for the malonylation of HGL-DTGs. However, all malonyl moieties of HGL-DTGs are rapidly lost when leaves are ingested by *M. sexta* larvae, suggesting that the malonylation of HGL-DTGs does not play a central role in anti-herbivore defense (Poreddy et al., 2015). Interestingly, disruption of the uniform malonylation patterns of HGL-DTGs leads to a specific reduction in the floral style lengths of *N. attenuata* flowers (Li et al., 2018). This shows that specific decorations of a plant’s specialized metabolites can play a crucial, but poorly understood, role in plant development. Malonylation of specialized metabolites, such as flavonoids or phenolic glucosides, is a common phenomenon and it has been shown that malonyl moieties alter the molecular properties of these compounds can enhance their solubility in water (Heller and Forkman, 1994) or sequester compounds to molecular compartments, such as vacuoles (Taguchi et al., 2010). Whether other intermediates in the HGL-DTG pathway are similarly influential for plant development remains to be explored.

To evaluate the defensive value of glycosylation and the potential (auto)toxicity of the 17-HGL aglycone for both plant cells and insect herbivores, we investigated the UDP-glycosyltransferases (UDP-GTs) responsible for the glycosylation of HGL-DTGs. To this end, we analyzed the tissue- and herbivory-specific co-regulation of the 107 predicted UDP-GTs in *N. attenuata* UDP-GTs with two bait genes previously characterized for their involvement in the biosynthesis of 17-HGL, which pinpointed on three novel UGT candidates. RNAi silencing of these UGTs by RNAi approach in *N. attenuata* and the related HGL-DTG-producing sympatric diploid species *N. obtusifolia* (syn. *N. trigonophylla* Dunal) (Heiling et al., 2016) as well as enzyme assays with recombinant proteins identified that UGT74P3 and UGT74P4 are responsible glucosylation of the C-3 and C-17 hydroxyl-groups to form lyciumoside I, while UGT91T1 in *N. attenuata* and NoUGT91T1-like in *N. obtusifolia* are rhamnosyltransferases in the HGL-DTG pathway. As summarized in one of our previous studies (Heiling et al., 2017), all rhamnosylated HGL-DTG identified so far possess rhamosyl moieties attach to glucose ones, pointing to the conclusion that rhamnosylation requires prio glucosylation. Finally, genetic manipulations revealed that glycosylation of the 17-HGL aglycone by UGT74P3 is crucial to prevent toxic effects to the plant as well as contributes to the metabolites` defensive function during the attack by *M. sexta* larvae

## Results

### Putative identification of UGTs responsible for HGL-DTG biosynthesis

To identify UGTs, a genome-wide survey of *N. attenuata* was performed and a total of 107 putative UGT sequences containing the PSPG motif at the C-terminus were detected (Supplemental Figure 1). These were phylogenetically characterized (Supplemental Figure 2, Supplemental Table 2), their amino acid sequence examined (Supplemental Figure 3, Supplemental Table 1, 3, 4) and their expression analyzed in a full-transcriptome microarray dataset obtained from leaf and root tissues collected at several time-points following simulated leaf herbivory (wounding and application with oral secretions [OS]) as described in supplemental method file 1.

HGL-DTGs are secondary metabolites of *N. attenuata*, whose biosynthetic steps of glycosylation are regulated by the defense phytohormone, jasmonic acid (Heiling et al., 2010). Genes involved in a shared biological process tend to be co-expressed in large-scale expression datasets due to the fact that they are frequently under the control of a common regulatory network (Saito et al., 2008). Following this rationale, we identified UGT candidates putatively responsible for the biosynthesis of HGL-DTGs by exploring gene co-expression in tissue-specific (local and systemic leaves as well as root tissue) transcriptomes analyzed at 1, 5, 9, 13, 17 and 21 h after the simulated herbivory treatment. For the resulting compendium consisting of 150 microarray expression profiles, we calculated the Pearson Correlation Coefficients (PCC) for 150 microarrays of the expressed UGTs with previously identified genes of the HGL-DTG pathway (*NaGGPPS* and *NaGLS*, Supplemental Table 5a/b). Transcript accumulation of three UGTs was increased in response to wounding and simulated herbivory and significantly correlated with *GGPPS* (*UGT91T1* – PCC = 0.823, *UGT74P3* – PCC = 0.608 and *UGT74P5* – PCC = 0.684) and *GLS* (*UGT91T1* – PCC = 0.899, *UGT74P3* –PCC = 0.872 and *UGT74P5* – PCC = 0.868). The PCC value between *GLS* and *GGPPS* transcript levels was 0.799 (Supplemental Figure 4). Furthermore, we compared the expression levels between shoot and root tissues of the above genes. Consistent with the absence of HGL-DTG accumulation in *N. attenuata* roots, transcript accumulation was 50-fold and 3225-fold lower in roots compared to leaves for *GGPPS* and *GLS*, respectively. The three candidate UGTs showed a similar profile with 2190-, 127-, and 20-fold lower root transcript levels of *UGT91T1*, *UGT74P3,* and *UGT74P5,* respectively. Focusing on the 5 h time point collected from systemic leaves, we detected that *GGPPS* (increased 6-fold), *GLS* (increased 8-fold), *UGT91T1* (increased 7-fold), *UGT74P3* (increased 8-fold) and *UGT74P5* (increased 8-fold) were all highly up-regulated, relative to untreated control and mechanical wounding conditions, in response to the simulated herbivory treatment (Supplemental Figure 4 and 5, Supplemental Data 1).

A phylogenetic alignment with functionally characterized UGTs (Supplemental Figure 6) showed a close relationship of UGT74P3 and UGT74P5 protein sequences with two previously characterized diterpene glucosyltransferases (SrUGT74G1 – steviol glycoside glucosyltransferase from *Stevia rebaudiana* and CsGIT2-crocetin glucosyltransferase from *Crocus sativus*). UGT91T1 showed a close phylogenetic relationship to three of the very few functionally characterized flavonoid rhamnosyltransferases (CmF7G12RT – flavonoid-1, 2-rhamnosyltransferase from *Citrus maxima*, GmF3G6R – flavonol-3-O-glucoside-α-1, 6-*L*-rhamnosyltransferase from *Glycine max* and PhA3ART – anthocyanidin-3-O-glucoside-α-1, 6-*L*-rhamnosyltransferase from *Petunia x hybrida*).

Based on the co-expression with known genes of the HGL-DTG pathway, the localization of high expression levels in leaf tissues and their phylogenetic relationships, we selected these three candidate UGTs for further characterization.

### Virus-induced gene-silencing reveals a role of three UGTs in HGL-DTG production

Using a well-established transient virus-induced gene-silencing (VIGS) approach (Saedler and Baldwin, 2004), we first examined the consequences of independently silencing the three candidate UGTs for HGL-DTG production. Seventeen days after inoculation with *Agrobacterium tumefaciens* harboring the appropriate constructs, leaf transcript abundance was reduced by 98.5% for *UGT91T1*, 85% for *UGT74P3* and 94.3% for *UGT74P5* relative to the empty vector controls (EV, pTV00) (Supplemental Figure 7). Co-silencing resulted in reduced transcript abundance of *UGT74P3* and *UGT74P5* in both, pTVUGT74P3 and pTVUGT74P5 plants (Supplemental Figure 8).

To test the hypothesis that *UGT91T1*, *UGT74P3* and *UGT74P5* control the glycosylation steps in the HGL-DTG pathway, we analyzed the leaf metabolome by UPLC/TOF-MS of the different VIGS plants under control conditions (lanolin, Lan) or after treatment with a lanolin paste containing methyl jasmonate (Lan + MeJA), known to strongly induce the *de novo* production of HGL-DTGs (Heiling et al., 2010). We putatively identified and annotated HGL-DTGs using a previously established rapid de-replication and identification workflow, which is based on high resolution MS data analysis and a library of putative and identified HGL-DTGs (Heiling et al., 2016).

Levels of rhamnosylated HGL-DTGs, most particularly of lyciumoside IV and attenoside, were reduced compared to pTV00 controls, while the non-rhamnosylated HGL-DTGs lyciumoside I and lyciumoside II increased in the HGL-DTG chemotype of pTVUGT91T1 VIGS plants (Figure 2A, Supplemental Data 2). This shift was more pronounced after the Lan + MeJA treatment: in transiently transformed pTVUGT91T1 plants, most rhamnosylated HGL-DTGs were barely detectable and non-rhamnosylated HGL-DTGs were dramatically increased compared to pTV00 controls (Figure 2A and B). Furthermore, we detected the 17-HGL aglycone as well as several novel compounds in pTVUGT74P3 and pTVUGT74P5 VIGS plant profiles (Figure 2A). We defined this class of compounds as intermediate HGL-DTGs and our MS annotation workflow suggested compounds with only one or two sugar moieties, malonylated or not at either the C-3 or C-17 hydroxy-group of the aglycone.

**Figure 2:**
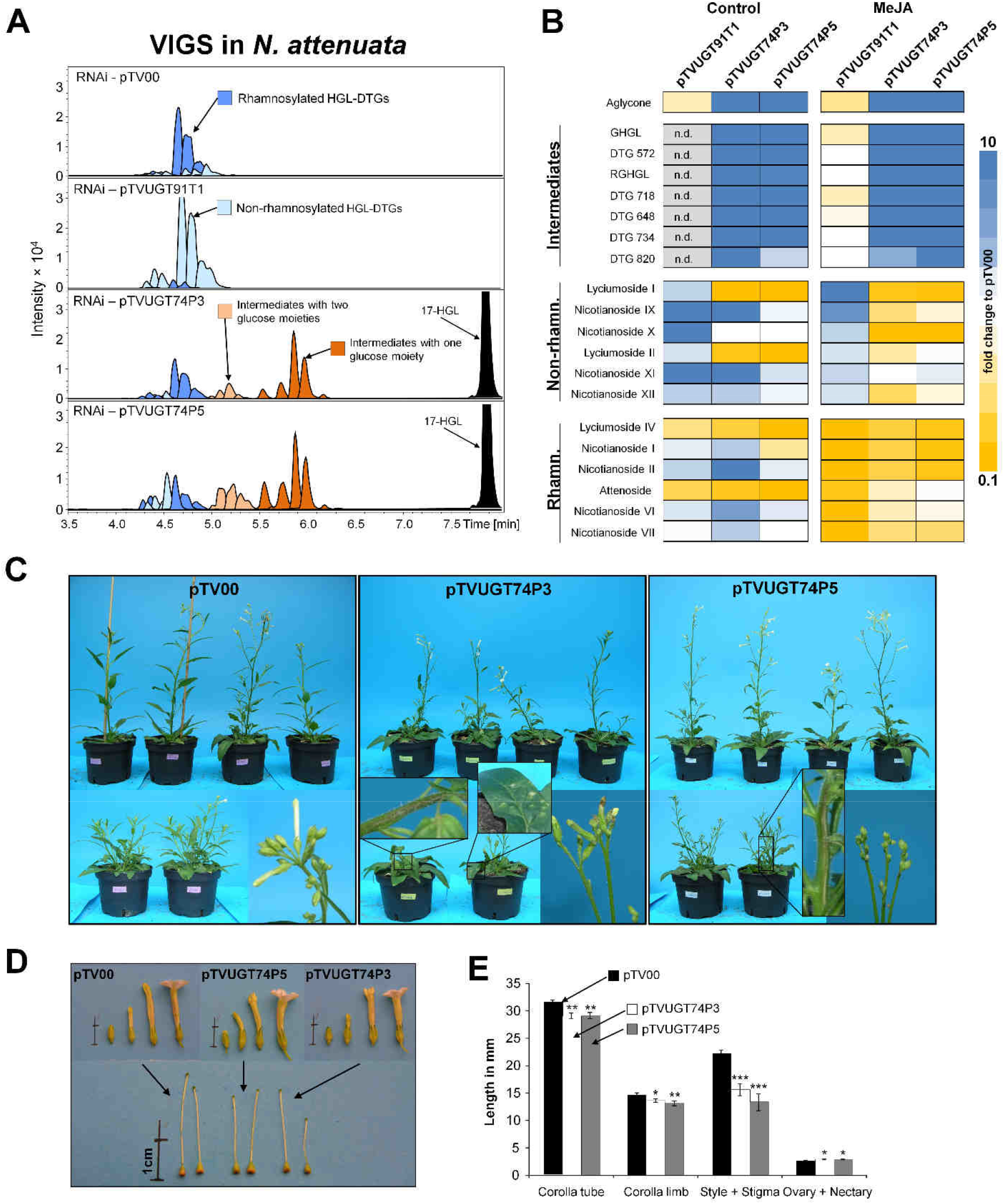
Metabolite profiling and morphologies of *N. attenuata* plants transiently-silenced in the expression of HGL-DTG-predicted UGTs by virus-induced gene silencing (VIGS) A) Extracted ion chromatograms (EIC) for identified HGL-DTGs in leaves of 37 day-old elongated *N. attenuata* plants silenced in *UGT91T1*, *UGT74P3* or *UGT74P5* transcript accumulation as well as in the empty vector controls (pTV00). HGL-DTGs were categorized into rhamnosylated, non-rhamnosylated and intermediates with one or two glucose moieties to facilitate visualization. The 17-hydroxygeranyllinallool (17-HGL) aglycone was only detected in transiently-silenced pTVUGT74P3 and pTVUGT74P5 lines. B) Heatmap visualization of the patterns of deregulation in control plants or in plants treated with 150 μg methyl jasmonate (MeJA) in 20 μL lanolin paste (N=5). The color gradient visualizes fold changes in individual HGL-DTGs for each of the VIGs constructs compared to the average in the pTV00 empty vector VIGS plants. C) Morphological alterations observed in pTV00, pTVUGT74P3 and pTVUGT74P5 transiently-transformed plants ranged from necrotic spots and tissues to necrotic apical meristem and flower buds frequently stalled in the opening process. Additional phenotypic details are provided in supplemental Figures S7a-c. D) and E) Morphological alterations of the corolla tube, corolla limb, style, ovary and nectary (N=20). Asterisks indicate significant differences between the empty vector control and pTVUGT74P3 or pTVUGT74P5 VIGS plants (*P ≤ 0.05, ** P < 0.01, *** P < 0.001).

Furthermore, we searched for orthologues to these three UGTs in *N. obtusifolia*, a sympatric tobacco species to *N. attenuata.* We identified No*UGT74P4* (95% homology to *UGT74P3*), No*UGT74P6* (93% homology to *UGT74P5*) and NoUGT91T1-like (92% homology to *UGT91T1*). Using VIGS, we inoculated *A. tumefaciens* harboring the appropriate constructs into 25-day-old *N. obtusifolia* plants and detected transcript reductions of 91.2% in No*UGT91T1-like*, 98.5% in No*UGT74P3* and 93.1% in pTVNo*UGT74P6,* 14 days after inoculation (Supplemental Figure 9).

pTVNoUGT91T1-like VIGS plants showed an overall decrease in rhamnosylated HGL-DTGs compared to pTV00 controls and an increase in non-rhamnosylated compounds (Figure 3A and 3B, Supplemental Data 3). In pTVNoUGT74P4 and pTVNoUGT74P6 VIGS plants compared to pTV00 controls, the abundance of intermediate HGL-DTGs as well as the 17-HGL aglycone was increased and non-rhamnosylated HGL-DTGs were decreased. Silencing both glucosyltransferases in *N. obtusifolia* by means of a double construct resulted in the same phenotypic alterations, namely the appearance of the intermediate HGL-DTGs and of the 17-HGL aglycone. Ten novel intermediate HGL-DTGs were putatively identified based on the de-replication workflow for HGL-DTGs. A detailed MS analysis and putative structure description of *N. attenuata* and *N. obtusifolia* HGL-DTGs can be found in the Supplemental Table 6 and 7 and Supplemental Figure 10a and b and 11a-c.

**Figure 3:**
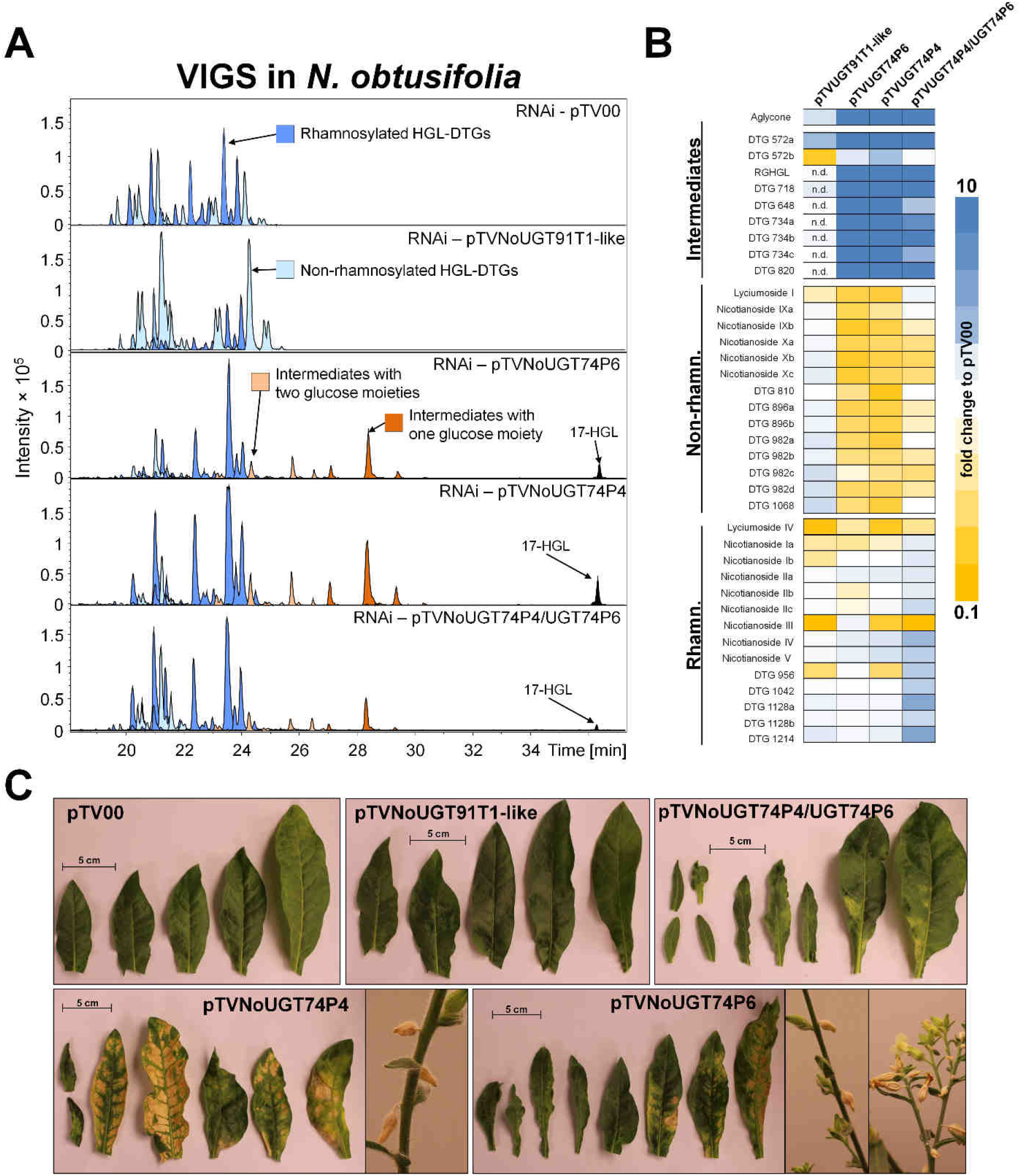
Metabolite profiling and morphologies of *N. obtusifolia* plants transiently-silenced in the expression of HGL-DTG-predicted UGTs by VIGS. A) Extracted ion chromatograms (EIC) for the identified HGL-DTGs of 37 day-old elongated *N. obtusifolia* plants silenced in No*UGT91T1-like,* No*UGT74P4* and No*UGT74P6* transcript accumulations as well as in the empty vector (pTV00) controls. Rhamnosylated, non-rhamnosylated and intermediate HGL-DTGs with one or two glucose moieties were color-categorized as in Fig. 3. The 17-HGL aglycone was only detected in pTVNoUGT74P4, pTVNoUGT74P6 and the double construct pTVNoUGT74P4/UGT74P6 VIGS plants. B) Heatmap visualization of deregulations in the leaf HGL-DTG profiles of transiently-transformed pTVNoUGt91T1-like, pTVNoUGT74P4, pTVNoUGT74P6 or pTVNoUGT74P4/UGT74P6 plants (N=5). The color gradient visualizes fold changes in individual HGL-DTGs for each of the VIGS constructs compared to the average in the pTV00 empty vector VIGS plants. C) Morphological alterations in pTV00, pTVNoUGT91T1-like, pTVNoUGT74P4, pTVNoUGT74P6 or pTVNoUGT74P4/UGT74P6 ranged from necrotic spots to a high percentage of stalled flower buds. Additional phenotypic details are reported in supplemental Figure 9.

### 17-HGL glucosylation activities of UGT74P3/P4 and UGT74P5 proteins

The co-silencing in both UGT74P3 and UGT74P5 resulted in strong overlap in the metabolic alterations of the HGL-DTG profiles produced by pTVUGT74P3 and pTVUGT74P5 plants and suggested that at least one of these enzymes might play a concerted role in the formation of lyciumoside I and lyciumoside II through glucosylation of the C-3 or C-17 hydroxyl-group of 17-HGL. HGL-DTGs typically contain one D-Glc moiety at the C-3 and one at the C-17 hydroxyl group. Lyciumoside II derives from lyciumoside I by the attachment of a second D-Glc at the C’-2 position of the C-17 D-Glc. To test this hypothesis, we analyzed the *in vitro* activity of the UDP-GT enzymes when recombinantly produced in *E. coli*. UDP-glucose was used as sugar donor with a mixture of all six isomers of synthesized 17-HGL (Supplemental Figure 12) as substrates. NaUGT74P3 and its *N. obtusifolia* homologue, NoUGT74P4, readily used UDP-glucose as a donor producing several novel products with a mass-to-charge ratio of *m/z* 491.3003 [M+Na]^+^ corresponding to a mono-glucosylated 17-HGL and two additional products with *m/z* 653.3509 [M+Na]^+^ corresponding to lyciumoside I and another di-glucosylated 17-HGL (Figure 4). Lyciumoside I was identified by retention time and MS/MS fragmentation to a purified authentic standard and the novel compounds were annotated based comparisons on their MS/MS fragmentation to an in-house database of HGL-DTG spectra. Thus, NaUGT74P3 and NoUGT74P4 function as UDP-glucosyltransferases for HGL-DTGs synthesis via their ability to catalyze glucosylation at C-3 and C-17 of the 17-HGL aglycone. When using UDP-glucose as donor and 17-HGL as substrate, NaUGT74P5 and NoUGT74P6 did not show any detectable product in the enzymatic assays. Furthermore, the combination of NaUGT74P5 with NaUGT74P3 or NoUGT74P4 did not result in any further detectable products.

**Figure 4:**
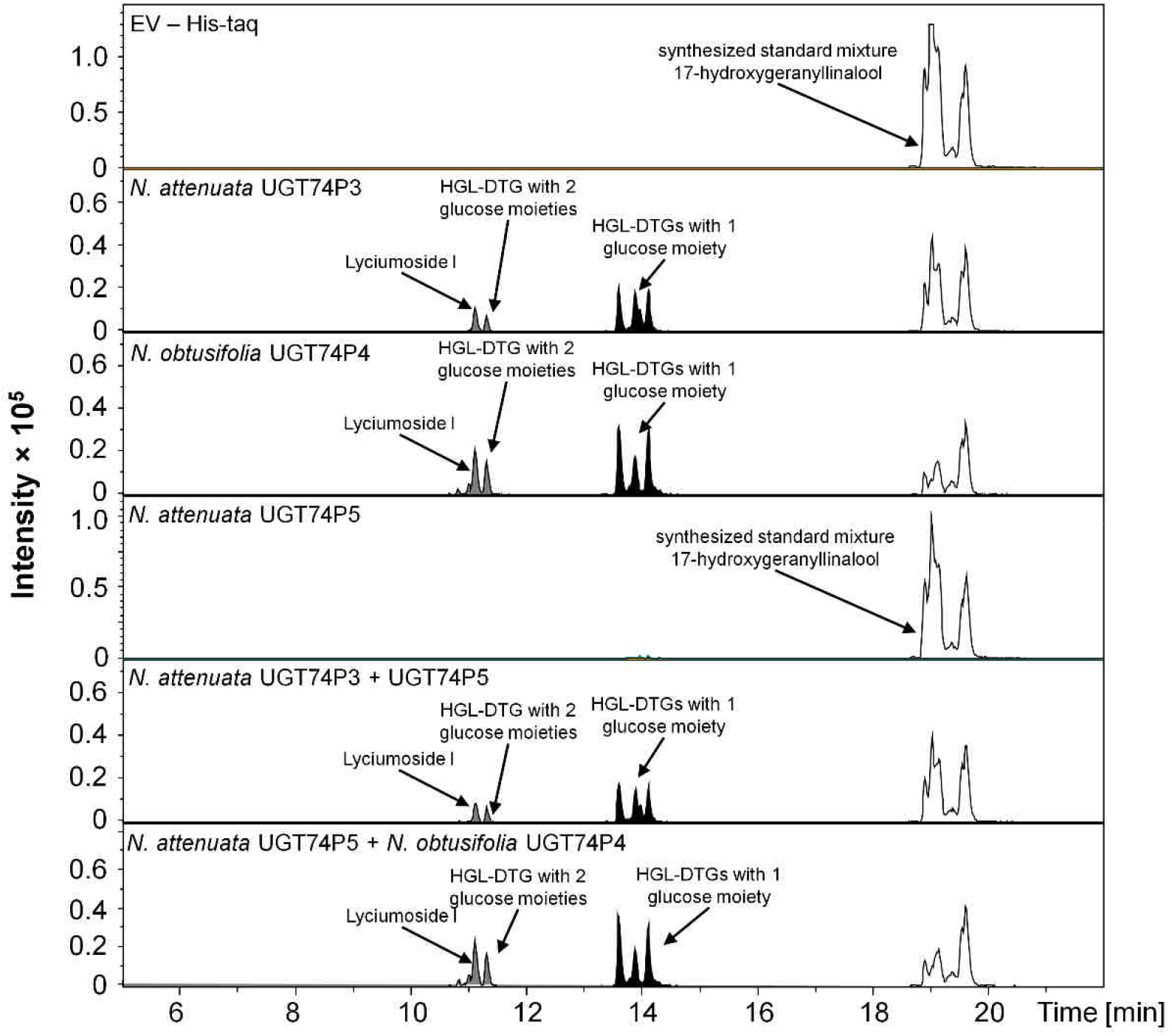
Recombinant UGT74P3, UGT74P4, and UGT74P5 proteins glucosylate the 17-HGL aglycone. Enzyme activity assays of the recombinant UGT74P3, UGT74P4 and UGT74P5 proteins expressed in *E. coli* BL21 DE3 cells. Chromatograms (EIC traces for aglycone m/z 329.2475, HGL-DTGs with 1 glucose moiety m/z 491.3003 and HGL-DTGs with 2 glucose moieties m/z 653.3507) of analyses of 4 UGTs recombinant protein incubated for 3 h in 50 mM Tris HCL pH 7.0 with 5 mM 17-HGL in the presence of 5 mM UDP-glucose. Additionally, activity assays combining UGT74P5 with UGT74P3 or UGT74P4 were performed, but did not differ from the enzyme assays using only UGT74P3 or UGT74P4.

### Virus-induced gene-silencing of UGTs causes morphological defects and necrosis

In addition to the mild TRV infection symptoms such as curly leaves and local chlorosis that were also seen in EV (pTV00) and pTVUGT91T1 plants (Supplemental Figure 13a), we noticed that VIGS plants expressing the pTVUGT74P3 and pTVUGT74P5 constructs displayed severe morphological alterations that ranged from the presence of necrotic spots to necrotic apical meristems and a high percentage of stalled flower buds (Figure 2C, Supplemental Figures 13b and c). Furthermore, we isolated organs of mature flowers collected at anthesis (Figure 2D) and observed significant reductions, compared to observations for pTV00 controls, in the lengths of styles and corolla tubes, and in the width of the corolla limb of pTVUGT74P3 and pTVUGT74P5 VIGS plants, as well as an increased combined lengths of ovary + nectary (Figure 2E).

*N. obtusifolia* plants showed the same symptoms. Plants transiently silenced with pTVNoUGT91T1-like did not display any additional phenotypic alterations compared with the pTV00 control plants. However, *N. obtusifolia* plants silenced in *UGT74P4* and *UGT74P6* expression showed similar morphological modifications as observed in *N. attenuata*, ranging from necrotic spots, misshaped and deformed leaves to stalled flower buds (Figure 3C, Supplemental Figure 14) with the exception of the apical meristem necrosis. Floral morphology was not examined in *N. obtusifolia*.

Finally, we generated a VIGS construct (pTVGLS) targeting the *geranyllinalool synthase* Na*GLS* which was also identified from the co-expression analysis. The encoded protein is predicted to be part of the HGL-DTG biosynthetic pathway by hydroxylating geranyllinalool to produce 17-HGL (Falara et al., 2014). 14 days after inoculation of *A. tumefaciens* with pTVGLS, we detected a 97.5% reduction of the *GLS* transcript abundance compared to pTV00 controls, which was associated with a strong reduction in almost all HGL-DTG types (Supplemental Figure 15 and Supplemental Data 4). No novel intermediate HGL-DTGs were detected, consistent with the expected absence of the precursor molecule, geranyllinalool. Beyond the typical TRV infection symptoms, no morphological alterations were detected, suggesting that the phenotype of the glycosyltransferase-impaired lines resulted from accumulations of intermediate HGL-DTGs or the aglycone 17-HGL.

### Plants stably-silenced for *UGT74P3* and *UGT74P5* display severe developmental defects

To further elucidate the role of UGT74P3 and UGT74P5 in controlling the flux of HGL-DTG synthesis in *N. attenuata* as well as the mechanisms responsible for the developmental defects detected during transient gene silencing, we generated 36 independent stably-transformed transgenic *N. attenuata* plants harboring *UGT74P3* and *UGT74P5* inverted-repeat (IR) silencing constructs in their genomes. Similar to the VIGS experiments, IR*ugt74p3* and IR*ugt74p5*-silenced plants displayed strong effects on growth and development. Almost all stable T_0_-transformants developed either a dwarfish growth or a “broom-like” appearance (Figure 5D). Compared to wild-type (WT) plants, many of the transformed plants were strongly retarded in their growth, with thicker woody, or possibly suberized side branches and deformed or thick, succulent-like leaves. Most flower buds of these transformed plants were small and aborted before fertilization. Few T_0_-transformants of both constructs exhibited a milder phenotype and produced mature flowers and seed capsules. The phenotypic characteristics of IR*ugt74p3* transformants could not be transferred to the T_2_ generation, as T_1_ plants aborted most flower buds early during development and did not produce fertile flowers. IR*ugt74p5* transformants could be propagated and two lines from the same T_1_ parental plant were established (Supplemental Figure 16). Similar to the T_0_ transformants, both IR*ugt74p5* lines displayed a milder phenotype compared to IR*ugt74p3*, but still with severe morphological alterations ranging from necrotic spots and necrotic apical meristem to small deformed or thicker succulent leaves (Supplemental Figure 17a-c) and numerous stalled flower buds (Figure 5C). IR*ugt74p5* plants were also smaller compared to WT (Supplemental Figure 18). Additionally, we established a heterozygous double construct *UGT74P3/UGT74P5* for which ~75% of the silenced plants exhibited similar morphological alterations. Similar to the IR*ugt74p3* plants, the homozygous double construct was lethal and did not produce seeds.

**Figure 5:**
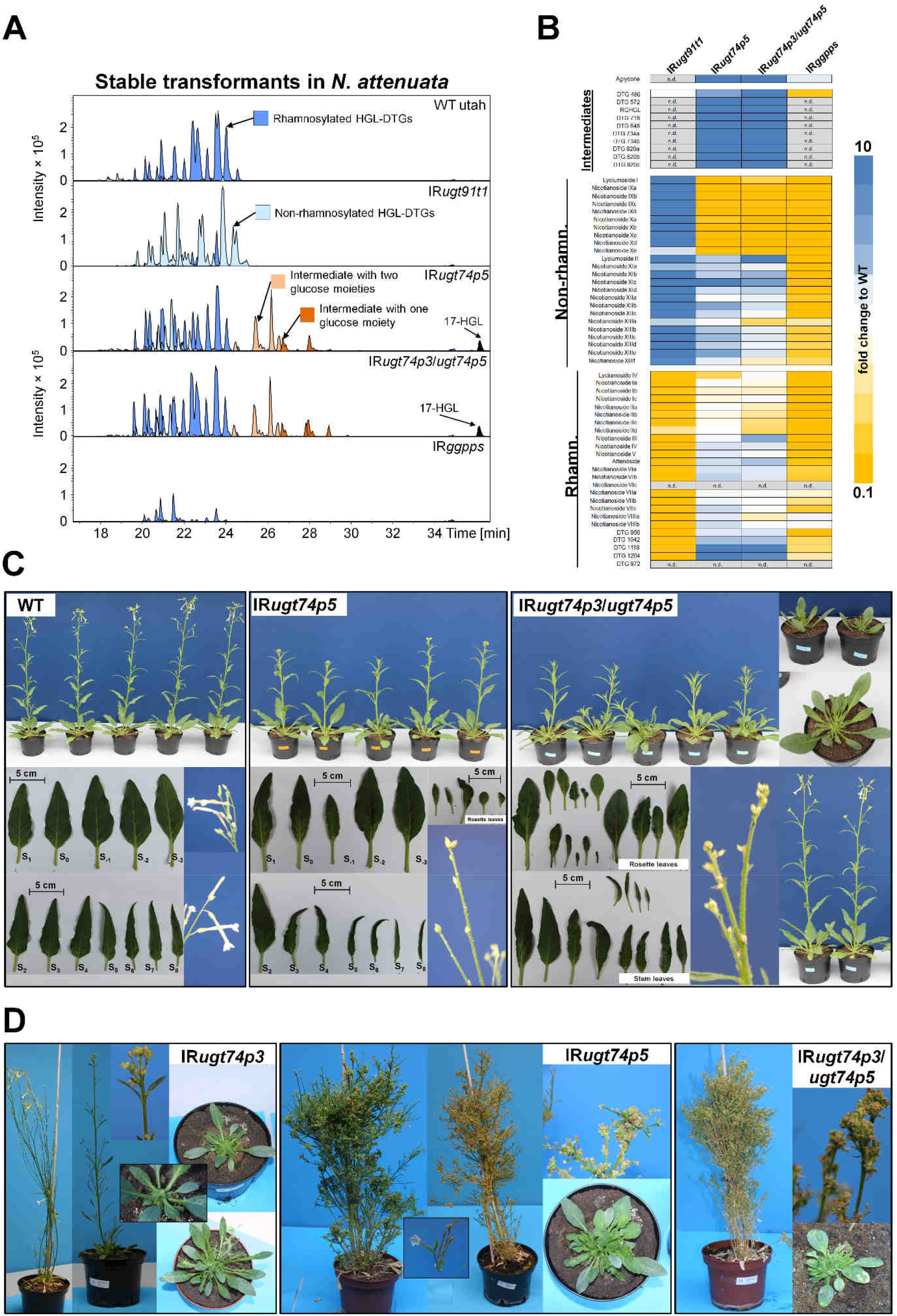
Metabolite profiling and morphological characterization of stably silenced *N. attenuata* plants. A) EICs for identified HGL-DTGs in leaves of 42 day-old elongated *N. attenuata* plants silenced in *GGPPS*, *UGT91T1*, *UGT74P3* and *UGT74P5* transcript accumulation as well as WT control plants. HGL-DTGs were categorized into rhamnosylated, non-rhamnosylated HGL-DTG and intermediates with one or two glucose moieties to facilitate visualization. The 17-HGL aglycone was only detected in IR*ugt74p5* and IR*ugt74p3/ugt74p5* plants. B) Heatmap visualization of deregulations in the leaf HGL-DTG profile of IR*ugt91t1*, IR*ugt74p5*, IR*ugt74p3/ugt74p5* and IR*ggpps* (N=5). The color gradient visualizes fold changes in individual HGL-DTGs for each of the stably-transformed lines compared to the average in the WT plants. C) Morphological alterations in IR*ugt74p5*, IR*ugt74p3/ugt74p5* with milder phenotypes ranged from necrotic spots and tissues, altered leaf shape and thickness, to apical meristem necrosis and a high percentage of stalled flower buds and overall highly stunted growth. Additional details of these phenotypes are shown in supplemental Figure S14a-c. D) 1-year-old independent T0-transformants silenced in the expression of *UGT74P3*, *UGT74P5* and *UGT74P3/UGT74P5*. Strong morphological alterations ranging from stunted growth, succulent leaves, stalled flower buds to a ‘broom’ like appearance were consistently detected among T0-transformants. Viable seeds were produced by a few transformants.

In contrast to the IR*ugt74p3* and IR*ugt74p5*-silenced plants, two independent stably-silenced *UGT91T1* lines did not show any morphological alteration compared to WT (Supplemental Figure 19).

In addition to the careful examination of shoot morphological alterations (roots were not examined), we analyzed transcript abundance of *UGT74P3*, *UGT74P5* and *UGT91T1* in leaves of 52-day-old plants of the different transgenic lines (Figure 6). The silencing efficiency for *UGT91T1* transcript abundance was 97.4% for IR*ugt91t1a* and 97.6% for IR*ugt91t1b*. *UGT74P5* transcript abundance was strongly repressed in all IR*ugt74p5* lines tested (83.9% in IR*ugt74p5b*, 96.8% in IR*ugt74p5a* and 92.3% in IR*ugt74p3/ugt74p5)*. The silencing efficiency for *UGT74P3* in IR*ugt74p3/ugt74p5* was 99.3%, but we also detected a strong co-silencing of *UGT74P3* expression in IR*ugt74p5b* (96.7%) and IR*ugt74p5a* (99.4%).

**Figure 6:**
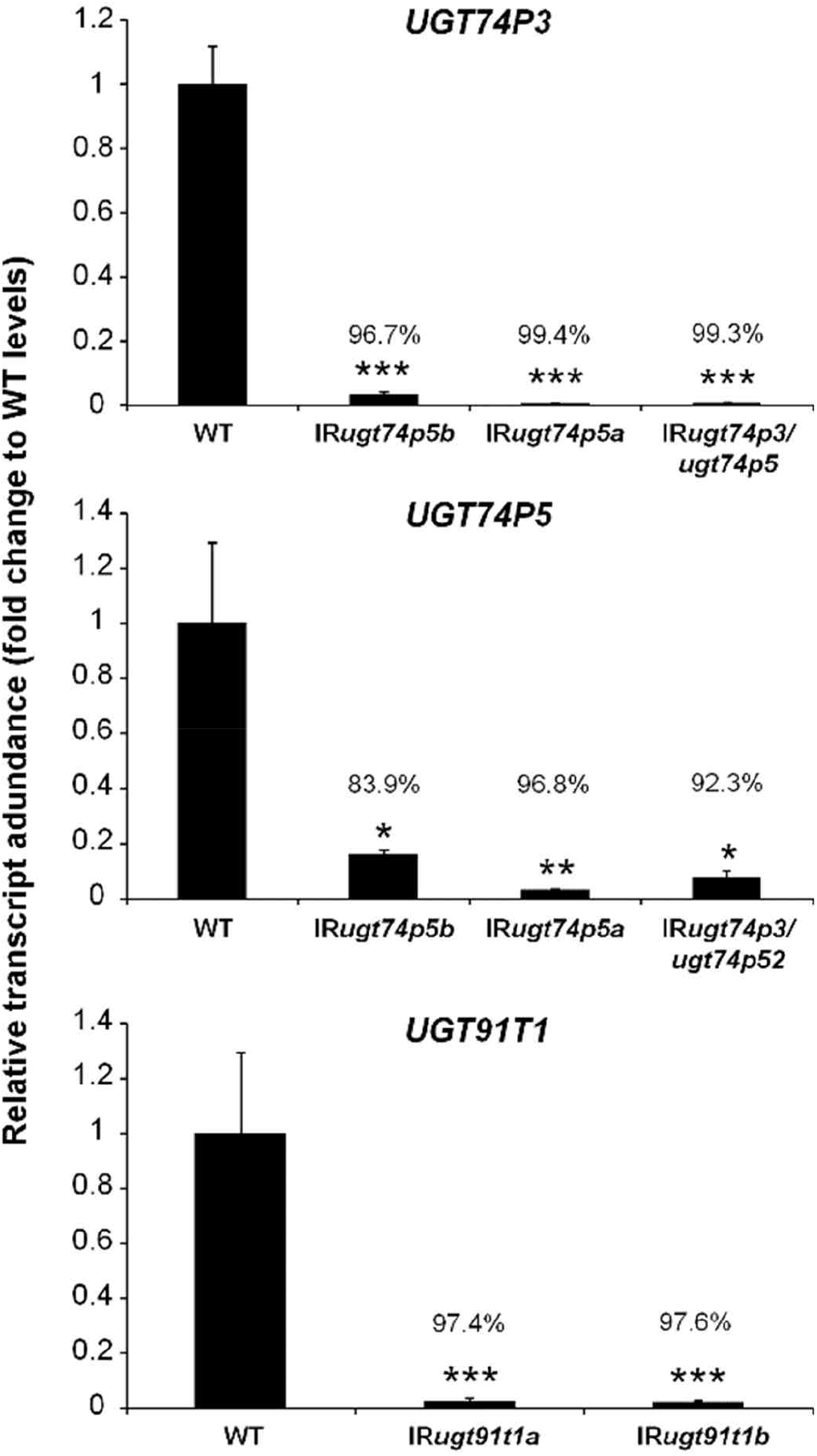
Silencing efficiency for the three 17-HGL-DTG biosynthetic UGTs in IR *ugt91t1*, IR *ugt74p5*, IR *ugt74p3/ugt74p5* plants. Relative transcript abundance of *UGT91T1*, *UGT74P3* and *UGT74P5* in leaves of stably transformed *N. attenuata* plants (N=4). Asterisks indicate significant differences between WT control and stable transformants (*P ≤ 0.05, ** P < 0.01, *** P < 0.001).

### Metabolic profiling confirms the unique HGL-DTG profiles of stably-silenced UGT lines

From UPLC/qTOF-MS measurements and the application of our de-replication workflow, we identified 60 HGL-DTGs in the leaf extracts of the different stably-silenced lines. HGL-DTGs were annotated and classified, based on their chemical composition, into three categories as rhamnosylated, non-rhamnosylated and biosynthetic intermediates only detectable upon disruption of glucosylation steps (Figure 5). Both stable lines impaired in *UGT74P5* expression showed the highest overall HGL-DTG levels (Supplemental Figure 20). Non-rhamnosylated lyciumoside I and its malonylated forms, nicotianoside IXa-d and Xa-e, were barely detectable and levels of rhamnosylated lyciumoside IV was strongly reduced. While rhamnosylated nicotianoside I and II remained constant compared to WT plants, both transgenic IR*ugt74p5* lines exhibited elevated levels of lyciumoside II and attenoside and their malonylated forms, nicotianoside XIa-d, XIIa-c, XIIIa-f and nicotianoside VIa-c, VIIa-c and VIIIa-b, respectively. More complex reconfiguration patterns were also detected such as for nicotianoside III which remained at unchanged levels in transgenic IR*ugt74p5 lines* but whose malonylated forms, nicotianoside IV and V, increased in concentrations. Furthermore, we putatively identified 10 novel highly abundant intermediate HGL-DTGs and the 17-HGL aglycone which were not detected in WT plants. Annotation of the MS/MS spectra of these novel HGL-DTGs indicated lower molecular weight HGL-DTG biosynthetic intermediates with either one (G-3-HGL, G-17-HGL) or two (RGHGL and DTG 648) sugar moieties attached to the 17-HGL (Figure 5, Supplemental Figures 11a-c, Supplemental Table 7). Additionally, malonylated forms of these compounds were also detected (DTG 572, DTG 718, DTG 734 and DTG 820) (Figure 5A and B, Supplemental Figures 11a-c, Supplemental Table 7). The transgenic line IR*ugt74p5b* was used for all further experiments. The heterozygous double construct IR*ugt74p3/ugt74p5* exhibited an almost identical pattern of accumulation of known and novel intermediate HGL-DTGs. The most prominent difference compared to the IR*ugt74p5* lines was the specific increase detected for lyciumoside II and lyciumoside IV.

In both stable lines impaired in *UGT91T1* expression, all rhamnosylated HGL-DTGs were strongly decreased. Reciprocally, all non-rhamnosylated HGL-DTGs were highly increased leading to a complete shift between rhamnosylated and non-rhamnosylated compounds within the HGL-DTG profile (Figure 5A and B). Neither intermediate HGL-DTGs nor the 17-HGL aglycone could be detected in either of the transgenic IR*ugt91t1* lines. IR*ugt91t1* line A was selected for all further experiments. Additionally we analyzed the HGL-DTG profiles in IR*ggpps* plants (Figure 5). Geranylgeranyl-pyrophosphate synthase (GGPPS) is responsible for the synthesis of GGPP, the precursor for all HGL-DTGs (Jassbi et al., 2008) and almost all HGL-DTGs were decreased and no intermediates could be detected. Only levels of nicotianoside VIIIa and VIIIb were increased. Surprisingly, traces of the 17-HGL aglycone were detected. A more detailed analysis of the HGL-DTG profiles in all stable silenced lines can be found in Supplemental Data 5.

In addition, we performed a detailed metabolite profiling of central carbon metabolism intermediates and specialized metabolites, which is further described in Supplemental Method File 2. This included quantitative profiles of 23 amino acids and biogenic amines, small organic acids, phenylpropanoids and derivatives, sugars and phytohormones, such as gibberellins, cytokinins and jasmonates (Supplemental Figures 21 and 22, Supplemental Data 6 and 7).

### Abolishing 17-HGL aglycone synthesis in IR *ggpps* prevents the strong morphological alterations that result from silencing *UGT74P3* and *UGT74P5*

To determine if the morphological alterations of plants transiently- and stably-silenced in *UGT74P3* and *UGT74P5,* are mediated by altered HGL-DTG profiles, we performed a VIGS experiment silencing *UGT74P3* and *UGT74P5* in *N. attenuata* plants impaired in *GGPPS* expression (IR*ggpps*) and WT plants. Quantification of 17-HGL concentrations in leaf tissues using a quantitative U(H)PLC-triple quadrupole-MS method (Supplemental Figure 23) showed accumulation of 17-HGL only in WT *N. attenuata* plants transformed with the pTVUGT74P3 or pTVUGT74P5 VIGS construct (Supplemental Figure 24). 17 days after inoculation, we analyzed the morphological alterations triggered by *UGT74P3* and *UGT74P5* silencing (Figure 7). WT plants transiently transformed with pTVUGT74P3 and pTVUGT74P5 exhibited the phenotype described above, ranging from necrotic spots and necrotic apical meristems to stalled and aborted flower buds (Figure 7A, Supplemental Figure 25, 26 and 27). The number of side branches increased and the rosette diameter decreased. IR*ggpps* plants transformed with pTVUGT74P3 and pTVUGT74P5 did not show any severe morphological alterations. Specifically, the number of side branches and rosette diameters of these plants were not different from those of WT plants (Figure 7B). This combination of genetic manipulations indicates that the ectopic accumulation of several intermediates and of 17-HGL might be directly responsible for the observed morphological alterations.

**Figure 7:**
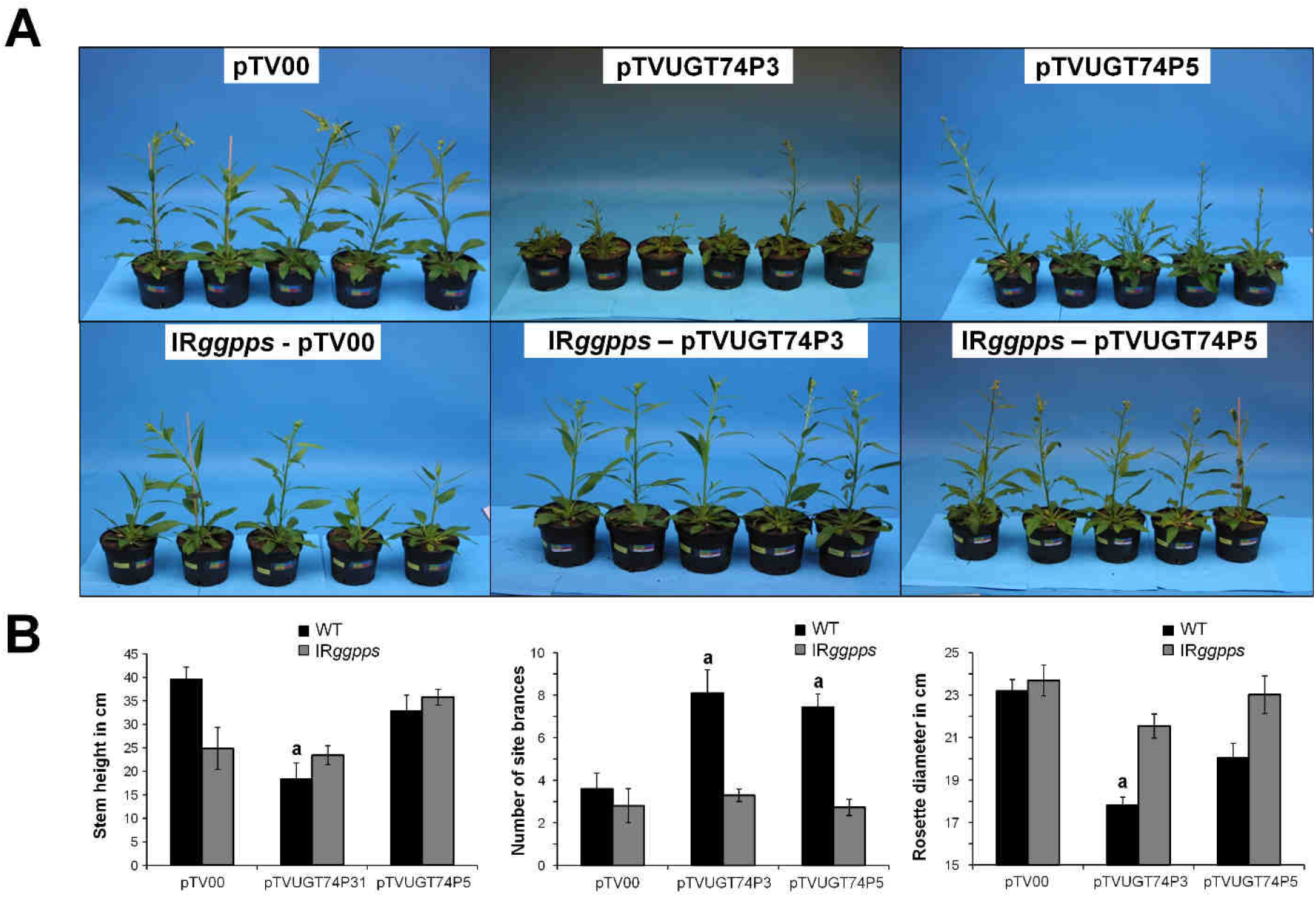
Abolishing 17-HGL aglycone synthesis by silencing Na*GGPPS* abrogates morphological alterations resulting from the silencing of *UGT74P3* and *UGT74P5*. A) Morphological alterations of 42-day-old *N. attenuata* plants stably transformed to silence Na*GGPPS* expression (IR*ggpps*) and WT after inoculation with *A. tumefaciens* harboring pTVUGT74P5, pTVUGT74P3 and the empty vector control (pTV00) VIGS constructs. B) Stem height, number of side branches and rosette diameter in WT and IR*ggpps*. Asterisks indicate significant differences between empty vector control and transient silenced lines (*P ≤ 0.05, ** P < 0.01, *** P < 0.001).

### Excess of 17-HGL triggers cell necrosis in *N. attenuata* leaves

We determined the absolute concentration of 17-HGL in leaves of *N. attenuata* plants (N=5) impaired in *UGT74P5* or *UGT74P3/UGT74P5* expression which exhibited the severe phenotype reported above. The concentration (mean ± SD) of 17-HGL was 94 ± 8 nmol/g FW in IR*ugt74p5* and 85.4 ± 17.6 nmol/g FW in IR*ugt74p4/ugt74p5* (Figure 8A). For the determination of the phytotoxic effect of 17-HGL, we used 32-day-old early-elongated *N. attenuata* (N=3) WT plants and 48-day-old flowering *N. attenuata* plants impaired in *GGPPS* expression (N=5). The average leaf mass was estimated between 1.3 – 1.6 g for the rosette leaves of WT and the stem leaves of transgenic IR*ggpps* plants. Three leaves of each plant were treated with either 20 μL of DMSO or DMSO with 140, 280, or 9800 nmol 17-HGL. Independent of the concentration of 17-HGL, DMSO application resulted in a mild dissolution of the epidermal surface. Analysis of the damaged leaf area, 1 day after application revealed strong necrotic regions at all three 17-HGL concentrations (Figure 8B). Significant increases in leaf damage were detected starting at 140 nmol/g FW for IR*ggpps* plants and 280 nmol/g FW in WT plants (Figure 8C).

**Figure 8:**
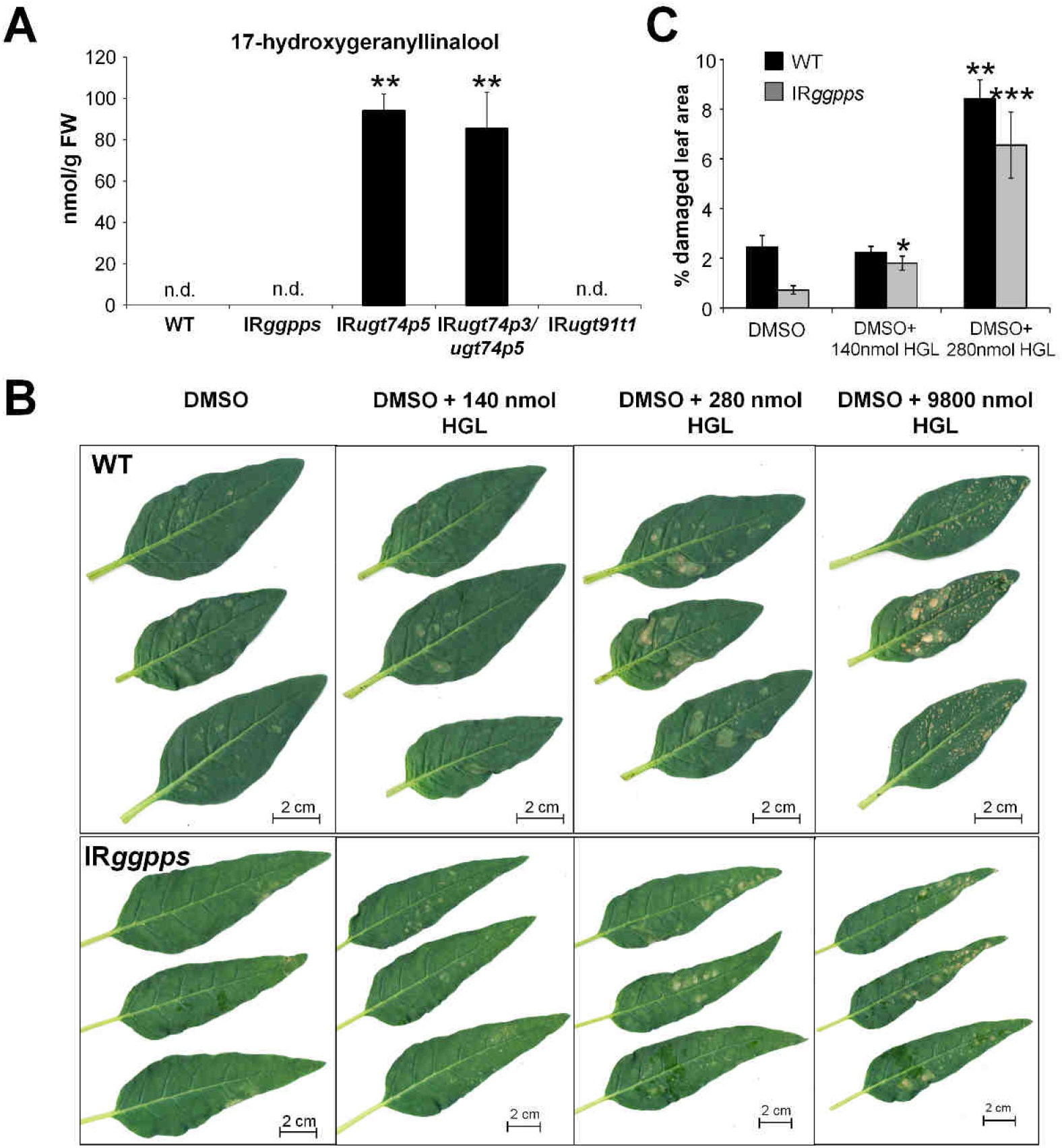
Application of 17-HGL aglycone results in necrotic lesions that phenocopy those observed in IR*ugt74p5* and IR*ugt74p3/ugt74p5* plants. A) Concentrations of HGL in leaf material of WT, IR*ggpps*, IR*ugt74p5*, IR*ugt74p3/ugt74p5* and IR*ugt91t1*. B) Necrotic leaf tissue of 32 day-old elongated WT and 48 day-old flowering IR*ggpps* plants treated with DMSO, DMSO + 140nmol HGL, DMSO + 280 nmol HGL and DMSO + 9800nmol HGL after 1 day. C) Percentage of damaged leaf area in WT (N=3) and IR*ggpps* (N=5). Asterisks indicate significant differences between Control (DMSO) and treated (+HGL) leaves (*P ≤ 0.05, ** P < 0.01, *** P < 0.001).

### *Manduca sexta* larvae perform poorly on transformed lines impaired in HGL-DTG glycosylation

HGL-DTGs are abundant and potent anti-herbivore defense metabolites in the aboveground tissues of *N. attenuata* (Heiling et al., 2010). Although 17-HGL has been suggested as a feeding deterrent (Jassbi et al., 2010), which of the many different HGL-DTGs or which structural component of HGL-DTGs accounts for the observed deterrent (Jassbi et al., 2006) and resistance effects against herbivores (Snook et al., 1997) remains unknown. To elucidate the defensive value of glucosylated and rhamnosylated HGL-DTGs, we conducted performance assays with *M. sexta* larvae reared on leaf disks of transgenic *N. attenuata* plants impaired in rhamnosylation (IR*ugt91t1*) and glucosylation (IR*ugt74p5*, IR*ugt74p3/ugt74p5*) of HGL-DTGs. Additionally, we included IR*ggpps*, which has been shown to dramatically decrease HGL-DTG levels and enhance growth of *M. sexta* (Heiling *et al.* 2010).

Caterpillars feeding on IR*ggpps* plants showed enhanced growth after 6 days and gained 1.6-fold higher larval mass after 12 days (P=0.000) compared to larvae feeding on WT leaf disks (Figure 9A). In contrast, *M. sexta* feeding on leaf disks from IR*ugt74p5* (~0.6 fold, P=0.028) and IR*ugt74p3/ugt74p5* (~0.6-fold, P=0.001) plants showed significantly reduced growth compared to larvae feeding on WT leaf disks. Caterpillars fed leaf disks of IR*ugt91t1* displayed only slight reductions in growth (~0.8 fold relative to WT).

**Figure 9:**
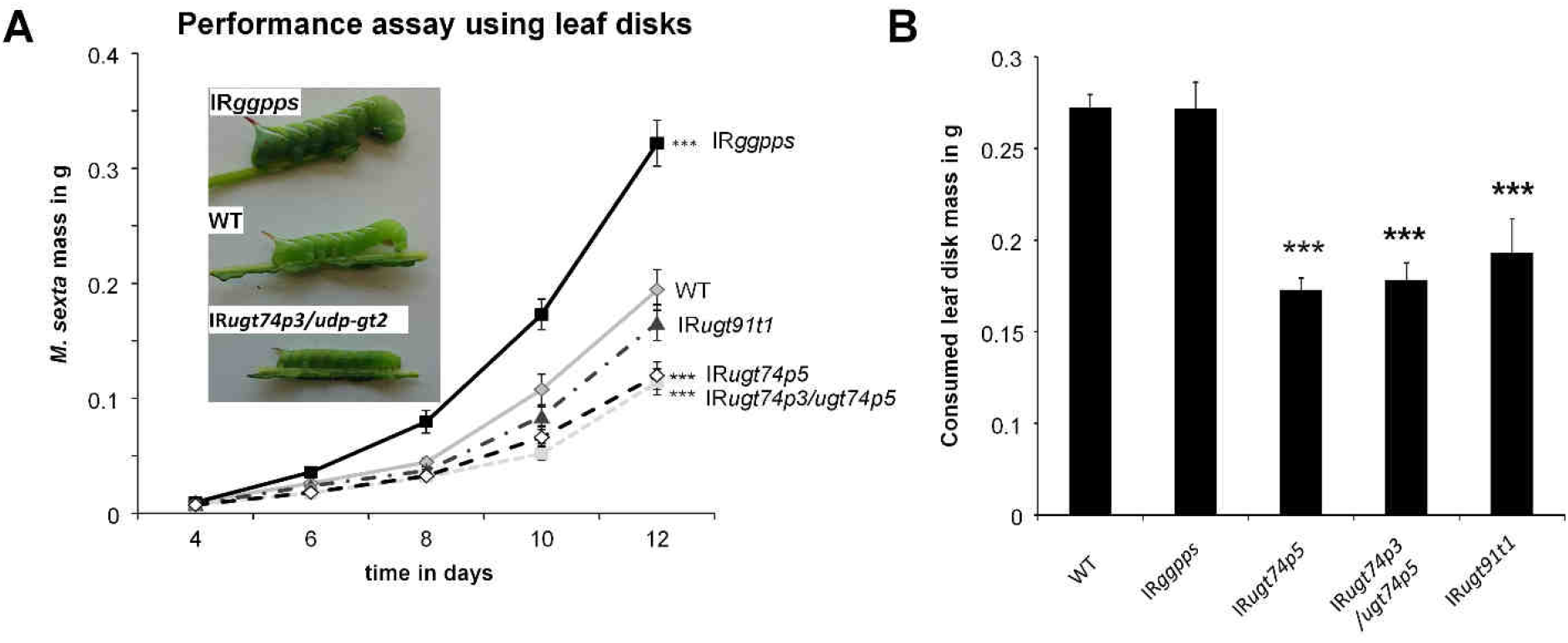
Performance assays show reduced growth of *M. sexta* fed on transformed lines impaired in HGL-DTG glycosylation. A) Mass of *M. sexta* larvae feeding on leaf disk material of four stably transformed plants impaired in glucosylation (IR*ugt74p5*, IR*ugt74p3/ugt74p5*) and rhamnosylation (IR*ugt91t1*) of HGL-DTGs as well as the formation of their precursor geranylgeranyl diphosphate (IR*ggpps*) (average ± SE; *n*=24 to 30). Larvae grow significantly larger on IR*ggpps* (P=0.005) and are significantly smaller on IR*ugt74p3/ugt74p5* (P=0.005) and *IRugt74p5* (P=0.006) by day 6 as determined by Mann-Whitney-Wilcox Pairwise Tests. For clarity, significance is only shown for days 12: *P<0.05, **P<0.01, ***P<0.001. B) Mass of consumed leaf disks after caterpillars fed leaf disks of transgenic plants (average ± SE; *n*=26 to 30 leaf disks with one larva). Larvae fed IR*ugt74p5*, IR*ugt74p3/ugt74p5* and IR*ugt91t1* consumed significantly less leaf disk material as determined by Mann-Whitney-Wilcox Pairwise Tests. *P<0.05, **P<0.01, ***P<0.001.

Additionally, we measured the mass of leaf tissue consumed by *M. sexta* larvae fed IR*ggpps*, IR*ugt74p5*, IR*ugt74p3/ugt74p5* and IR*ugt91t1* leaf disks between days 8 to 10 and observed a strong decrease in consumption of IR*ugt74p5*, IR*ugt74p3/ugt74p5* and IR*ugt91t1* leaf disks relative to WT (Figure 9B). A detailed statistical analysis can be found in Supplemental Data 8.

## Discussion

Plants have evolved highly diversified specialized metabolism pathways to resist both abiotic and biotic stresses as well as a series of mechanisms to mitigate cost/benefit trade-offs of specialized metabolite production and thereby maintain competitive ability. Innovations in specialized metabolism frequently result from the modification or direct recruitment of pre-existing scaffolds produced by core metabolic pathways. These scaffolds serve as substrates for modifying enzymes that add many different types of decorations (Gachon et al., 2005). In this respect, the large numbers of UGTs that populate plant genomes define a versatile glycosylation toolbox that has likely facilitated the functional diversification of secondary metabolism across plant lineages.

Here we identified and phylogenetically characterized 107 UDP-glycosyltransferases of the superfamily 1 in *N. attenuata*. We identified three novel UGTs, based on co-expression analysis, which are responsible for the synthesis of 17-HGL-DTGs in *N. attenuata*. UGT74P3 and UGT74P4 are GT-type enzymes that use UDP-glucose to attach the first glucose moieties to the C-3 and C-17 hydroxyl groups of 17-HGL. Additionally, we show that the glucosylation of the 17-HGL aglycone is crucial to prevent an autotoxic effect of 17-HGL accumulation, which causes severe necrosis of the leaves. Collectively, this functional study provides novel insights into the biosynthesis of broad-spectrum anti-herbivore HGL-DTGs and provides evidence consistent with the hypothesis that UGTs play a central role in a plant’s ability to manage the “toxic waste dump” problem of chemical defense.

### A geranyllinalool synthase (GLS) is required for HGL-DTG production

Geranyllinalool (GL) is an acyclic diterpene alcohol which is widely distributed across the plant kingdom, occurring in several essential oils (Sandeep and Paarakh, 2009) and is a precursor for both HGL-DTGs in solanaceous species and for the volatile C_16_-homoterpene 4,8,12-trimethyltrideca-1,2,7,11-tetraene (TMTT) which is emitted from the foliage of a wide range of plant species including *Solanum lycopersicum* (Ament et al., 2004), *Zea mays* (Hopke et al., 1994), *Medicago truncatula* (Leitner et al., 2010) and *A. thaliana* (Van Poecke et al., 2001). Although geranyllinalool is present in many plant species, the enzymes responsible for its biosynthesis have been discovered only recently in *A. thaliana* (Herde et al., 2008), *S. lycopersicum*, and *N. attenuata* in which *GLS* is constitutively expressed in leaf and flower tissues and is induced by methyl jasmonate treatment (Falara et al., 2014). Based on the tissue-specific correlation of *GLS* transcript abundance with the levels of HGL-DTGs in reproductive organs, the authors predicted that NaGLS is involved in the biosynthesis of HGL-DTGs (Falara et al., 2014). Here we provide additional evidence demonstrating its coordinated expression with other HGL-DTG biosynthetic genes (*GGPPS*, *UGT91T1* and *UGT74P3*) and demonstrate that HGL-DTG levels are highly reduced after the transient silencing of Na*GLS*.

### Gene-to-gene co-expression analysis identifies three novel UGTs responsible for the glycosylation of HGL-DTGs

To identify the UGTs responsible for the glycosylation of HGL-DTGs, we mined the expression profiles of 76 UGTs in a full-transcriptome datasets created for leaf and roots collected from plants after simulated herbivory. More specifically, we explored co-linearity patterns in the regulation of this set of UGTs and *GGPPS* and *GLS* which are involved in upstream steps of the biosynthesis of 17-HGL. A previous study by our group (Gulati et al., 2013) detected, using the same transcriptome datasets, a tight co-regulation between genes of the non-mevalonate pathway (Na*DXS,* Na*DXR*) and Na*GGPPS*. Our analysis pinpointed three novel glycosyltransferases strongly correlated with GGPPS and GLS, which were not identified in the previous study. Gulati and colleagues analyzed tissue-specific gene-to-gene and gene-to-metabolite associations using fold-change data in response to herbivory and used very stringent statistical parameters. Here, we used a simpler co-expression approach relying on overall gene-to-gene correlations across all tissues to identify significant correlations in expression dynamics among UGTs and previously characterized upstream genes. Furthermore, a detailed analysis of the expression pattern of each of the 76 UGTs (Supplemental Figure 4) revealed that herbivory has dramatic effects on the local induction of UGT expression, an observation consistent with the importance of glycosylation reactions in maintaining cellular homeostasis after damage.

Together with the three candidate UGTs identified in *N. attenuata*, we also identified and biochemically characterized orthologues in *N. obtusifolia* whose genome was recently published along with that of *N. attenuata* (Xu et al., 2017).

### UDP-Rha: 17-HGL-DTG rhamnosyltransferase activity of UGT91T1

Very little is known about the physiological function of rhamnosylation in secondary metabolism. For example, Hsu and colleagues (Hsu et al., 2017) suggested that rhamnosylation is an essential step in the synthesis of lobelinin and therefore responsible for the color of Lobelia flowers. Rhamnosylation of flavonols is thought to modulate auxin homeostasis in *rol1-2* mutants of *A. thaliana* (Kuhn et al., 2016). However, the enzymatic characterization of UDP-rhamnosyltransferases is still thwarted by the high costs of UDP-rhamnose (UDP-Rha). Despite important efforts in the establishment of efficient UDP-Rha production systems (Irmisch et al., 2018), only a few UDP-rhamnosyltransferases have been functionally characterized and enzymatic assays remain challenging (Mo et al., 2016; Irmisch et al., 2018).

Here we show that UGT91T1 shares a high degree of sequence similarity with functionally characterized rhamnosyltransferases for flavonoids and anthocyanins (CmF7G12RT, GmF3G6R and PhA3ART – Supplemental Figure 6), is tightly and tissue-specifically co-regulated with the accumulation patterns of rhamnosylated HGL-DTGs and exhibits strong co-expression with *GLS*, *GGPPS* and *UGT74P3*. Silencing of *UGT91T1* expression in *N. attenuata* results in the almost complete loss (88.5% - 98.8% in different tissues compared to WT, Supplemental Table 8) of rhamnosylated HGL-DTGs. These changes in HGL-DTG rhamnosylation pattern suggest that UGT91T1 is responsible for the rhamnosylation at the C’-4 hydroxyl-group of glucose on both the C-3 and the C-17 hydroxyl-group of the aglycone. Due to the high biological variability among transiently-silenced plants and the complexity of the HGL-DTG profile in *N. obtusifolia, which* produces lower levels of rhamnosylated HGL-DTGs, the biochemical function of No*UGT91T1-like*, the orthologue of UGT91T1 in *N. obtusifolia*, could not be conclusively evaluated.

### UGT74P3 and UGT74P4 function as UDP-Glu: 17-HGL-DTG glucosyltransferases

Both UDP-glycosyltransferases clustered together in the UGT74 family, which belongs to phylogenetic group L and includes glycosyltransferases that are responsible for the glucosylation of indole-3-acetic acid in *Zea mays* (Szerszen et al., 1994) and *A. thaliana* (Jin et al., 2013). Importantly, additional members of this family show catalytic activity towards diterpene and triterpene glycosides. For example, UGT74M1 glucosylates the carboxylic acid moiety of the triterpene gypsogenic acid (Meesapyodsuk et al., 2007), UGT74-345-2 is involved in the glucosylation of mogroside (Itkin et al., 2016, 2018), and UGT74G1 catalyzes the glucosylation of cyclic diterpene glycosides in *Stevia rebaudiana* (Richman et al., 2005). Both UGT74P3 and UGT74P4 are closely related to UGT74G1 and CsGLT2, a UGT responsible for crocetin glucosylation in *Crocus sativus* (Moraga et al., 2004). The expression profile of *UGT74P3* in *N. attenuata* revealed a strong tissue-specific correlation with HGL-DTG accumulation and *GLS* expression.

Recombinant enzyme activity assays demonstrated that UGT74P3 and UGT74P4 encode enzymes that catalyze the transfer of glucose to the C-3 and C-17 hydroxyl groups of 17-HGL in order to form the putative intermediate products 3-*O*-glucopyranosyl-17-HGL (G-3-HGL) and 17-*O*-glucopyranosyl-17-HGL (G-17-GHL) (Figure 4), as well as lyciumoside I in which both hydroxyl groups are glucosylated. Neither UGT74P3 nor UGT74P4 showed activity producing lyciumoside II, which includes an additional glucose moiety linked to the C’-2 hydroxyl group of the glucose at the C-17 hydroxyl group of 17-HGL, or other larger HGL-DTGs with additional sugar moieties. However, the functional characterization of enzymes in the UGT family can be very challenging and *in vitro* enzyme assays do not necessarily reflect native metabolic activities *in planta*.

Metabolic profiling of transiently- and stably-silenced lines provided further evidence that UGT74P3 and UGT74P4 are essential for the formation of lyciumoside I through the attachment of two glucose moieties. Silencing of *UGT74P3* and *UGT74P4* expression significantly reduces the amount of lyciumoside I and its malonylated forms in leaf tissue material. Interestingly, compounds further downstream in the HGL-DTG biosynthetic pathway (lyciumoside II, lyciumoside IV, attenoside) are more affected in the transiently than in the stably–silenced lines, which might be due to the infection with the virus vector and a possible involvement of HGL-DTGs in the immune response of plants against viruses (Ramegowda et al., 2014). In the stable lines impaired in *UGT74P3* expression, we observed a slight reduction of lyciumoside IV, but an increase of attenoside and lyciumoside II, indicating that the downstream model of the HGL-DTG biosynthetic pathway is incomplete and that other UGTs are involved in the glycosylation of larger molecular weight HGL-DTGs.

In addition to the changes in known HGL-DTG patterns, we also observed an accumulation of novel intermediate HGL-DTGs that lacked either glucosylation at the C-3 or C-17 hydroxyl group of the aglycone as well as the 17-HGL aglycone itself in both stably- and transiently-silenced lines. This suggests that *N. attenuata* is unable to reroute the excess of 17-HGL and provides us with the opportunity to study the effects of glucosylation of HGL-DTGs *in vivo*.

In contrast to UGT74P3 and UGT74P4, we were not able to decipher the function of UGt74P5 in *N. attenuata* and of its orthologue in *N. obtusifolia*. The high sequence similarity between UGT74P3 and UGT747P5 of ~83%, indicated a related function as a UGT. Furthermore, *UGT74P5* expression highly correlates with that of all other genes related to HGL-DTG biosynthesis (*UGT74P3*, *UGT91T1*, *GLS* and *GGPPS*), exhibits a similar temporal dynamic after simulated insect herbivory and shows very low transcript abundance in roots. UGT74P5 showed no activity towards the 17-HGL aglycone, lyciumoside I or the putative intermediates G-3-HGL and G-17-HGL, indicating that UGT74P5 might not directly contribute to the biosynthesis of HGL-DTGs or be responsible for the glucosylation of HGL-DTGs that were not tested in these enzyme assays.

The co-silencing of *UGT74P5* and *UGT74P3* expression in the stable and transiently silenced lines (Figure 6, Supplemental Figure 8), likely due to the high sequence similarity of both UGTs, did not allow us to further characterize the specific function of UGT74P5. The metabolic alterations of these lines likely reflect the silencing of both genes. Based on the enzymatic assays, we inferred that the reduction of lyciumoside I and the accumulation of G-3-HGL, G-17-HGL and 17-HGL directly result from the silencing of *UGT74P3*, leaving it unclear whether UGT74P5 is involved in the synthesis of HGL-DTGs, is non-functional or has a still-unknown enzymatic function beyond the diterpene metabolism.

### Phytotoxicity observed in *UGT74P3*/*UGT74P5* silenced lines

In addition to the striking shifts in HGL-DTG metabolism detected in the stably- and transiently-silenced lines impaired in *UGT74P3* and *UGT74P5* expression, we also observed strong morphological alterations that ranged from small deformed or thicker succulent leaves to numerous stalled flower buds as well as necrotic spots, stunted growth and apical meristem necrosis.

The loss of glycosylation results in the ectopic accumulation of hydrophobic aglycones, which are known to be responsible for a wide range of morphological alterations and growth retardation effects. Some of the best examples, come from the biosynthesis of steroidal alkaloids. Silencing *GAME-1*, a UDP-galactosyltransferase responsible for the glycosylation of steroidal alkaloids, results in severe developmental defects due to the altered sterol composition in membranes triggered by the accumulation of tomatidine (Itkin et al., 2011). Moreover, alterations of the glycosylation of the saponin, hederagenin, severely affected growth in *M. truncatula* (Naoumkina et al., 2010). Loss of function mutants, *sad3* and *sad4,* in *Avena sativa* accumulate the intermediate monodeglucosylated diterpeneglycoside, avenacin A-1, which disrupts membrane trafficking and results in epidermal degeneration and reduced root hair formation (Mylona et al., 2008). Knocking out *UGT74B1*, a UGT responsible for glucosinolate biosynthesis in *A. thaliana,* leads to the accumulation of toxic levels of thiohydroximates, increased auxin levels in seedlings and results in a chlorotic phenotype (Grubb et al., 2004).

The severe growth defects, stalled flower buds and necrosis we observed in *UGT74P3/UGT74P5*-silenced plants may have several explanations. First blocking HGL-DTG glycosylation might increase the endogenous pool of UDP-glucose, resulting in a preferential synthesis of other glycosides or the accumulation of other compound classes (Naoumkina et al., 2010). This might influence hormone homeostasis and therefore contribute to the severe alterations (Grubb et al., 2004; Kuhn et al., 2016). Auxin levels, which have been suggested to mediate some of these observed developmental alterations, are only changed in *GGPPS*-silenced plants but not in the UGT-silenced lines showing the morphological defects. Alterations of the UDP-glucose pool can also influence UDP-glucuronic acid (UDP-GlcA) biosynthesis. Reduction of UDP-GlcA in *ugd2* and *ugd3* mutants of *A. thaliana* leads to swollen cell walls and developmental defects associated with changes in the pectic network (Reboul et al., 2011). IR*ugt74p5* and IR*ugt74p3/ugt74p5* plants exhibit lower glucuronic acid levels (Supplemental Data 6), suggesting that UDP-GlcA production is downregulated, which could mediate the observed smaller leaves and the dwarfish growth phenotype. similar reduction in glucuronic acid levels in *GGPPS*-silenced plants, which do not show the altered growth effects. Alternatively, *UGT74P3/UGT74P5*-silenced plants also exhibited a dramatic increase in levels of intermediate HGL-DTGs and the aglycone 17-HGL compared to other transgenic lines that did not show any morphological phenotypes (Figure 8A). From these results we infer that the accumulations of these intermediates may be toxic for the plant.

When *UGT74P3* and *UGT74P5* were silenced in the background of stably-silenced IR*ggpps* lines, which do not produce the 17-HGL precursor, no developmental abnormalities were detected, indicating that the developmental abnormalities are associated with the altered HGL-DTG metabolism and not due to unexpected off-target effects (Senthil-Kumar and Mysore, 2011). Applications of a synthetic mixture of 17-HGL isomers to leaves of *N. attenuata* resulted in necrotic lesions that pheno-copied those observed in IR*ugt74p5* and IR*ugt74p3/ugt74p5* plants. IR*ggpps* plants, which are strongly down-regulated in the expression of HGL-DTG biosynthetic genes and HGL-DTG accumulation, were more susceptible to the toxic effects of exogenous applications of 17-HGL than WT plants and showed necrotic lesions in response to 17-HGL concentrations lower than those observed in IR*ugt74p3/ugt74p5* plants. The lower toxic effects observed in WT leaves may be due to the chemical sequestration of 17-HGL and intermediate HGL-DTGs by active glycosylation with the HGL-DTG pathway. Interestingly, *N. sylvestris* accumulates 17-HGL without producing the glycosylated forms (Heiling et al., 2016), suggesting that other species might be tolerant to or store 17-HGL in special compartments. In short, silencing *UGT74P3* and *UGT74P5* results in pleiotropic morphological affects that are associated with the accumulation of the phytotoxic aglycone 17-HGL.

### Defensive function of HGL-DTGs

HGL-DTGs have been studied for more than two decades and are described to be potent anti-herbivore defense compounds (Heiling et al., 2010) with deterrent (Jassbi et al., 2006) and resistance (Snook et al., 1997) effects against herbivores. However, so far it remains unclear which of the many HGL-DTGs, their structural components or post-ingestive modifications account for their mode of action. The impressive structural diversity of HGL-DTGs in *N. attenuata* plants results largely from glycosylation and malonylation reactions. While all malonyl decorations are instantaneously lost in the alkaline pH environment of the midgut of *M. sexta* larvae and therefor may play more important roles *in planta* (Poreddy et al., 2015; Li et al., 2018), glycosylation leads to a constant and stable pool of potential defensive compounds. Larvae feeding on leaf disks of transgenic IR*ggpps* plants grow larger than larvae feeding on WT leaves, indicating that the overall abundance of HGL-DTGs is a major factor of the plant’s resistance against *M. sexta* (Figure 9), consistent with earlier studies (Jassbi et al., 2008; Heiling et al., 2010; Jassbi et al., 2010). The reduced growth of larvae feeding on leaf disks of transgenic IR*ugt74p5* and IR*ugt74p3/ugt74p5* plants, which have higher total levels of HGL-DTGs, is also consistent with the inference that the total abundance of HGL-DTGs is defensively relevant. For example, *N. obtusifolia* is a perennial plant, which co-occurs with the annual *N. attenuata* in the Great Basin Desert and is much less attacked by the tobacco hornworm *M. sexta*. This species lacks *N. attenuata*’s strong OS-elicited signaling system and hence produces attenuated jasmonate bursts and trypsin proteinase inhibitors (TPI) accumulations (Pearse et al., 2006; Anssour and Baldwin, 2010; Xu et al., 2015). Furthermore, *N. obtusifolia* has constitutive levels of HGL-DTGs that are five times higher than those of *N. attenuata* and *M. sexta* larval performance is five times lower after 22 days feeding on *N. obtusifolia* (Jassbi et al., 2010). However, the HGL-DTG profile in *N. obtusifolia* is structurally different (Heiling et al., 2016), suggesting that the composition or degree of glycosylation is central for the plant’s defense.

Importantly, the specific structures responsible for the mode of action of HGL-DTGs remain unclear. Feeding of purified lyciumoside IV, the most abundant HGL-DTG in the leaves of *N. attenuata*, causes mortality in *M. sexta* larvae silenced in β*-glucosidase 1* (*BG1*) expression (Poreddy et al., 2015). However, the single-compound feeding experiment does not reflect the chemical diversity of HGL-DTGs that is normally consumed by larvae when they feed on plants. In contrast to the results of Poreddy and colleagues, we show that, while consuming less leaf tissue, *M. sexta* larvae grow normally when feeding on leaf disks of transgenic IR*ugt91t1* plants, which have reduced levels of lyciumoside IV, but increased levels of the non-rhamnosylated HGL-DTGs lyciumoside I and II. Furthermore, we observed increased levels of novel intermediate HGL-DTGs and the aglycone, 17-HGL, in IR*ugt74p5* and IR*ugt74p3/ugt74p5* plants, from which we observed a strong antifeedant/deterrent effect, which showed a strong antifeedant/deterrent effect (Figure 9B) and reduced growth performance of *M. sexta* larvae (Figure 9A). The differential consumption and growth performance responses on the transgenic plants with different HGL-DTG profiles suggest that both pre- and post-ingestive resistance mechanisms may be at play. Given the severe metabolic and pleiotropic morphological phenotypes of IR*ugt74p5*, IR*ugt74p3/ugt74p5* and IR*ggpps* plants, it is challenging to draw strong inferences about the defensive function of specific intermediates in the HGL-DTG biosynthetic pathway.

We are still in the early stages of understanding how plants solve the “toxic waste dump” problem. Two common solutions are well documented: 1) producing metabolites (e.g., nicotine) that are specifically toxic to tissues or organs that plants lack (e.g., nervous systems and neuro-muscular junctions populated with nicotinic acetylcholine receptors); and 2) sequestering pro-toxins apart from their toxin-releasing lytic enzymes (e.g., compartmentalization of cyanogenic glycosides and glucosinolates and their active enzymes). However, these examples are likely to be special cases and the solutions that plants have evolved to solve the “toxic waste dump” problem for a majority of their toxic defensive metabolites lie in the details of their biosynthetic pathways. Here we advance our understanding of the toxicity of HGL-DTGs by showing that glucosylation plays a central role in *N. attenuata*’s solution of maintaining an HGL-DTG-based defense. Extending the metabolomics analyses to the “digestive duet” that occurs between plant and insect (“frassomics”) will allow for a better understanding of the post-ingestive fate of DTGs in larval guts and will help to answer a central and festering question that remains unanswered from this work: are the specific molecules that are responsible for the defensive function of HGL-DTGs the same as those responsible for the clear autotoxicity?

## Methods

### Plant material and growth conditions

Seed germination and growth condition have been described previously (Krugel et al., 2002). Seeds of the 31^th^ generation of an inbred line of *N. attenuata* Torr. Ex. Watts were used as wild type plants in all experiments. *N. obtusifolia* plants were cultivated under the same growth conditions with the exception of not applying liquid smoke to the seeds. Plants for virus-induced gene-silencing were transferred after 20 days to a York Chamber with 22°C for a 16h light/ 8h dark cycle.

### *M. sexta* growth conditions

*M. sexta* eggs were obtained from an in-house colony in which insects are reared in a growth chamber (Snijders Scientific, Tilburg, Netherlands, http://www.snijderslabs.com) at 26°C:16-h light and 24°C:8-h dark, 65% relative humidity, until hatching.

### Performance Assays

Freshly hatched neonates were fed leaf disk material, taken from the between-vein laminal tissue of the four lowest stem leaves of 43-day-old flowering transgenic *N. attenuata* plants (WT, IR*ugt91t1*, IR*ugt74p5*, IR*ugt74p2/ugt74p5*, IR*ggpps*), in round plastic PE-packing cups. Leaf disk material was exchanged every 2 days until day 6 and then exchanged every day until day 12.

### Plant transformation and screening of stably-silenced *N. attenuata* plants

Transformation of *N. attenuata* was performed as described in Krugel et al. (2002) using the pRESC8 vector (Gase and Baldwin, 2012) containing the hygromycin phosphotransferase II gene (*hptII*) from pCAMBIA-1301 (GenBank AF234297) and a 306 bp long fragment for *UGT91T1*, a 310 bp long fragment for *UGT74P3* or a 295 bp long fragment for *UGT74P5*. All primers are shown in supplemental table 9. Diploid plants were selected by flow cytometry of leaf material of elongated *N. attenuata* transformants performed on a CCA-II flow cytometer (Partec, http://www.partec.com) as described by Bubner et al. (Bubner et al., 2006). Afterwards, seeds were collected and individuals with the T-DNA insertion were selected for hygromycin resistance by addition of 35 mg/L hygromycin B to the germination medium. After 10 days, the ratio of seedlings surviving the antibiotic treatment was determined. Seedlings were chosen with a survival rate of 50 – 90% and 12 T1 plants per line were checked for T-DNA insertion. To confirm the integrity of the T-DNA insertion, we performed a diagnostic PCR using the primer pairs PROM FOR/INT REV and INT FOR/TER REV (Gase et al., 2011). Genomic DNA (gDNA) was isolated from leaves of *N. attenuata* using a modified cetyltrimethylammonium bromide method (Bubner et al., 2004). PCR was performed using DreamTaq™ DNA-Polymerase (Fermentas, http://www.fermentas.com) according to the instructions of the manufacturer with 1 μg of gDNA. Homozygosity of T2 plants was determined by screening for resistance to hygromycin B. To confirm single insertions, we performed a Southern blot as described by Jassbi et al. (Jassbi et al., 2008), with the exception that a 287 bp *hptII* probe obtained by PCR with primer pair (HYG1-18/HYG2-18) was used (Gase et al., 2011). Labelling was performed with the GE Healthcare (http://www.gehealthcare.com) Readyprime DNA labelling system and ProbeQuant g-50 microcolumn according to the manufacturer’s protocol. 10.5 μg of gDNA was digested with the restriction enzymes EcoRV and XbaI from New England Biolabs (http://www.neb.com) and blotted onto a nylon membrane (GeneScreenPlus; Perkin Elmer, http://www.perkinelmer.com) according to the manufacturer’s instructions.

### Virus-induced gene-silencing (VIGS)

Vector construction, plant growth, and inoculation conditions were as described by Saedler and Baldwin (Saedler and Baldwin, 2004). Briefly, 200-to 300-bp fragments of *N. attenuata* and *N. obtusifolia* target genes were amplified by PCR using primer pairs as listed in supplemental table 9. Amplified fragments were cloned in the vector pTV00 (Ratcliff et al., 2001). *Agrobacterium tumefaciens* strain GV3101 was transformed by electroporation with the resulting plasmids. We used the empty pTV00 vector as a negative control in all experiments. Four leaves of 24-27 day old *N. attenuata* and 25 day old *N. obtusifolia* plants were infiltrated with a 1:1 mixture of *A. tumefaciens* transformed with pBINTRA (Ratcliff et al., 2001) and one pTV00 derivative carrying a fragment of a gene of interest. pTVPDS, targeting for a *Phytoene desaturase,* was used as a positive control to monitor silencing progress. Due to the depletion of carotenoids, silencing the phytoene desaturase gene causes bleaching of tobacco leaves. VIGS-silenced plants were treated 14 days after inoculation, when the bleaching phenotype was fully established in the pTVPDS plants.

### Plant treatment

In order to analyze the regulatory function of jasmonate signaling on HGL-DTG biosynthesis, petioles of five elongated plants (38-days-old) were treated with either 20μL lanolin paste containing 150 μg methyl jasmonate (Lan + MeJA) or with 20 μL pure lanolin (Lan). Treated leaves were harvested from elicited and unelicited plants at 72 h after treatment, flash-frozen in liquid nitrogen, and stored at −80°C until use.

### Determination of 17-HGL phytotoxicity

For the determination of the phytotoxic effect of 17-HGL, we used 32-day-old early-elongated *N. attenuata* WT plants (N=3) and 48-day-old flowering *N. attenuata* plants impaired in *GGPPS* expression (N=5). Three leaves of each plant were inoculated with either 20 μL DMSO, DMSO with 140 nmol HGL, DMSO with 280 nmol HGL or DMSO with 9800 nmol HGL. The damaged leaf tissue was analyzed after 24 h using ImageJ (Fiji, https://fiji.sc).

### RT-qPCR analysis of transcript levels

Total RNA was extracted from an aliquot of approximately 200 mg of powdered leaf material of *N. attenuata* and *N. obtusifolia* ground in liquid nitrogen following the protocol of Kistner and Matamoros (Kistner and Matamoros, 2005). DNase treatment was performed using the TURBO DNA-free™ kit (Invitrogen). RNA quality was checked on a 1% agarose gel and the concentration was determined spectrophotometrically at 260 nm. A total of 1 μg of DNA-free RNA was reverse transcribed using oligo(dT)18 primers and the SuperScript II enzyme (Invitrogen) following the manufacturer’s recommendations. All RT-qPCR assays were performed using Taykon™ No ROX SYBR® Master Mix dttp Blue (Eurogenetics, http://www.eurogentec.com) on a Stratagene MX3005P instrument (http://www.stratagene.com) as recommended by the manufacturer. To normalize transcript levels, primers specific for the *Nicotiana attenuata elongation factor-1*α gene (EF1-α; accession no. GBGF01000210.1) were used. Specific primers in the 5′ to 3′ direction used for SYBR Green-based analyses are listed in supplemental table 9.

### Heterologous expression of UDP-glucosyltransferases and enzymatic activity assays

The four UGT cDNAs coding for NaUGT74P3, NoUGT74P4, NaUGT74P5 and NoUGT74P6 were cloned into a Gateway® pDEST17 expression vector (ThermoFisherScientific, http://www.thermofisher.com) using pET28a empty vector as control. Integrity of the sequence was checked by Sanger-sequencing using an ABI PRISM 3130 Genetic analyzer (AppliedBiosystems, http://www.thermofisher.com) and the appropriate gene-specific primers (Supplemental Table 9).

The resulting plasmid was transformed into BL21 (DE3) *E. coli* which is optimized for the expression of eukaryotic genes. 50 mL LB medium containing 50 μg/mL carbenicillin were inoculated with 500 μL of an overnight culture corresponding to each candidate gene. Cultures were grown at 37°C until the OD_600_ reached 0.6. Protein expression was induced by adding 1 mM IPTG and incubation at 18 °C overnight. The cells were harvested by centrifugation for 10 min at 4,500 × *g*. The pellet was suspended in 10 mL ice cold lysis buffer (50 mM Tris HCl pH 7.5, 1% Triton 100, 200 mM NaCl, 1 mg/ml lysozyme, and 1 tablet of Protease inhibitor Cocktail (Roche)) and sonicated 6 times for 10 s with 10 s pauses at 200-300 W, followed by centrifugation at 10,000 × *g* for 60 min. The supernatant was purified using Ni-NTA agarose (QIAGEN) in accordance with the manufacturer’s instructions. The purified protein was desalted using the Amicon Ultra-15 Centrifugal Filter Unit (Merck) and the desalted protein was used for activity assays. Reactions were performed in 100 μL reaction volumes containing 100 mM Tris HCl pH 7.5, 2.5 μg protein, 5 mM UDP-α-D-glucose (Calbiochem, http://www.merckmillipore.com) and 250 μg/mL 17-hydroxygeranyllinalool, for 3 h at 30°C. The reaction was stopped by adding 400 μL methanol, and the mixture was used for HGL-DTG analysis using a high resolution time-of-flight mass spectrometer.

### Quantification of primary metabolites and phytohormones in plant tissues

For the quantitative analysis of primary metabolites and phytohormones, we used leaf material of 42-day-old elongated *N. attenuata* transgenic lines silenced in *GGPPS*, *UGT91T1*, *UGT74P5* and *UGT74P3/UGT74P5* expression as well as WT plants. Sample preparation and analysis of primary metabolites and phytohormones were performed based on Schaefer et al. (Schafer et al., 2016). Peak integration was performed using the operating Software MS Workstation (Bruker Daltonics). For the quantitative analysis of GPP, FPP and GGPP, we followed the protocol developed by Nagel et al. (Nagel et al., 2014).

### Quantification of 17-hydroxygeranyllinalool in plant tissues

Approximately 50 mg of leaf material was flash frozen and ground in liquid nitrogen and then aliquoted. Each aliquot was extracted with 500 μL 80% methanol containing 0.2 ng/μL testosterone, and shaken twice at 1000 strokes for 45 sec using a GenoGrinder 2000 (SPEX SamplePrep, http://www.spexsampleprep.com/). Homogenized samples were then centrifuged at 16 000 × *g* for 20 min at 4°C. The supernatant was centrifuged again at 16 000 × *g* for 20 min at 4°C and then diluted 1:10 with 80% methanol. We established a chromatographic method using a mixture of solvent A: water (Milli-Q, Merck, http://www.emdmillipore.com) with 0.1% acetonitrile and 0.05% formic acid and solvent B: methanol. U(H)PLC for the quantification of 17-HGL was performed using a Zorbax Eclipse XDB-C18 column (particle size 1.8 μm, column length 3.0 × 50 mm) from Agilent Technologies (http://www.agilent.com). The chromatographic separation was achieved using a U(HPLC) Advance (Bruker Daltonics) with the following gradient: 0-0.5 min at 10% of B, 0.5-1 min up to 90% of B, 1-4 min up to 100% of B, 4-5 min at 100% of B, 5-5.05 min down to 10% of B and from 5.05-6 min at 10% of B. The injection volume was 5 μL and the flow rate 0.5mL min^−1^.

MS detection was performed on an EvoQ Elite QqQ-MS equipped with a HESI (heated electrospray ionization) ion source (Bruker Daltonics) and the following HESI conditions: spray voltage 4500 V, cone temperature 350°C, cone gas flow 35^a^ (arbitrary units), heated probe temperature 500°C, probe gas flow 60^a^ and nebulizer gas 60^a^. Compounds were detected in multiple reaction monitoring (MRM) mode using specific precursor ion/product ion after positive ionization: the [M-H_2_O+H]^+^ ion was used as precursor for 17-HGL, 289/81 (quantifier), 289/107 (qualifier), the [M+H]^+^ ion was used as precursor for testosterone, 289/97 (quantifier), 289/109 (qualifier). Further details are given in supplemental figure 23. The areas were analyzed using the MS Workstation operating software from Bruker Daltonics (http://www.bruker.com).

### Structural determination by nuclear magnetic resonance spectroscopy (NMR)

17-Hydroxygeranyllinalool was provided by HPC24 Standards (www.hpc-standards.com) and the general structure was verified by 1D and 2D NMR spectroscopy (for ^1^H NMR, see Supplemental Figure 12). A Bruker AVANCE 400 NMR spectrometer (Bruker, Rheinstetten, Germany), equipped with a 5 mm BBFO probe, was used to record ^1^H NMR, DEPT 135, ^1^H-^1^H COSY, HSQC and HMBC spectra in MeOH-*d*_4_ at 300 K. Spectra were processed using TOPSPIN 3.1. (BrukerBiospin).

### Rapid screening of HGL-DTGs via ultrahigh-pressure liquid chromatography/time of flight mass spectrometry

All materials were ground in liquid nitrogen and split into aliquots of 10-100 mg fresh weight (FW), dependent on the tissue. Each aliquot was extracted in 100 μL - 1 mL extraction solution (80% methanol; ratio 1/10 FW/extraction solution) containing two steel balls by shaking twice at 1200 strokes/min for 60 sec using a Geno/Grinder 2000. Homogenized samples were then centrifuged at 16 000 × *g* for 20 min at 4°C. The supernatant was centrifuged again at 16 000 × *g* for 20 min at 4°C. Two independent chromatographic methods were used to resolve HGL-DTGs. Both methods used a mixture of solvent A: water with 0.1% acetonitrile and 0.05% formic acid and solvent B: acetonitrile and 0.05% formic acid. U(H)PLC for method A was performed using a Dionex UltiMate 3000 rapid separation LC system (Thermo Fisher, http://www.thermofisher.com), combined with a Thermo Acclaim RSLC 120 C18 column (particle size 2.2 μm, average pore diameter 120Å, column dimension 2.1 × 150 mm). Gradient elution steps were as follows: 0-0.5 min at 10% of B, 0.5–6.5 min up to 80% of B and 6.5–8 min at 80% of B followed by returning to the starting conditions and column equilibration. For method B sample gradient steps were as follows: 0–3 min at 10% B, 3–12 min up to 20% B, 12–17 min up to 35% B, 17–23 min up to 40% B, 23–25 min up to 45% B, 25–30 min up to 50% B, 30–40 min up to 90% B and 40–45 min at 90% B, followed by returning to the starting conditions and column equilibration. The injection volume was 2 μL and the flow rate 0.4 mL/min for method A and B.

MS detection was performed using a micrOTOF-Q II, an Impact II and a maXis UHR-Q-TOF-MS system (Bruker Daltonics) equipped with an electrospray ionization (ESI) source operating in positive ion mode. ESI conditions for the micrOTOF-Q II system were: end plate offset 500 V, capillary voltage 4500 V, capillary exit 130 V, dry temperature 180°C and a dry gas flow of 10 L min^−1^. ESI conditions for the Impact II UHR-Q-TOF-MS system were capillary voltage 4500 V, end plate offset 500 V, nebulizer 2 bar, dry temperature 200°C and a dry gas flow of 8 L min^−1^. ESI conditions for the maXis UHR-Q-TOF-MS system were capillary voltage 4500 V, end plate offset 500 V, nebulizer 1.8 bar, dry temperature 200°C and a dry gas flow of 8 L min^−1^. MS data were collected over a range of *m/z* from 100 to 1600. Mass calibration was performed using sodium formate (50 mL isopropanol, 200 μL formic acid, 1 mL 1 M NaOH in water). Data files were calibrated using the Bruker high-precision calibration algorithm. Lock mass calibration was performed for the profiling of the stable lines using signal *m/z* 622.0289 (molecular formula C_12_H_19_F_12_N_3_O_6_P_3_) from the ESI Tuning Mix (Agilent Technologies, http://www.agilent.com). MS/MS experiments were performed using AutoMS/MS runs at various CID voltages from 12.5 to 22.5 eV for ammonium adducts. Instrument control, data acquisition and reprocessing were performed using HyStar 3.1 (Bruker Daltonics). Molecular formulae were determined using SmartFormula 3D. SmartFormula calculates the elemental compositions from accurate mass as well as the isotopic pattern information using MS (SmartFormula) and MS + MS/MS information (SmartFormula 3D) (Krebs and Yates, 2008; Kind and Fiehn, 2010). The mass tolerance was set to 4 mDa, and the filter H/C element ratio was set between 1 and 3. Isotope peaks were assigned using the Simulate Pattern Tool of the DataAnalysis software version 4.2 (Bruker Daltonics). We used QuantAnalysis (Bruker Daltonics) to integrate the peak areas.

### Dereplication of HGL-DTGs

The dereplication workflow relies on a comprehensive MS and MS/MS database constructed by our group for HGL-DTGs of several solanaceous species (Heiling et al., 2016) and a detailed rule-set for the annotation of fragmentation patterns of the different moieties decorating the 17-hydroxygeranyllinalool (17-HGL) aglycone. The MS and MS/MS database is based on the retention time and mass spectrometric data of purified HGL-DTGs which are used as authentic standards. Novel HGL-DTGs are annotated based on their spectral similarity to the MS and MS/MS in-house database. For the visualization and identification of HGL-DTG profiles, we computed the extracted ion chromatogram EIC *m/z* 271.2420. This *m/z* fragment corresponds to the 17-HGL aglycone lacking both hydroxyl-groups and is produced by in-source fragmentation during ionization for all HGL-DTGs independently of the type and degree of metabolic decorations. The trace returned for *m/z* 271.2420 allows for the visualization of the complete HGL-DTG chemotype and to rapidly assess variations within this chemotype that result from the single gene manipulations. Following application of the de-replication workflow, we subdivided, based on the presence of diagnostic *m/z* signals, the chemotype between rhamnosylated and non-rhamnosylated HGL-DTGs. Lyciumoside I and lyciumoside II and their malonylated derivatives corresponded to non-rhamnosylated HGL-DTGS, while lyciumoside IV, attenoside and nicotianoside III as well as their malonylated forms, represented the major rhamnosylated compounds. We provide a detailed description of the identification of known and novel HGL-DTGs in Supplemental Tables 1a/b. The identification levels are based on community standards reported in (Sumner et al., 2007). Raw MS metabolomics data have been deposited in the open metabolomics database Metabolights, www.ebi.ac.uk/metabolights (accession no. MTBLS1819).

### Statistical analysis

Data were analyzed using Excel (Microsoft, http://www.microsoft.com), SPSS 20.0 (SPSS Inc, http://www-01.ibm.com/software/analytics/spss/) and RStudio (RStudio Inc, https://www.r-project.org) using the package xlsx. Unless otherwise stated, parametric data were compared using ANOVA followed by Fisher LSD/Holm-Bonferroni post hoc tests or Mann-Whitney-Wilcox Pairwise Test (for heteroskedastic data). The phylogenetic tree was constructed using the maximum-likelihood method in MEGA5.0 (http://www.megasoftware.net/).

## ACCESSION NUMBERS

For the analysis of the HGL-DTG pathway, we used the following GenBank accessions for *N. attenuata*: NaGLS, KJ755868; NaGGPPS, EF382626; NaUGT74P3, KX752207; NaUGT91T1, KX752209; NaUGT74P5, KX752208), *N. obtusifolia* (NoUGT74P4, KX752210; NoUGT74P6, KX752211; NoUGT91T1-like, MG051326). The MS metabolomics data set has been deposited in the open metabolomics database Metabolights, www.ebi.ac.uk/metabolights (accession no. MTBLS1819).

## Supplemental Data

**Supplemental Method File 1:** Phylogenetic characterization of the UDP-glycosyltransferases in *N. attenuata* and *N. obtusifolia*

**Supplemental Method File 2:** Altered general and specialized metabolite levels

**Supplemental Figure 1:** Alignment of the UGT C-terminal consensus sequence of 112 family 1 glycosyltransferases from *N. attenuata* and *N. obtusifolia*.

**Supplemental Figure 2:** Phylogenetic analysis of the *N. attenuata* UGT superfamily shows 16 major groups

**Supplemental Figure 3:** Amino acid composition of all identified UGTs of the superfamily 1 in *N. attenuata*

**Supplemental Figure 4:** Phylogenetic relationships and herbivory-induced tissue-specific expression of 110 predicted UDP-glycosyltransferases (UGT)

**Supplemental Figure 5:** Transcriptomic variation of UGTs after treatment with OS in *N. attenuata*

**Supplemental Figure 6:** Phylogenetic tree analysis for the UDP-glycosyltransferases used for stable and transient silencing.

**Supplemental Figure 7:** Silencing efficiency for the three transiently-silenced 17-HGL-DTG biosynthetic UGTs in pTVUGT91T1, pTVUGT74P3 and pTVUGT74P5.

**Supplemental Figure 8:** Co-silencing efficiency of UGT74P3 and UGT74P5 in pTVUGT74P3 and pTVUGT74P5.

**Supplemental Figure 9:** Silencing efficiency for the three transiently-silenced 17-HGL-DTG biosynthetic UGTs in pTVUGT91T1-like, pTVUGT74P4 and pTVUGT74P6 in *N. obtusifolia*.

**Supplemental Figure 10a/b:** Mass spectrometric characterization and annotation of novel HGL-DTGs in transiently-silenced *N. obtusifolia* plants impaired in No*UGT74P4* and No*UGT74P6* expression.

**Supplemental Figure 11a/b/c:** Characterization and annotation of novel HGL-DTGs via MS/MS in stably-silenced *N. attenuata* plants impaired in *UGT74P3* and *UGT74P5* expression.

**Supplemental Figure 12:** ^1^H NMR spectrum of synthetic 17-hydroxygeranyllinalool (17-HGL, HPC24 Standards)

**Supplemental Figure 13a/b/c:** Morphological characterization of *N. attenuata* plants transiently-silenced in *UGT91T1, UGT74P3* and *UGT74P5* expression

**Supplemental Figure 14:** Morphological characterization of *N. obtusifolia* plants transiently-silenced in No*UGT91T1-like*, No*UGT74P4*, No*UGT74P6* and No*UGT74P4*/*UGT74P6* expression

**Supplemental Figure 15:** Metabolite profiling and morphological characterization of *N. attenuata* plants transiently-silenced via virus-induced gene silencing (VIGS) of geranyllinalool synthase (GLS)

**Supplemental Figure 16:** Southern Blot

**Supplemental Figure 17a/b/c:** Morphological characterization of the stable transformed *IRugt74p5* Line A, Line B and *IRugt74p3/ugt74p5*.

**Supplemental Figure 18:** Characterization of growth parameters in IR*ugt91t1*, IR*ugt74p5*, IR*ugt74p3/ugt74p5* and IR*ggpps*

**Supplemental Figure 19:** Morphological characterization of IR*ugt91t1*

**Supplemental Figure 20:** Overall abundance of HGL-DTGs

**Supplemental Figure 21:** Disrupting HGL-DTG glycosylation reorganizes general, specialized and hormonal metabolic pathways

**Supplemental Figure 22:** Characterization of free prenyldiphosphates in IR*ggpps* and WT

**Supplemental Figure 23:** Quantitative 17-HGL method

**Supplemental Figure 24:** 17-HGL concentration in IR*ggpps* and WT plants transiently transformed with pTV00, pTVUGT74P3 and pTVUGT74P5

**Supplemental Figure 25:** Characterization of the morphological phenotype of WT and IR*ggpps* plants transiently transformed with pTV00.

**Supplemental Figure 26:** Characterization of the morphological phenotype of WT and IR*ggpps* plants transiently transformed with pTVUGT74P3.

**Supplemental Figure 27:** Characterization of the morphological phenotype of WT and IR*ggpps* plants transiently transformed with pTVUGt74P5.

**Supplemental Table 1:** Molecular weight of UGTs in *N. attenuata*

**Supplemental Table 2:** Phylogenetic grouping of 107 UGTs in *N. attenuata*

**Supplemental Table 3:** UGT Amino acid composition in *N. attenuata*

**Supplemental Table 4:** SignalIP4.1 – signal peptide cleavage sites

**Supplemental Table 5a/b:** Pearson Correlation of all UGTs to NaGLS and NaGGPPS

**Supplemental Table 6:** MS/MS Measurements for HGL-DTGs in *N. obtusifolia*

**Supplemental Table 7:** MS/MS Measurements for HGL-DTGs in *N. attenuata*

**Supplemental Table 8a-b:** Tissue-specific HGL-DTG modulation

**Supplemental Table 9:** Primers

**Supplemental Data 1:** Relative UGT expression vs. time in *N. attenuata*

**Supplemental Data 2:** HGL-DTG profiles of transiently-silenced *N. attenuata* plants

**Supplemental Data 3:** HGL-DTG profiles of transiently-silenced NaGLS *N. attenuata* plants

**Supplemental Data 4:** HGL-DTG profiles of transiently-silenced *N. obtusifolia* plants

**Supplemental Data 5:** HGL-DTG profiles of stably-silenced *N. attenuata* plants

**Supplemental Data 6:** General and specialized metabolites in stably-silenced *N. attenuata* plants

**Supplemental Data 7:** Phytohormone profiles in stably-silenced *N. attenuata* plants

**Supplemental Data 8:** Statistical analysis of the performance assay and consumed leaf disk mass

## ACKNOWLEDGEMENTS

We thank the gardening staff at the Max Planck Institute for Chemical Ecology. We thank Thomas Hahn, Nicolas Heinzel, Alexander Weinhold, Mario Kallenbach and Matthias Schoettner for the analytical and technical support as well as Dapeng Li and Felipe Yon for the fruitful discussions. We thank Raimund Nagel for his help with the terpene quantification. Additionally, we thank Eric McGale and Jyotasana Gulati for their statistical wisdom and Michael Court for assistance naming UGTs. We thank the Max Planck Society and the International Max Planck Research School on the Exploration of Ecological Interactions with Chemical and Molecular Techniques for financial support. We acknowledge the European Research Council advanced grant ClockworkGreen to I.T.B. (number 293926; I.T.B.) for funding.

## Notes

https://www.ebi.ac.uk/metabolights/MTBLS1819/descriptors

### Competing Interest Statement

The authors have declared no competing interest.

