## Supplemental Method File 1 for "Specific decorations of 17-hydroxygeranyllinalool diterpene glycosides solve the autotoxicity problem of chemical defense in *Nicotiana attenuata*"

GTs are highly divergent, polyphyletic, and represent one of the largest multigene super-families in plant genomes (Mackenzie et al., 1997; Ross et al., 2001). GTs catalyze the transfer of activated sugars as donor molecules to specific acceptors, including sugars, lipids, proteins, nucleic acids as well as small hydrophilic molecules (Weadge, 2000; Lairson et al., 2008). GTs have been classified into 103 families according to their sequence similarity to other carbohydrate-active enzymes, their catalytic mechanisms (inverting and retaining), 3D-structures (Coutinho et al., 2003), sugar donors, transferred sugars, acceptors and the combination of modules which can be catalytic or not (Lombard et al., 2014) as well as the presence of conserved sequence motifs (Campbell et al., 1997; 1998, CAZY - http://www.cazy.org). Family 1 corresponds to the uridine-5'-phosphate (UDP)-glycosyltransferases (UGTs) which are characterized by the utilization of UDP-activated sugars, like UDP-glucose, UDP-rhamnose, UDP-galactose and UDP-xylose as donor molecules (Merken and Beecher, 2000) for the glycosylation of complex structurally variable substrates like flavonoids, terpenes, and even phytohormones. While most UGTs perform O-glycosylations, for some xenobiotics however, UDP-glucose is used to glycosylate N-, S- or even C-linkages (Brazier-Hicks et al., 2009). Plant UGTs are promiscuous and are known to have broad substrate specificity, which is generally limited by regiospecificity (Lim et al., 2003). However, in some cases UGTs have also been shown to be highly specific. The diversity of used substrates is thought to result from the high variability in the N-terminal region of UGTs (Bowles et al., 2005; Bowles et al., 2006). The C-terminal region contains a conserved 44 amino acid sequence motif known as the plant secondary product glycosyltransferase (PSPG)-box, which is thought to be the UDP-sugar binding site (Vogt and Jones, 2000) and is often the only region of significant similarity in sequence alignments. Plant UGTs of the family 1 have been thoroughly studied across the plant kingdom, including Arabidopsis thaliana (Li et al., 2001; Ross et al., 2001; Paquette et al., 2003), Glycine max (Rehman et al., 2016), Gossypium hirsutum (Huang et al., 2015), Linum usitatissimum (Barvkar et al., 2012), Medicago truncatula (Achnine et al., 2005) and Zea mays (Li et al., 2014). These genome-wide investigations provide systematic and global insights into the relationship between the structure and function of this multigene family.

**Genome-wide inference and phylogenetic analysis of *N. attenuata* UGTs**

A genome-wide survey of *N. attenuata* identified a total of 107 putative UGT sequences containing the PSPG motif at the C-terminus (Supplemental Figure 1). The length of the deduced proteins varied from 437 – 514 aa, averaging 474 aa with a predicted molecular weight ranging from 49 kDa to 58 kDa (Supplemental Table 1). All UGT sequences started with a methionine, none of these sequences contained premature stop codons. The constructed phylogenetic tree resulted in the classification of UGT protein sequences into 18 major groups (A-R) (Supplemental Figure 2). These groups are consistent with the previously described classification of UGTs in Arabidopsis (Li et al., 2001; Ross et al., 2001), maize (Li et al., 2014), flax (Barvkar et al., 2012) or cotton (Huang et al., 2015; Rehman et al., 2016) with the exception of the subfamily UGT95 which did not cluster with the UGT92 subfamily as part of group M (Huang et al., 2015). We therefore created a new group R, specific to *N. attenuata* that contains the UGT95 proteins. Group Q was absent in the classification of *N. attenuata* UGTs. Proteins classified as part of the same phylogenetic group exhibit percentages of aa similarity ranging from 22 % to 100 % (Supplemental Table 2). Highest average similarities were detected in group G with 73% and groups K and M with 59%, while the lowest were detected in groups A (23%), E (24%) and L (22%). Altogether, the 105 UGTs of *N. attenuata* clustered into 26 subgroups (sequence similarity of ~40% according to UGT nomenclature) and only two UGT sequences (NaUGTg20981; NaUGTg16298) did not cluster in any subgroup and remained unclassified.

To characterize the identified UGTs, we analyzed the amino acid composition and compared it to its soybean relatives. Leucine was the most abundant amino acid across all UGTs from *N. attenuata* (10%). Cysteine, histidine, methionine, tryptophan and tyrosine were the least common amino acids (1.6 – 2.7%) (Supplemental Figure 3, Supplemental Table 3). These results are consistent with those observed in soybean (Rehman et al., 2016).

We used SignalP4.1 to identify whether UGT proteins contain signal peptides for subcellular localization and detected that only two sequences (NaUGTg28668, NaUGTg29262) contained such motifs (Supplemental Table 4).

Using a full-transcriptome microarray dataset obtained from leaf and root tissues collected at several time-points following simulated leaf herbivory by application of *Manduca sexta* oral secretions (OS), we detected that, of the 107 UGTs identified in *N. attenuata*, 76 were expressed in either leaves or roots, while 31 were not expressed in these two tissue types (Supplemental Figure 4, Supplemental Data 1). The most pronounced changes in UGT transcript abundance appeared after 1h in locally OS-treated leaves while in systemic leaves and roots, induced levels of UGT expression peaked after 5 h. Consistent with the observation that UGTs often play pivotal metabolic functions in response to insect herbivory, only 12 UGTs were not affected by the OS treatment. To explore the tissue-level expression of the identified UGTs, we mined a previously published RNA-seq atlas established for 21 different tissue types from *N. attenuata* (Brockmoller et al., 2017). In addition to the previously mentioned 76 UGTs detected in leaves and roots after simulated herbivory, we found 17 UGTs expressed in different tissues other than leaves and roots. For the remaining 14 UGTs we did not detect any significant transcript abundance levels (Supplemental Figure 4). 45 UGTs were differentially expressed (2-fold change relative to controls) 1h after the simulated herbivory treatment, 41 UGTs after 5 h and 27 UGTs after 17 h (Supplemental Figure 5).

**Phylogenetic characterization of the UGT family 1 in *N. attenuata***

UGTs represent 0.29% (137/47912), of all genes in flax (Barvkar et al., 2012), 0.33 – 0.38% (142/37505, *G. raimondii*; 146/40134 *G. arboreum*; 196/59089, *G. hirsutum)*; in different cotton species (Huang et al., 2015) and up to 0.44% of all genes in *Arabidopsis (120/27416*, (Paquette et al., 2003)). In wild and cultivated tobacco species, only a handful of family 1 UGTs have been functionally characterized. This includes a salicylic acid glucosyltransferase (SAGT – (Lee and Raskin, 1999)), several UGTs responsible for the glucosylation of phenolics, especially naphthols (*Nt*GT1a, *Nt*GT1b and *Nt*GT3 – (Taguchi et al., 2001; Taguchi et al., 2003)) and two putative flavonoid UDP-glycosyltransferases (UGT-A, UGT-B – Li et al., 2016). To enrich our understanding of this family in *N. attenuata*, we performed a genome-wide analysis for the identification of UGTs and analyzed their phylogenetic relationships. We identified 107 putative UGTs and showed that the family-1 UGTs represent 0.32% (107/33449 – Xu et al., 2017) of the expressed genes in the genome, which is less than in *Arabidopsis* but about the same as in flax or cotton.

Additionally, we found that five phylogenetic groups, namely A, D, E, L and O, seem to have expanded more than others (Supplemental Table 2) in *N. attenuata*. This observation has been made in earlier studies in other higher plants (Caputi et al., 2012; Rehman et al., 2016). We foresee that this phylogenetic analysis in combination with expression studies provides an instrumental data platform for further characterization of UGT functions in the genus *Nicotiana*.

**Material and Methods**

**Identification of UDP-glycosyltransferase sequences in *N. attenuata***

Sequences of the *N. attenuata* genome (MJEQ00000000) and a 454 *N. attenuata* shotgun assembly (GBGF00000000) were used. To identify members of the UGT family, we used the 44-amino acid conserved sequence of the plant secondary product glycosyltransferase (PSPG)-box motif as a query to perform a local BLASTP search. The e-Value threshold was set to 1e-10. All UGT candidates were verified using HMMER (<http://www.ebi.ac.uk/Tools/hmmer>) to confirm the presence of the UDP-glycosyltransferase domain (pfam00201). The identified UGTs were named based on the *N. attenuata* gene identifiers. Verified UGTs with a proven function were classified and named based on the HUGO Gene Nomenclature Committee (Mackenzie et al., 1997).

**Phylogenetic analysis of *N. attenuata* UGTs**

Identified UGT sequences were aligned using Clustal W with default gap penalties, the phylogenetic tree was constructed using maximum-likelihood (JTT matrix-based model, bootstrap value: 1000 replicates) and neighbor joining (bootstrap value: 1000, p-distance and pairwise deletion) in MEGA 5.0 (<http://www.megasoftware.net>). Twenty-eight reference UGT peptide sequences belonging to various phylogenetic groups (A-O) were used for the phylogenetic comparison (Supplemental Figure 2). Subgroups were classified based on sequence similarity to the twenty-eight UGTs. Sequence similarity of the subgroup UGT93 was calculated based on the alignment to ZOG1 and ZOX1 from *Phaseolus vulgaris* (ZOX1 AAD51778; ZOG1 AAD04166) via Geneous (<http://www.geneous.com>). The total amino acid composition was calculated using the MEGA sequence data explorer tool. The molecular mass was calculated using Protein Molecular Weight (http://www.bioinformatics.org/sms2/protein_mw.html).

**Microarray data sets**

To explore the temporal expression dynamics of *N. attenuata* UGTs, we mined a dataset produced by our laboratory that is publicly available at the Gene Expression Omnibus database (accession number GSE30287), and consists of 150 published microarray expression profiles. This microarray data set was originally published in (Kim et al., 2011) and its experimental design was as follows. To simulate *Manduca sexta* feeding, the laminas of three leaves per plant (two source leaves at nodes +2 and +1 and one source-sink transition leaf at node 0) were mechanically wounded with a fabric pattern wheel on both sides of the midrib, and immediately, 20 µL of *M. sexta* OS (diluted 1:10 in water) was applied to the fresh puncture wounds (W+OS). For each time point (1, 5, 9, 13, 17, and 21 h after treatment), treated leaves or control leaves at the same nodal positions, systemic leaves (two sink leaves at nodes −1 and −2), and the complete root system were collected from six plants and immediately flash frozen in liquid nitrogen. As described by Gulati et al. (Gulati et al., 2013; Gulati et al., 2014), raw intensities of the microarray data-set were normalized using the 75th percentile value procedure and log_2_ and baseline transformed prior to statistical analysis. We compared the expression of all identified UGTs at 1h, 5h and 17h in local, systemic and root tissue using Tukeys post hoc test (p-Value < 0.05). The correlation to *NaGGPPS* and *NaGLS* was performed using Pearson correlation calculations across all 134 microarrays. Significance levels for correlation values (*r*) were determined following the number of transcript pairs (*n*) using the equation t = r × (n-2)^0.5^ / (1-r) ^0.5^.

For the expression of the UGTs in other tissues we analyzed an RNAseq dataset (PRJNA317743) of 21 different tissues in *N. attenuata*.

For the construction of the UDP-glycosyltransferase tree in *N. attenuata*, we used the following GenBank accessions: NaUGT g00526 - KX752100, NaUGT g00527 - KX752101, NaUGT g01854 - KX752102, NaUGT g02083 - KX752103, NaUGT g02093 - KX752104, NaUGT g02821 - KX752105, NaUGT g02942 - KX752106, NaUGT g03342 - KX752107, NaUGT g3727 - KX752108, NaUGT g03748 - KX752109, NaUGT g04160 - KX752110, NaUGT g04820 - KX752111, NaUGT g05060 - KX752112, NaUGT g05219 - KX752113, NaUGT g05426 - KX752114, NaUGT g05573 - KX752115, NaUGT g06355 - KX752116, NaUGT g08104 - KX752117, NaUGT g9179 - KX752118, NaUGT g09326 - KX752119, NaUGT g09327 - KX752120, NaUGT g10741 - KX752121, NaUGT g11159 - KX752122, NaUGT g11521 - KX752123, NaUGT g11522 - KX752124, NaUGT g11850 - KX752125, NaUGT g11851 - KX752126, NaUGT g12088 - KX752127, NaUGT g13005 - KX752128, NaUGT g13538 - KX752129, NaUGT g13945 - KX752130, NaUGT g15196 - KX752131, NaUGT g16288 - KX752132, NaUGT g16298 - KX752133, NaUGT g16426 - KX752134, NaUGT g18508 - KX752135, NaUGT g18870 - KX752136, NaUGT g19018 - KX752137, NaUGT g19190 - KX752138, NaUGT g19344 - KX752139, NaUGT g19346 - KX752140, NaUGT g20123 - KX752141, NaUGT74P5 - KX752142, NaUGT g20981 - KX752143, NaUGT g21652 - KX752144, NaUGT g21654 - KX752145, NaUGT g21846 - KX752146, NaUGT g22203 - KX752147, NaUGT g22204 - KX752148, NaUGT g22205 - KX752149, NaUGT g22481 - KX752150, NaUGT g22983 - KX752151, NaUGT g23136 - KX752152, NaUGT g23176 - KX752153, NaUGT g23995 - KX752154, NaUGT g24514 - KX752155, NaUGT g24515 - KX752156, NaUGT g24585 - KX752157, NaUGT g24697 - KX752158, NaUGT g24726 - KX752159, NaUGT g25918 - KX752160, NaUGT g26396 - KX752161, NaUGT91T1 - KX752162, NaUGT g27096 - KX752163, NaUGT g27477 - KX752164, NaUGT g28046 - KX752165, NaUGT g28047 - KX752166, NaUGT g28309 - KX752167, NaUGT g28311 - KX752168, NaUGT g28331 - KX752169, NaUGT g28668 - KX752170, NaUGT g29220 - KX752171, NaUGT g29262 - KX752172, NaUGT g29493 - KX752173, NaUGT g30507 - KX752174, NaUGT g30575 - KX752175, NaUGT g30577 - KX752176, NaUGT g30863 - KX752177, NaUGT g30898 - KX752178, NaUGT g31158 - KX752179, NaUGT g31196 - KX752180, NaUGT g32016 - KX752181, NaUGT g32892 - KX752182, NaUGT g32894 - KX752183, NaUGT g33134 - KX752184, NaUGT g34091 - KX752185, NaUGT g34877 - KX752186, NaUGT g34878 - KX752187, NaUGT g34991 - KX752188, NaUGT g35131 - KX752189, NaUGT g35881 - KX752190, NaUGT g35911 - KX752191, NaUGT g36017 - KX752192, NaUGT g36713 - KX752193, NaUGT g37675 - KX752194, NaUGT g38211 - KX752195, NaUGT g38353 - KX752196, NaUGT g38610 - KX752197, NaUGT g38855 - KX752198, NaUGT g39264 - KX752199, NaUGT g40324 - KX752200, NaUGT g40325 - KX752201, NaUGT g40889 - KX752202, NaUGT g41120 - KX752203, NaUGT74P3 - KX752204, NaUGT g41996 - KX752205, NaUGT g43590 - KX752206.
