## Supplemental Method File 2 for "Specific decorations of 17-hydroxygeranyllinalool diterpene glycosides solve the autotoxicity problem of chemical defense in *Nicotiana attenuata*"

**Disrupting HGL-DTG biosynthetic flux influences central metabolism**

Quantitative profiling of 40 general and specialized metabolites (23 amino acids and biogenic amines, 4 small organic acids, 10 phenylpropanoids and derivatives and 4 sugars) using U(H)PLC-triple-quadrupole MS was performed on extracts from IR*ugt91t1*, IR*ugt74p5*, IR*ugt74p3/ugt74p5*, IR*ggpps* and WT leaves (Supplemental Figure 21, Supplemental Data 5). The IR*ugt91t1* leaf extracts of line A and B contained modest but significantly higher levels of α-keto-glutaric acid and Line B also contained higher levels of quercetin but not significantly higher levels of other flavonoids, such as quercetin-3-*O*-glucoside, quercetin-3-*O*-sophoroside, rutin or kaempferol-3-*O*-rutinoside. IR*ugt74p5* and IR*ugt74p3/ ugt74p5* leaf extracts exhibited higher levels of the biogenic amines, tyramine and tryptamine, of the amino acids L-tryptophan and L-cysteine, α-keto-glutaric acid and the phenylpropanoids caffeic acid, ferulic acid and synapylaldehyde. On the other hand, lower levels of glucuronic acid and shikimic acid were also detected. Interestingly, IR*ggpps* exhibited most changes in central metabolism, especially among amino acids. IR*ggpps* extracts contained higher levels in 11 amino acids (L-glutamine, L-histidine, L-lysine, L-proline, L-phenylalanine, L-threonine, L-tryptophan, L-tyrosine, L-valine, L-cysteine and L-methionine). Especially L-tryptophan (increased by more than 10-fold), L-histidine (increased by 6.6-fold) and L-glutamine (increased by 6-fold) levels were altered dramatically compared to those of WT. The biogenic amine tyramine, the phenylpropanoid derivatives, scopoletin and scopolin, and the small organic acids, succinic acid and α-keto-glutaric acid (138 × higher in IR*ggpps* than in WT) were increased as well. Furthermore L-alanine, L-aspartic acid, L-glutamic acid and shikimic acid were significantly reduced. Noteably, the concentration of L-aspartic acid was 13 times lower. Likewise, the free sugars glucose (7.6-fold), fructose (4.6-fold) and glucuronic acid (2-fold) were highly reduced in *IRggpps* as well. Furthermore, we investigated the amount of the free prenyldiphosphates, GPP, FPP and GGPP, in IR*ggpps* compared to WT. We observed a significant reduction of GGPP (4.3-fold) and an increase in GPP (9.7-fold) and FPP (5.1-fold) in *IRggpps* (Supplemental Figure 22).

**Leaves of plants impaired in HGL-DTG biosynthesis exhibit altered phytohormone levels**

We observed a severe developmental phenotype in *N. attenuata* plants impaired in *UGT74P5* and *UGT74P3/UGT74P5* expression*, which* was similar to that reported for the phenotypes of plants abrogated in phytohormone signaling or biosynthesis (Ueguchi-Tanaka et al., 2005; Rodó et al., 2008). For this reason, we analyzed the phytohormone profiles of all lines with strong developmental and growth phenotypes. The quantitative profiling of 26 phytohormones and derivatives using U(H)PLC-triple-quadrupole-MS was performed on extracts from IR*ugt91t1*, IR*ugt74p5*, IR*ugt74p3/ugt74p5*, IR*ggpps* and wild type leaves (Supplemental Figure 21B, Supplemental Data 6).

First, our attention was directed to the gibberellin pathway, which is essential for many developmental processes in plants and therefore, when disturbed, could be responsible for the observed morphological alterations. We analyzed the most active gibberellins GA_1_, GA_3_, GA_4_ and GA_7_ (Schneider et al., 1989; Olszewski et al., 2002) for altered levels in leaf tissues. Of these compounds, we only detected GA_3_ which was not significantly changed in any stable construct compared to wild type. Additionally we were able to measure large changes in GA_8_, GA_20_ and GA_51_. We showed that leaf extracts of IR*ugt74p5* line A and B and IR*ugt74p3/ugt74p5* contained higher gibberellin GA_20_ levels. The highest concentration was measured in the IR*ugt74p5* line A exhibiting 114 pmol/g FW. In IR*ugt74p5* line B, a concentration of 53.6 pmol/g FW was observed. The heterologous double construct with the most severe morphological phenotype, accumulated only 12.9 pmol/g FW, inconsistent with the hypothesis that GA_20_ would be responsible for these alterations. Neither IR*ugt91t1* nor IR*ggpps* plants showed detectable levels of GA_20_. Additionally, GA_8_ was 3.8 times (Line A) and 3.2 times (Line B) higher in IR*ugt74p5* compared to WT, but not detectable in IR*ugt74p3/ugt74p5.* In contrast, GA_51_ was only detectable in WT and both IR*ugt91t1* lines.

Further we investigated cytokinin levels which are essential for plant development (Mok and Mok, 1994). Our analysis of both IR*ugt91t1* lines revealed that they closely resembled WT plants in their cytokinin levels. Only dihydrozeatin (DHZ) could not be detected and line B had slightly increased levels of *cis*-zeatin (cZ). Both IR*ugt74p5* transformed lines showed increased levels of isopentenyladenine (IP), cZ and *cis*-zeatin-N7-glycoside (cZ7G). DHZ and dihydrozeatin-riboside (DHZR) were not detected and *cis*-zeatin-O-glucoside-riboside (cZROG) was reduced. Additionally *trans*-zeatin-riboside (tZR) was reduced in IR*ugt74p5* line A*.* IR*ugt74p3/ugt74p5* exhibited higher levels of cZ7G and lower levels of cZROG. In addition, DHZR could not be detected in the heterologous double construct. Interestingly, strong alterations were detected in IR*ggpps*. Leaf extracts of IR*ggpps* plants contained higher concentrations of the cytokinins cZ, cZR, DHZ, DHZR, czROG, *trans*-zeatin-N7-glycoside (tZ7G), cZ7G and dihydrozeatin-N7-glycoside (DHZ7G). The highest induction was observed for DHZR, which was increased 24-fold. The higher levels of cytokinins could explain the shorter stems and delayed flowering of IR*ggpps* plants.

Additionally to cytokinins and gibberellins, we analyzed jasmonate levels for all constructs. Interestingly strong alterations were detected in both IR*ugt74p5* lines. Particularly high concentrations of COOH-JA-Ile, OH-JA and JA-Ile were observed. The only difference between both IR*ugt74p5* lines was the high concentration of jasmonic acid (JA) in Line A, which could not be detected in Line B. Surprisingly IR*ugt74p3/ugt74p5* transformed plants showed no significant differences from the levels found in WT plants. However, COOH-JA-Ile, OH-JA and JA-Ile were clearly detectable in the heterologous double construct. In WT plants only JA-Ile and OH-JA-Ile could be observed. Furthermore, the phytohormones abscisic acid (ABA), indole acetic acid (IAA) and salicylic acid (SA) were measured in the leaf extracts. IR*ggpps* showed highly increased IAA levels (3.4-fold) and about half the levels in ABA. No significant differences in the concentration of SA were observed across all tested constructs.

The diverse morphological deformities of IR*ugt74p5* and IR*ugt74p3/ugt74p5* plants might be a nonspecific stress response triggered by the toxicity of 17-HGL (Bowles et al., 2005; Bowles et al., 2006; Mylona et al., 2008; Naoumkina et al., 2010; Itkin et al., 2011), which interferes with phytohormone homeostasis or the accumulation of yet unknown compound classes. For example, *ugt74b1* mutants of *A. thaliana,* which are impaired in glucosinolate biosynthesis, display phenotypes of auxin overproduction, such as epinastic cotyledons and incomplete leaf vascularization (Grubb et al., 2004). Over-expression of a zeatin O-glucosylation gene in *Zea mays* leads to growth retardation and tassel-seed formation (Rodo et al., 2008). Furthermore, *gid1* mutants impaired in a soluble receptor for gibberellins in *Oryza sativa* displayed a severe dwarf phenotype with a loss of GA-responsiveness (Ueguchi-Tanaka et al., 2005). When we analyzed gibberellins in IR*ugt74p5* and IR*ugt74p3/ugt74p5* plants, we found no difference in gibberellic acid (GA_3_) contents, but found high levels of GA_20_, which is thought to be a mobile signal in the elongation of internodes and flower induction (Proebsting et al., 1992; Ross et al., 2001). The increased levels of GA_20_ is roughly consistent with a phenotype observed in the later developmental stages of IR*ugt74p5* and IR*ugt74p3/ugt74p5* plants as well as in their T_0_-transformants, namely the prevalence of multiple internodes with stalled flower buds.
