## Supplemental Table 2 for "Specific decorations of 17-hydroxygeranyllinalool diterpene glycosides solve the autotoxicity problem of chemical defense in *Nicotiana attenuata*"

Supplemental Table 2: Phylogenetic grouping of 107 UGTs in *N. attenuata*

| Gr. No. | Group | Included families | NO. of sequences | % Similarity of full length protein | | |
|---|---|---|---|---|---|---|
| | | | | MIN | MAX | AVERAGE |
| 1 | A | 79,91,94 | 16+1 | 22.85 | 100.00 | 36.66 |
| 2 | B | 89 | - | - | - | - |
| 3 | C | 90 | 1 | - | - | - |
| 4 | D | 73 | 12 | 35.25 | 92.40 | 45.96 |
| 5 | E | 71,72,88,96, 99 | 19 | 24.25 | 100.00 | 37.98 |
| 6 | F | 78 | 2 | 49.78 | 49.78 | 49.78 |
| 7 | G | 85 | 6 | 56.08 | 95.93 | 73.24 |
| 8 | H | 76 | 1 | - | - | - |
| 9 | I | 83,712 | 3 | 38.67 | 51.87 | 44.19 |
| 10 | J | 87 | 1 | - | - | - |
| 11 | K | 86 | 3 | 54.70 | 63.05 | 59.13 |
| 12 | L | 74,75,84 | 14+2 | 21.93 | 100.00 | 40.85 |
| 13 | M | 92 | 2 | 59.99 | 59.99 | 59.99 |
| 14 | N | 82 | 1 | - | - | - |
| 15 | O | 98, 93 | 17 | 35.04 | 82.58 | 47.59 |
| 16 | P | 709 | 5 | 47.42 | 55.23 | 51.70 |
| 17 | Q | 705 | - | - | - | - |
| 18 | R | 95 | 2 | 57.99 | 57.99 | 57.99 |
