## Supplemental Table 3 for "Specific decorations of 17-hydroxygeranyllinalool diterpene glycosides solve the autotoxicity problem of chemical defense in *Nicotiana attenuata*"

Supplemental Table 3: UGT Amino acid composition in *N. attenuata* . All frequencies are given in percent.

| Amino acid sequence | Ala | Cys | Asp | Glu | Phe | Gly | His | Ile | Lys | Leu | Met | Asn | Pro | Gln | Arg | Ser | Thr | Val | Trp | Tyr | Total |
|---|---|---|---|---|---|---|---|---|---|---|---|---|---|---|---|---|---|---|---|---|---|
| NaUGT g43590 | 5.5 | 2.0 | 3.9 | 9.2 | 3.7 | 5.0 | 1.8 | 6.8 | 7.7 | 9.6 | 2.6 | 4.6 | 5.0 | 3.3 | 2.6 | 8.5 | 5.0 | 7.9 | 2.4 | 2.8 | 457 |
| NaUGT g00526 | 5.9 | 0.6 | 4.9 | 8.2 | 5.5 | 6.3 | 2.3 | 6.8 | 6.3 | 11.0 | 2.1 | 4.2 | 5.5 | 3.6 | 4.4 | 7.8 | 3.6 | 5.5 | 2.1 | 3.2 | 473 |
| NaUGT g00527 | 5.3 | 1.1 | 5.6 | 9.0 | 6.8 | 4.9 | 1.3 | 6.0 | 7.5 | 10.5 | 1.5 | 3.6 | 6.6 | 2.6 | 4.3 | 7.3 | 4.3 | 6.6 | 2.1 | 3.2 | 468 |
| NaUGT g01854 | 5.9 | 1.5 | 4.2 | 8.4 | 4.4 | 6.4 | 2.4 | 5.7 | 6.2 | 11.9 | 3.1 | 4.0 | 6.8 | 2.6 | 4.2 | 6.4 | 4.6 | 7.9 | 1.3 | 2.2 | 455 |
| NaUGT g02083 | 6.4 | 1.5 | 4.4 | 6.6 | 4.4 | 4.6 | 4.6 | 6.8 | 4.8 | 9.8 | 1.9 | 5.0 | 4.8 | 4.1 | 4.6 | 9.3 | 5.4 | 8.1 | 1.5 | 1.7 | 482 |
| NaUGT g02093 | 5.6 | 2.6 | 3.6 | 7.9 | 4.4 | 8.1 | 1.6 | 7.1 | 7.9 | 10.1 | 2.6 | 3.0 | 4.8 | 4.2 | 3.8 | 9.7 | 3.2 | 7.5 | 1.6 | 0.6 | 496 |
| NaUGT g02821 | 5.0 | 1.1 | 2.8 | 6.0 | 5.8 | 5.2 | 3.4 | 7.8 | 5.8 | 8.2 | 2.8 | 6.5 | 5.4 | 3.0 | 5.2 | 11.4 | 4.7 | 6.5 | 1.7 | 1.7 | 464 |
| NaUGT g02942 | 7.3 | 1.1 | 3.9 | 8.8 | 4.9 | 4.7 | 3.4 | 6.2 | 5.6 | 7.7 | 2.1 | 3.6 | 6.9 | 3.6 | 4.7 | 8.4 | 4.1 | 9.2 | 2.1 | 1.7 | 467 |
| NaUGT g03342 | 4.5 | 2.4 | 4.9 | 6.9 | 4.5 | 6.9 | 2.6 | 6.9 | 5.5 | 11.1 | 2.8 | 3.8 | 5.3 | 3.6 | 4.7 | 9.1 | 3.0 | 6.9 | 1.8 | 2.8 | 494 |
| NaUGT g3727 | 5.7 | 1.5 | 4.2 | 5.9 | 6.1 | 4.7 | 4.4 | 6.1 | 7.6 | 8.1 | 3.4 | 4.9 | 5.1 | 4.4 | 3.8 | 8.9 | 3.6 | 8.3 | 1.7 | 1.5 | 472 |
| NaUGT g03748 | 6.5 | 1.5 | 3.0 | 8.8 | 6.3 | 6.0 | 1.9 | 7.3 | 6.7 | 8.0 | 3.2 | 4.3 | 6.0 | 2.6 | 3.9 | 8.6 | 4.5 | 7.1 | 1.9 | 1.7 | 464 |
| NaUGT g04160 | 5.5 | 2.0 | 5.1 | 7.7 | 5.5 | 7.5 | 2.4 | 7.3 | 7.3 | 9.5 | 2.4 | 5.5 | 4.6 | 2.2 | 3.5 | 8.1 | 4.0 | 5.5 | 1.3 | 3.3 | 455 |
| NaUGT g04820 | 6.2 | 1.2 | 5.2 | 8.3 | 5.0 | 6.4 | 1.5 | 8.3 | 7.1 | 9.3 | 3.3 | 4.1 | 4.4 | 3.7 | 3.7 | 8.3 | 4.4 | 5.8 | 1.2 | 2.5 | 482 |
| NaUGT g05060 | 4.5 | 1.5 | 4.0 | 5.3 | 5.9 | 5.3 | 3.6 | 7.2 | 8.3 | 9.1 | 2.8 | 5.5 | 4.5 | 4.2 | 4.0 | 10.4 | 4.0 | 6.6 | 1.7 | 1.5 | 471 |
| NaUGT g05219 | 4.5 | 0.6 | 4.5 | 8.0 | 5.1 | 7.0 | 2.5 | 9.0 | 7.0 | 9.0 | 3.3 | 7.2 | 5.7 | 2.0 | 2.9 | 8.0 | 3.7 | 5.7 | 2.7 | 1.8 | 489 |
| NaUGT g05426 | 5.5 | 2.5 | 4.1 | 10.0 | 4.9 | 6.1 | 2.0 | 7.2 | 7.4 | 10.2 | 3.7 | 4.9 | 4.9 | 3.5 | 3.7 | 6.7 | 3.1 | 6.7 | 2.0 | 0.8 | 489 |
| NaUGT g05573 | 5.1 | 1.5 | 5.3 | 7.0 | 5.7 | 7.0 | 2.1 | 6.4 | 5.3 | 10.6 | 2.1 | 4.9 | 6.1 | 3.6 | 4.4 | 7.2 | 4.0 | 7.2 | 2.3 | 2.1 | 472 |
| NaUGT g06355 | 6.4 | 1.7 | 4.5 | 7.2 | 7.0 | 6.2 | 2.3 | 4.2 | 7.9 | 9.8 | 3.0 | 3.4 | 6.6 | 4.0 | 2.3 | 6.6 | 4.7 | 9.3 | 1.1 | 1.9 | 471 |
| NaUGT g08104 | 4.7 | 1.2 | 3.9 | 7.5 | 6.7 | 6.1 | 2.0 | 7.3 | 7.5 | 9.8 | 3.1 | 8.0 | 5.1 | 2.9 | 3.1 | 8.6 | 3.5 | 5.3 | 2.2 | 1.6 | 510 |
| NaUGT g9179 | 4.4 | 1.1 | 3.4 | 6.9 | 6.1 | 5.3 | 4.8 | 8.4 | 6.5 | 6.9 | 1.7 | 5.3 | 5.1 | 4.6 | 4.0 | 10.3 | 4.6 | 7.6 | 1.7 | 1.3 | 475 |
| NaUGT g09326 | 5.5 | 2.0 | 5.1 | 7.7 | 4.5 | 5.7 | 2.9 | 6.1 | 6.5 | 10.4 | 2.4 | 4.5 | 5.9 | 2.2 | 2.9 | 7.9 | 6.7 | 6.7 | 2.4 | 1.8 | 491 |
| NaUGT g09327 | 5.7 | 2.0 | 5.3 | 7.5 | 4.7 | 5.9 | 2.9 | 5.5 | 6.1 | 10.0 | 2.6 | 4.9 | 6.3 | 2.4 | 2.6 | 7.3 | 6.5 | 7.3 | 2.4 | 1.8 | 491 |
| NaUGT g10741 | 7.8 | 2.0 | 5.4 | 6.8 | 5.0 | 6.5 | 1.5 | 7.6 | 7.8 | 10.0 | 2.4 | 5.2 | 5.2 | 3.3 | 3.1 | 7.4 | 2.0 | 7.0 | 2.8 | 1.1 | 459 |
| NaUGT g11159 | 4.9 | 2.3 | 4.5 | 9.9 | 5.5 | 7.0 | 2.1 | 7.4 | 6.6 | 10.1 | 2.9 | 4.3 | 4.1 | 5.1 | 3.7 | 5.5 | 3.5 | 6.8 | 2.5 | 1.4 | 487 |
| NaUGT g11521 | 5.5 | 2.0 | 4.1 | 8.7 | 3.5 | 5.0 | 1.7 | 7.4 | 8.1 | 9.4 | 2.6 | 4.6 | 5.5 | 3.5 | 2.4 | 8.1 | 4.8 | 7.9 | 2.4 | 2.8 | 458 |
| NaUGT g11522 | 5.5 | 2.0 | 3.9 | 9.2 | 3.7 | 5.0 | 1.8 | 6.8 | 7.7 | 9.6 | 2.6 | 4.6 | 5.0 | 3.3 | 2.6 | 8.5 | 5.0 | 7.9 | 2.4 | 2.8 | 457 |
| NaUGT g11850 | 9.0 | 1.1 | 4.5 | 8.4 | 4.9 | 5.8 | 2.4 | 6.2 | 6.7 | 11.2 | 1.3 | 1.7 | 5.6 | 3.9 | 3.7 | 8.8 | 3.4 | 6.7 | 1.7 | 3.0 | 465 |
| NaUGT g11851 | 7.6 | 1.3 | 5.9 | 7.2 | 5.3 | 5.3 | 3.0 | 7.0 | 5.1 | 10.1 | 2.3 | 2.7 | 5.9 | 3.8 | 2.7 | 8.9 | 4.2 | 6.6 | 1.9 | 3.2 | 473 |
| NaUGT g12088 | 4.7 | 2.0 | 4.9 | 7.3 | 5.1 | 6.9 | 2.6 | 5.3 | 6.7 | 9.3 | 2.4 | 4.3 | 6.9 | 3.4 | 3.2 | 7.9 | 3.4 | 8.3 | 1.8 | 3.4 | 493 |
| NaUGT g13005 | 7.3 | 1.5 | 5.0 | 7.3 | 6.2 | 5.4 | 3.1 | 6.4 | 4.8 | 10.4 | 2.5 | 3.5 | 6.6 | 3.5 | 3.7 | 7.9 | 4.6 | 7.5 | 1.7 | 1.2 | 482 |
| NaUGT g13538 | 6.0 | 1.9 | 3.5 | 7.3 | 5.0 | 6.0 | 1.5 | 7.6 | 8.6 | 8.4 | 3.0 | 5.6 | 5.4 | 2.4 | 2.8 | 9.1 | 4.5 | 5.4 | 2.4 | 3.5 | 463 |
| NaUGT g13945 | 5.2 | 2.1 | 4.9 | 9.0 | 4.1 | 4.7 | 3.2 | 6.4 | 5.2 | 10.3 | 3.2 | 5.6 | 4.3 | 3.4 | 4.5 | 8.4 | 4.7 | 7.3 | 1.5 | 1.9 | 466 |
| NaUGT g15196 | 4.8 | 2.0 | 4.8 | 7.6 | 3.0 | 6.8 | 2.2 | 6.4 | 7.4 | 10.4 | 2.0 | 3.6 | 4.8 | 4.0 | 3.6 | 8.0 | 5.2 | 9.4 | 1.6 | 2.4 | 500 |
| NaUGT g16288 | 6.6 | 1.7 | 4.8 | 4.8 | 6.2 | 6.2 | 2.7 | 6.0 | 5.6 | 10.8 | 2.3 | 6.8 | 6.0 | 4.1 | 1.9 | 7.1 | 4.8 | 7.3 | 1.7 | 2.7 | 482 |
| NaUGT g16298 | 6.0 | 0.9 | 4.5 | 6.8 | 5.8 | 10.7 | 2.1 | 6.2 | 5.3 | 11.8 | 1.5 | 4.3 | 6.0 | 3.2 | 3.8 | 6.0 | 3.0 | 8.5 | 1.9 | 1.7 | 468 |
| NaUGT g16426 | 5.9 | 1.1 | 4.4 | 8.0 | 2.5 | 5.9 | 2.3 | 7.6 | 6.9 | 10.5 | 2.9 | 4.0 | 4.8 | 2.9 | 4.2 | 8.2 | 6.7 | 6.9 | 1.7 | 2.3 | 475 |
| NaUGT g18508 | 4.4 | 1.8 | 5.5 | 5.3 | 5.7 | 6.1 | 2.6 | 8.5 | 6.1 | 7.7 | 2.2 | 3.9 | 6.1 | 3.9 | 3.1 | 9.2 | 5.5 | 7.4 | 2.2 | 2.8 | 457 |
| NaUGT g18870 | 5.7 | 1.2 | 4.1 | 8.3 | 4.7 | 5.9 | 3.3 | 6.7 | 7.5 | 9.6 | 3.9 | 2.6 | 5.3 | 3.7 | 3.0 | 8.3 | 5.5 | 5.9 | 2.0 | 2.8 | 492 |
| NaUGT g19018 | 6.3 | 2.1 | 4.6 | 6.0 | 5.2 | 6.0 | 2.1 | 9.2 | 7.1 | 9.2 | 3.8 | 5.6 | 4.8 | 4.0 | 2.1 | 8.3 | 4.2 | 5.6 | 2.1 | 1.9 | 480 |
| NaUGT g19190 | 5.8 | 2.1 | 5.4 | 7.6 | 4.8 | 6.0 | 2.5 | 5.6 | 6.4 | 10.7 | 2.5 | 4.3 | 6.4 | 2.3 | 3.1 | 8.1 | 6.2 | 6.4 | 2.3 | 1.7 | 484 |
| NaUGT g19344 | 5.3 | 1.9 | 5.5 | 8.7 | 4.4 | 7.0 | 1.9 | 5.7 | 7.6 | 10.4 | 2.3 | 4.7 | 4.7 | 3.4 | 3.2 | 8.7 | 3.6 | 6.4 | 2.1 | 2.5 | 472 |
| NaUGT g19346 | 3.5 | 2.2 | 3.9 | 7.9 | 4.5 | 4.7 | 2.2 | 8.7 | 8.3 | 11.4 | 3.3 | 5.1 | 4.3 | 5.1 | 2.6 | 8.7 | 4.5 | 6.1 | 1.6 | 1.4 | 492 |
| NaUGT g20123 | 6.0 | 1.5 | 5.2 | 7.9 | 4.7 | 5.2 | 3.2 | 7.9 | 4.7 | 9.2 | 3.0 | 4.3 | 5.2 | 3.6 | 4.5 | 8.2 | 4.9 | 6.7 | 1.5 | 2.6 | 466 |
| NaUGT74P5 | 4.5 | 1.1 | 5.6 | 7.9 | 4.9 | 4.7 | 1.6 | 6.7 | 7.9 | 10.3 | 2.7 | 4.7 | 5.2 | 2.2 | 2.2 | 11.2 | 4.0 | 7.6 | 2.5 | 2.2 | 445 |
| NaUGT g20981 | 7.5 | 0.8 | 4.7 | 6.5 | 4.9 | 6.7 | 3.2 | 6.1 | 5.5 | 10.9 | 2.4 | 5.7 | 6.3 | 2.2 | 4.0 | 8.1 | 4.3 | 6.3 | 2.2 | 1.8 | 494 |
| NaUGT g21652 | 4.5 | 2.2 | 5.8 | 7.1 | 6.0 | 6.7 | 1.5 | 6.0 | 6.3 | 10.4 | 2.2 | 6.9 | 4.3 | 2.6 | 3.7 | 7.3 | 5.2 | 7.1 | 2.2 | 1.9 | 463 |
| NaUGT g21654 | 4.7 | 1.9 | 5.0 | 7.8 | 5.2 | 6.9 | 3.0 | 5.8 | 7.1 | 9.3 | 2.8 | 6.5 | 5.0 | 2.2 | 3.4 | 7.1 | 4.1 | 8.2 | 1.7 | 2.4 | 464 |
| NaUGT g21846 | 5.0 | 2.7 | 6.0 | 5.8 | 5.0 | 5.2 | 2.3 | 7.5 | 4.1 | 12.0 | 3.1 | 3.5 | 5.0 | 3.3 | 5.8 | 7.5 | 6.2 | 6.0 | 1.9 | 2.3 | 483 |
| NaUGT g22203 | 5.0 | 1.1 | 3.8 | 9.5 | 6.1 | 5.6 | 2.5 | 6.5 | 8.6 | 9.2 | 4.1 | 5.0 | 5.2 | 2.7 | 2.7 | 7.9 | 3.4 | 7.4 | 1.4 | 2.5 | 444 |
| NaUGT g22204 | 4.5 | 1.7 | 5.4 | 8.8 | 5.1 | 4.1 | 1.9 | 9.0 | 8.8 | 9.9 | 2.6 | 5.8 | 5.6 | 2.6 | 3.4 | 6.9 | 3.2 | 6.9 | 1.7 | 2.4 | 467 |
| NaUGT g22205 | 5.3 | 1.1 | 4.6 | 8.3 | 5.3 | 5.9 | 2.4 | 7.0 | 8.3 | 9.2 | 3.1 | 5.9 | 6.3 | 3.1 | 2.8 | 5.5 | 3.5 | 8.5 | 1.5 | 2.4 | 457 |
| NaUGT g22481 | 4.3 | 3.5 | 5.4 | 5.6 | 5.4 | 5.8 | 2.7 | 7.9 | 7.9 | 10.6 | 2.1 | 6.0 | 5.0 | 2.5 | 3.1 | 8.1 | 4.8 | 6.6 | 1.9 | 1.0 | 483 |
| NaUGT g22983 | 5.1 | 1.5 | 4.2 | 8.3 | 4.5 | 5.7 | 2.1 | 8.1 | 6.8 | 8.5 | 2.1 | 5.3 | 4.2 | 3.2 | 5.1 | 10.0 | 4.2 | 6.8 | 2.1 | 2.1 | 471 |
| NaUGT g23136 | 6.7 | 1.7 | 3.8 | 9.9 | 5.5 | 6.3 | 2.3 | 5.5 | 6.7 | 7.6 | 3.2 | 3.6 | 4.2 | 2.5 | 4.8 | 8.0 | 6.3 | 7.6 | 1.9 | 2.1 | 476 |
| NaUGT g23176 | 6.5 | 1.3 | 4.6 | 6.3 | 4.0 | 7.1 | 2.3 | 5.2 | 6.1 | 12.1 | 2.3 | 4.2 | 7.1 | 3.1 | 3.6 | 7.9 | 4.4 | 8.4 | 1.3 | 2.3 | 478 |
| NaUGT g23995 | 7.5 | 1.8 | 5.3 | 5.9 | 3.7 | 6.6 | 2.0 | 6.8 | 5.9 | 11.7 | 3.1 | 3.5 | 4.8 | 3.1 | 2.9 | 11.5 | 4.0 | 5.3 | 2.4 | 2.2 | 454 |
| NaUGT g24514 | 5.7 | 1.9 | 5.7 | 6.1 | 4.9 | 6.8 | 2.3 | 6.6 | 3.8 | 11.2 | 3.2 | 3.8 | 4.7 | 3.8 | 5.9 | 7.6 | 5.7 | 6.8 | 1.9 | 1.7 | 473 |
| NaUGT g24515 | 3.4 | 2.9 | 4.8 | 7.3 | 4.0 | 6.3 | 1.9 | 7.5 | 5.9 | 10.9 | 3.1 | 4.4 | 6.3 | 3.4 | 4.8 | 7.3 | 3.8 | 8.4 | 1.9 | 1.7 | 477 |
| NaUGT g24585 | 5.5 | 0.9 | 4.4 | 9.4 | 4.2 | 5.0 | 2.4 | 6.4 | 7.9 | 10.3 | 3.1 | 5.3 | 6.6 | 2.6 | 3.9 | 5.9 | 3.7 | 7.7 | 1.5 | 3.3 | 456 |
| NaUGT g24697 | 6.1 | 1.1 | 3.8 | 8.1 | 4.5 | 5.6 | 3.2 | 7.4 | 7.7 | 9.7 | 4.3 | 4.5 | 5.4 | 3.2 | 3.2 | 7.4 | 3.6 | 7.2 | 1.1 | 2.7 | 443 |
| NaUGT g24726 | 5.5 | 1.6 | 4.7 | 5.9 | 4.1 | 10.5 | 4.7 | 7.4 | 4.9 | 9.0 | 3.5 | 2.5 | 8.6 | 4.1 | 3.3 | 8.6 | 2.5 | 4.7 | 1.8 | 2.1 | 512 |
| NaUGT g25918 | 5.2 | 0.9 | 5.2 | 8.0 | 3.9 | 4.5 | 3.7 | 10.0 | 7.4 | 10.2 | 2.8 | 4.1 | 5.6 | 3.9 | 3.0 | 7.6 | 2.8 | 7.1 | 1.5 | 2.6 | 462 |
| NaUGT g26396 | 3.1 | 2.9 | 5.8 | 7.4 | 5.6 | 6.4 | 2.3 | 6.2 | 6.4 | 10.7 | 2.5 | 5.4 | 5.4 | 3.1 | 4.5 | 8.7 | 3.9 | 5.4 | 2.3 | 2.3 | 485 |
| NaUGT91T1 | 9.0 | 0.8 | 4.4 | 6.9 | 4.4 | 6.9 | 2.1 | 6.1 | 6.3 | 11.6 | 1.5 | 3.8 | 8.0 | 2.9 | 4.4 | 6.7 | 2.1 | 7.8 | 2.1 | 2.1 | 476 |
| NaUGT g27096 | 4.8 | 1.5 | 4.4 | 7.5 | 6.7 | 5.6 | 2.1 | 9.2 | 5.9 | 10.5 | 2.7 | 6.3 | 4.2 | 3.1 | 4.2 | 9.0 | 2.5 | 6.3 | 0.8 | 2.7 | 478 |
| NaUGT g27477 | 6.1 | 2.3 | 5.7 | 5.9 | 4.8 | 5.5 | 3.4 | 8.4 | 6.5 | 9.0 | 2.1 | 4.6 | 5.2 | 4.0 | 4.0 | 7.5 | 4.8 | 6.7 | 1.7 | 1.9 | 477 |
| NaUGT g28046 | 5.5 | 1.5 | 5.1 | 7.7 | 5.5 | 6.8 | 2.1 | 5.5 | 4.9 | 11.9 | 3.4 | 4.1 | 6.4 | 2.8 | 4.3 | 7.9 | 3.4 | 7.2 | 1.7 | 2.1 | 469 |
| NaUGT g28047 | 6.2 | 1.5 | 4.5 | 7.2 | 6.0 | 6.8 | 2.8 | 6.4 | 5.5 | 11.5 | 3.6 | 4.3 | 5.8 | 2.8 | 4.1 | 6.8 | 4.1 | 6.6 | 1.5 | 2.1 | 469 |
| NaUGT g28309 | 5.3 | 2.3 | 3.9 | 9.9 | 5.1 | 6.8 | 2.5 | 6.6 | 7.0 | 9.9 | 3.1 | 4.3 | 3.9 | 5.1 | 4.3 | 5.5 | 3.7 | 7.4 | 2.3 | 1.2 | 487 |
| NaUGT g28311 | 4.7 | 2.9 | 3.5 | 8.2 | 3.7 | 6.1 | 2.4 | 7.3 | 6.7 | 10.0 | 3.3 | 5.7 | 5.3 | 3.9 | 4.7 | 6.7 | 3.5 | 8.0 | 2.0 | 1.4 | 490 |
| NaUGT g28331 | 6.1 | 2.4 | 4.9 | 6.9 | 4.7 | 7.3 | 2.2 | 6.1 | 7.1 | 10.1 | 2.8 | 3.7 | 6.5 | 4.5 | 2.0 | 7.5 | 4.5 | 6.3 | 1.8 | 2.6 | 507 |
| NaUGT g28668 | 5.6 | 1.1 | 3.2 | 7.9 | 5.9 | 6.5 | 1.4 | 5.6 | 7.2 | 11.3 | 2.5 | 5.4 | 5.0 | 2.5 | 2.3 | 10.8 | 5.4 | 7.4 | 1.8 | 1.1 | 443 |
| NaUGT g29220 | 4.9 | 2.3 | 5.7 | 5.5 | 4.7 | 4.9 | 3.1 | 8.8 | 5.3 | 9.0 | 1.8 | 6.2 | 5.1 | 3.9 | 3.9 | 9.4 | 4.9 | 5.3 | 2.3 | 2.7 | 487 |
| NaUGT g29262 | 4.7 | 0.9 | 5.6 | 7.5 | 6.0 | 6.9 | 2.6 | 5.4 | 6.0 | 12.2 | 1.5 | 5.6 | 5.8 | 3.0 | 4.7 | 6.7 | 3.9 | 5.8 | 3.0 | 2.4 | 466 |
| NaUGT g29493 | 5.3 | 1.4 | 5.1 | 6.0 | 4.5 | 5.1 | 3.3 | 6.6 | 5.8 | 9.9 | 4.3 | 4.7 | 4.3 | 4.3 | 5.5 | 8.0 | 5.3 | 7.0 | 1.8 | 1.9 | 513 |
| NaUGT g30507 | 5.8 | 2.2 | 4.9 | 6.2 | 4.0 | 6.2 | 1.6 | 9.1 | 7.6 | 9.6 | 3.1 | 5.1 | 5.3 | 2.7 | 3.1 | 6.5 | 5.1 | 6.5 | 2.9 | 2.4 | 449 |
| NaUGT g30575 | 5.7 | 2.2 | 4.2 | 8.4 | 4.0 | 5.7 | 2.6 | 7.1 | 5.5 | 9.1 | 2.6 | 4.2 | 5.7 | 3.3 | 5.7 | 9.3 | 3.8 | 6.8 | 1.8 | 2.2 | 453 |
| NaUGT g30577 | 6.0 | 1.7 | 4.5 | 6.5 | 4.9 | 5.2 | 3.7 | 6.0 | 6.0 | 7.5 | 3.2 | 5.6 | 6.2 | 3.2 | 4.3 | 8.6 | 4.9 | 7.3 | 1.9 | 2.6 | 465 |
| NaUGT g30863 | 5.4 | 1.8 | 5.8 | 5.4 | 5.2 | 5.6 | 4.4 | 7.7 | 5.8 | 9.3 | 1.4 | 5.6 | 4.4 | 3.6 | 3.0 | 7.1 | 5.8 | 6.9 | 2.0 | 3.4 | 496 |
| NaUGT g30898 | 5.8 | 0.6 | 5.0 | 7.2 | 3.8 | 5.8 | 2.0 | 7.2 | 7.0 | 11.0 | 1.8 | 5.6 | 5.2 | 3.4 | 3.0 | 9.0 | 5.2 | 6.0 | 2.0 | 3.0 | 498 |
| NaUGT g31158 | 6.7 | 1.5 | 4.2 | 8.6 | 4.0 | 5.9 | 2.5 | 5.1 | 6.1 | 11.8 | 2.9 | 3.8 | 6.9 | 2.5 | 4.0 | 6.9 | 5.3 | 7.6 | 1.3 | 2.3 | 475 |
| NaUGT g31196 | 5.8 | 1.0 | 4.0 | 8.4 | 2.5 | 5.8 | 2.3 | 7.1 | 6.9 | 11.1 | 2.5 | 3.3 | 5.4 | 2.7 | 4.4 | 7.5 | 7.3 | 7.3 | 1.7 | 2.9 | 479 |
| NaUGT g32016 | 4.6 | 2.7 | 5.0 | 7.5 | 5.0 | 6.4 | 2.5 | 6.0 | 6.4 | 10.8 | 2.9 | 4.2 | 6.2 | 3.1 | 3.5 | 8.1 | 5.6 | 5.0 | 2.5 | 1.9 | 481 |
| NaUGT g32892 | 6.0 | 1.5 | 4.8 | 8.7 | 5.4 | 5.6 | 1.9 | 7.1 | 6.2 | 9.5 | 2.1 | 4.4 | 5.4 | 2.7 | 4.4 | 9.8 | 3.3 | 7.7 | 1.0 | 2.7 | 482 |
| NaUGT g32894 | 5.9 | 1.3 | 4.8 | 8.4 | 5.2 | 5.0 | 1.9 | 6.1 | 6.3 | 8.8 | 3.3 | 4.6 | 6.1 | 2.9 | 3.8 | 10.5 | 4.0 | 7.7 | 1.0 | 2.5 | 478 |
| NaUGT g33134 | 6.9 | 2.1 | 4.7 | 7.3 | 4.1 | 5.8 | 3.0 | 7.5 | 4.9 | 9.7 | 2.8 | 4.3 | 4.5 | 3.2 | 5.8 | 8.6 | 3.6 | 6.7 | 1.5 | 3.0 | 466 |
| NaUGT g34091 | 5.4 | 2.5 | 4.8 | 7.5 | 4.1 | 5.4 | 2.1 | 6.8 | 7.5 | 11.4 | 2.7 | 4.6 | 6.0 | 2.9 | 2.9 | 7.7 | 5.2 | 5.8 | 2.3 | 2.7 | 483 |
| NaUGT g34091 | 5.4 | 2.5 | 4.8 | 7.5 | 4.1 | 5.4 | 2.1 | 6.8 | 7.5 | 11.4 | 2.7 | 4.6 | 6.0 | 2.9 | 2.9 | 7.7 | 5.2 | 5.8 | 2.3 | 2.7 | 483 |
| NaUGT g34877 | 6.8 | 1.9 | 4.3 | 8.5 | 4.1 | 6.4 | 2.7 | 6.4 | 6.4 | 9.3 | 3.7 | 3.7 | 5.4 | 4.5 | 4.1 | 6.8 | 5.6 | 6.0 | 1.9 | 1.6 | 485 |
| NaUGT g34878 | 7.1 | 1.9 | 3.7 | 8.7 | 4.8 | 6.8 | 2.7 | 6.8 | 5.8 | 9.1 | 3.5 | 3.7 | 5.2 | 4.6 | 3.7 | 7.3 | 5.8 | 4.8 | 2.1 | 1.9 | 482 |
| NaUGT g34991 | 5.3 | 1.8 | 5.1 | 8.1 | 7.3 | 7.5 | 1.8 | 9.5 | 8.1 | 10.7 | 2.4 | 5.1 | 5.1 | 3.0 | 2.8 | 5.7 | 3.0 | 4.0 | 2.0 | 1.8 | 495 |
| NaUGT g35131 | 4.3 | 1.7 | 4.3 | 8.7 | 4.1 | 5.0 | 2.0 | 6.7 | 8.3 | 9.6 | 3.0 | 4.8 | 5.0 | 3.9 | 2.2 | 8.9 | 4.6 | 8.0 | 2.4 | 2.4 | 460 |
| NaUGT g35881 | 6.0 | 0.9 | 3.8 | 7.3 | 6.6 | 8.2 | 2.2 | 8.6 | 6.0 | 8.4 | 1.5 | 6.9 | 6.0 | 2.4 | 1.3 | 8.2 | 5.3 | 6.9 | 2.2 | 1.3 | 452 |
| NaUGT g35911 | 6.7 | 1.5 | 4.2 | 8.6 | 4.0 | 5.9 | 2.5 | 5.1 | 6.1 | 11.8 | 2.9 | 3.8 | 6.9 | 2.5 | 4.0 | 6.9 | 5.3 | 7.6 | 1.3 | 2.3 | 475 |
| NaUGT g36017 | 5.9 | 1.8 | 4.1 | 5.7 | 4.9 | 6.1 | 3.7 | 8.1 | 5.9 | 8.3 | 3.9 | 3.3 | 4.5 | 3.3 | 4.1 | 10.4 | 4.9 | 7.3 | 1.6 | 2.4 | 492 |
| NaUGT g36713 | 5.1 | 1.5 | 4.7 | 7.0 | 5.9 | 5.9 | 2.1 | 7.0 | 6.6 | 8.7 | 3.4 | 3.4 | 6.3 | 3.8 | 4.2 | 7.8 | 5.7 | 7.2 | 1.9 | 1.9 | 473 |
| NaUGT g37675 | 6.5 | 1.3 | 4.2 | 6.9 | 3.3 | 6.9 | 2.7 | 5.6 | 5.6 | 12.9 | 2.3 | 4.8 | 7.1 | 3.1 | 3.8 | 7.3 | 5.0 | 6.9 | 1.7 | 2.3 | 480 |
| NaUGT g38211 | 6.0 | 1.4 | 4.6 | 8.9 | 2.8 | 6.4 | 2.2 | 7.8 | 5.8 | 10.7 | 2.2 | 4.2 | 5.4 | 3.2 | 3.2 | 7.8 | 5.2 | 8.0 | 1.4 | 2.4 | 497 |
| NaUGT g38353 | 6.5 | 1.5 | 4.2 | 9.2 | 5.8 | 6.0 | 2.3 | 7.5 | 7.3 | 8.3 | 2.9 | 3.8 | 3.8 | 2.9 | 4.6 | 8.1 | 5.2 | 6.9 | 1.9 | 1.5 | 480 |
| NaUGT g38610 | 4.6 | 1.5 | 5.1 | 7.4 | 4.2 | 4.9 | 2.7 | 5.9 | 4.6 | 10.8 | 3.6 | 4.0 | 7.4 | 4.2 | 3.8 | 8.2 | 4.4 | 8.6 | 1.5 | 2.5 | 474 |
| NaUGT g38855 | 5.7 | 1.6 | 4.8 | 7.8 | 4.6 | 5.3 | 3.7 | 5.3 | 6.7 | 9.9 | 2.3 | 4.6 | 5.7 | 4.1 | 3.4 | 8.0 | 5.3 | 6.7 | 2.1 | 2.5 | 436 |
| NaUGT g39264 | 5.4 | 2.8 | 4.7 | 7.9 | 5.8 | 6.2 | 1.5 | 8.4 | 7.5 | 8.8 | 1.9 | 7.9 | 3.9 | 2.4 | 3.6 | 7.3 | 3.0 | 6.4 | 2.4 | 2.4 | 467 |
| NaUGT g40324 | 6.6 | 1.7 | 4.2 | 8.5 | 5.5 | 6.1 | 1.7 | 7.8 | 5.9 | 9.5 | 2.1 | 4.0 | 5.5 | 2.5 | 5.1 | 8.7 | 3.4 | 6.3 | 1.9 | 3.0 | 473 |
| NaUGT g40325 | 6.2 | 1.2 | 5.6 | 7.9 | 5.0 | 5.8 | 1.7 | 6.6 | 5.6 | 10.8 | 3.3 | 3.3 | 5.8 | 3.1 | 5.6 | 8.3 | 5.0 | 5.8 | 1.2 | 2.1 | 482 |
| NaUGT g40889 | 5.8 | 1.3 | 4.5 | 7.8 | 4.0 | 5.1 | 2.2 | 6.3 | 4.9 | 9.2 | 3.1 | 3.8 | 5.6 | 3.8 | 6.5 | 10.7 | 3.8 | 7.4 | 2.0 | 2.2 | 448 |
| NaUGT g41120 | 4.8 | 2.2 | 4.6 | 7.9 | 3.9 | 5.5 | 4.4 | 6.1 | 5.0 | 10.5 | 2.6 | 4.1 | 4.8 | 3.5 | 5.5 | 9.0 | 4.4 | 7.2 | 2.0 | 2.2 | 458 |
| NaUGT74P3 | 4.9 | 1.8 | 5.1 | 8.0 | 4.7 | 5.1 | 2.0 | 7.3 | 8.5 | 10.2 | 2.7 | 4.2 | 4.7 | 3.1 | 1.1 | 7.8 | 5.1 | 8.9 | 2.4 | 2.2 | 449 |
| NaUGT g41996 | 6.1 | 1.1 | 3.8 | 8.1 | 4.5 | 5.6 | 3.2 | 7.4 | 7.7 | 9.7 | 4.3 | 4.5 | 5.4 | 3.2 | 3.2 | 7.4 | 3.6 | 7.2 | 1.1 | 2.7 | 443 |
| NoUGT-RT1 | 8.4 | 0.8 | 4.4 | 6.5 | 5.3 | 7.8 | 2.5 | 5.0 | 6.3 | 11.1 | 1.5 | 4.4 | 6.7 | 3.2 | 4.0 | 7.4 | 2.1 | 8.6 | 2.1 | 1.9 | 476 |
| NoUGT74P4 | 5.1 | 1.8 | 5.6 | 7.3 | 4.9 | 5.1 | 2.2 | 7.1 | 8.0 | 10.0 | 2.9 | 4.0 | 4.9 | 3.3 | 1.3 | 7.6 | 5.1 | 8.9 | 2.4 | 2.2 | 449 |
| NoUGT-GT2 | 4.3 | 1.3 | 6.3 | 7.8 | 4.7 | 4.7 | 1.6 | 7.2 | 8.1 | 10.5 | 2.7 | 4.0 | 4.7 | 2.5 | 2.2 | 10.5 | 4.5 | 7.6 | 2.5 | 2.2 | 446 |
| Avg. | 5.7 | 1.7 | 4.6 | 7.6 | 4.9 | 6.0 | 2.5 | 6.9 | 6.6 | 10.0 | 2.7 | 4.6 | 5.5 | 3.3 | 3.7 | 8.1 | 4.4 | 6.9 | 1.9 | 2.2 | 474.05 |
