## Supplemental Table 6 for "Specific decorations of 17-hydroxygeranyllinalool diterpene glycosides solve the autotoxicity problem of chemical defense in *Nicotiana attenuata*"

Supplemental Table 6: Characterization and Assignment of novel and known HGL-DTG in *N. obtusifolia*

| Putative ID | ID | RT (min) | MS/MS Fragments | Mol Form | Adduct | Theo. Mass | m/z | Mean (ppm) | Structure | | | metabolite identification level according to Sumner et al. 2007 | Reference |
|---|---|---|---|---|---|---|---|---|---|---|---|---|---|
| | | | | | | | | | $R_1$ | $R_2$ | Malonyl-groups | | |
| DTG 648 | | 24.1 | 451, 433, 325, 289, 289, 271, 231, 215, 203, 201, 189, 179, 177, 175, 163, 163, 161, 149, 147, 145, 135, 127, 123, 121, 119, 109, 107, 95, 85 | $C_{32}H_{54}O_{12}$ | $[M+NH_4]^+$ | 648.3954 | 648.3954 | 0.00 | 2 Hex# | | | 3 | *not described* |
| DTG 734a | | 24.3 | 411, 393, 289, 271, 249, 231, 215, 163, 145, 105 | $C_{35}H_{56}O_{15}$ | $[M+NH_4]^+$ | 734.3954 | 734.3957 | 0.41 | 2 Hex + 1 Ma# | | 1* | 3 | *not described* |
| DTG 734b | | 24.9 | 537, 519, 411, 393, 375, 357, 289, 271, 271, 267, 265, 249, 231, 215, 213, 203, 189, 177, 175, 163, 161, 159, 149, 147, 145, 135, 133, 127, 123, 121, 119, 109, 107, 105, 95, 85 | $C_{35}H_{56}O_{15}$ | $[M+NH_4]^+$ | 734.3973 | 734.3957 | -2.18 | 2 Hex + 1 Ma# | | 1* | 3 | *not described* |
| DTG 734c | | 25.1 | - | $C_{35}H_{56}O_{15}$ | $[M+NH_4]^+$ | 734.3937 | 734.3957 | 2.72 | 2 Hex + 1 Ma# | | 1* | 3 | *not described* |
| DTG 820a | | 25.5 | 519, 515, 497, 479, 461, 443, 339, 333, 315, 291, 289, 273, 271, 265, 249, 231, 215, 213, 203, 201, 189, 189, 177, 175, 163, 161, 149, 147, 145, 135, 127, 123, 121, 119, 109, 107, 105, 95 | $C_{38}H_{58}O_{18}$ | $[M+NH_4]^+$ | 820.3973 | 820.3961 | -1.46 | 2 Hex + 2 Ma# | | 2* | 3 | *not described* |
| DTG 486 | | 27.6 | 289, 271 | $C_{26}H_{44}O_7$ | $[M+Na]^+$ | 491.2979 | 491.2990 | 2.24 | 1 Hex# | | | 3 | *not described* |
| DTG 718 | | 27.5 | 683, 665, 537, 519, 501, 435, 417, 399, 395, 381, 377, 359, 341, 315, 289, 275, 273, 271, 269, 265, 257, 255, 249, 237, 231, 229, 217, 215, 215, 213, 203, 201, 191, 189, 189, 187, 177, 175, 163, 161, 149, 147, 147, 145, 137, 135, 133, 129, 127, 123, 121, 119, 111, 109, 109, 107, 105, 95, 85, 81 | $C_{35}H_{56}O_{14}$ | $[M+NH_4]^+$ | 718.4027 | 718.4008 | -2.64 | 1 Hex + 1 DHex + 1 Ma# | | 1* | 3 | *not described* |
| DTG 572a | | 28.4 | 289, 273, 271, 249, 231, 215, 203, 201, 177, 175, 163, 161, 159, 149, 145, 135, 127, 121, 109, 105, 95 | $C_{29}H_{46}O_{10}$ | $[M+NH_4]^+$ | 572.3470 | 572.3429 | -7.16 | 1 Hex + 1 Ma# | | 1* | 3 | *not described* |
| DTG 572b | | 29 | - | $C_{28}H_{46}O_{10}$ | $[M+Na]^+$ | 577.2960 | 577.2983 | 3.98 | 1 Hex + 1 Ma# | | 1* | 3 | *not described* |
| DTG 632 | | 26.5 | 597, 579, 451, 435, 433, 417, 399, 381, 363, 337, 335, 313, 309, 289, 283, 273, 271, 269, 257, 255, 243, 237, 231, 229, 219, 217, 215, 203, 201, 201, 193, 191, 189, 187, 177, 175, 173, 165, 163, 163, 161, 159, 149, 147, 147, 145, 145, 139, 137, 135, 133, 129, 127, 123, 121, 119, 111, 109, 107, 105, 95, 85, 81 | $C_{32}H_{54}O_{11}$ | $[M+NH_4]^+$ | 632.4019 | 632.4004 | -2.37 | 1Hex - 1DHex# | | | 3 | *not described* |

Putative ID, putative identification of metabolite; RT (min), Retention time of Dionex Ultimate 3000 System coupled to a ESI-QTOF-MS (Impact II), in min; Mol Form, molecular formula of the metabolite; Adduct, Adduct of the metabolite in positive mode; Theo. Mass, theoretical monoisotopic mass calculated for the adduct; *m/z*, measured mass to charge ratio; Mean delta (ppm), deviation between the average of found accurate mass and real accurate mass, in ppm ; Structure, Structure of the metabolite: $R_1$ - residue at the C-17 of the HGL, $R_2$ - residue at the C-3 of the HGL, Hex = Hexose, DHex = Deoxyhexose, Ma = malonylgroup, # position of the sugar decoration at the HGL is unknown, * position of the malonylgroup is unknown
