## Supplemental Table 7 for "Specific decorations of 17-hydroxygeranyllinalool diterpene glycosides solve the autotoxicity problem of chemical defense in *Nicotiana attenuata*"

Supplemental Table 7: Characterization and Assignment of novel and known HGL-DTG in *N. attenuata*

| | | | | | | | | Structure | | | | |
|---|---|---|---|---|---|---|---|---|---|---|---|---|
| Putative ID | RT (min) | MS/MS Fragments | Mol Form | Adduct | Theo. Mass | m/z | Mean (ppm) | $R_1$ | $R_2$ | Malonyl-groups | metabolite identification level according to Sumner et al. 2007 | Reference |
| Attenoside | 18.8 | 939, 921, 759, 633, 613, 597, 579, 489, 471, 451, 435, 433, 417, 327, 325, 309, 289, 273, 271, 215, 177, 163, 163, 149, 147 | $C_{44}H_{74}O_{21}$ | $[M+NH_4]+$ | 956.5061 | 956.5044 | -1.78 | Glc - Glc (2-1) | Glc - Rha (4-1) | | 1 | Jassbi *et al.* (2006) |
| HGL-DTG 1042 | 19.1 | 557, 417, 411, 395, 271, 249 | $C_{47}H_{76}O_{24}$ | $[M+NH_4]+$ | 1042.5065 | 1042.5045 | -1.92 | Glc - Glc - Glc - Rha# | | 1* | 3 | Heiling *et al.* 2016 |
| HGL-DTG 956 | 18.6 | 471, 325, 309, 289, 271 | $C_{44}H_{74}O_{21}$ | $[M+NH_4]+$ | 956.5061 | 956.5061 | 0.00 | Glc - Glc - Glc - Rha# | | | 3 | Heiling *et al.* 2016 |
| Lyciumoside I | 20.8 | 451, 433, 343, 325, 289, 271, 231, 229, 215, 203, 201, 191, 189, 177, 175, 163, 163, 161, 159, 149, 147, 145, 135, 123, 121, 119, 109 | $C_{32}H_{54}O_{12}$ | $[M+NH_4]+$ | 648.3954 | 648.3951 | -0.46 | Glc | Glc | | 1 | Yahara *et al.* (1993) |
| Lyciumoside II | 18.9 | 613, 487, 451, 433, 325, 289, 271, 215, 163, 163, 145 | $C_{38}H_{64}O_{17}$ | $[M+NH_4]+$ | 810.4482 | 810.4489 | 0.86 | Glc - Glc (2-1) | Glc | | 1 | Yahara *et al.* (1993) |
| Lyciumoside IV | 20.4 | 777, 759, 597, 579, 489, 471, 451, 435, 433, 417, 399, 381, 327, 325, 309, 291, 289, 273, 271, 255, 231, 215, 203, 201, 189, 177, 175, 163, 163, 161, 149, 147, 147, 145, 129, 121, 119, 109 | $C_{38}H_{64}O_{16}$ | $[M+NH_4]+$ | 794.4533 | 794.4535 | 0.25 | Glc | Glc - Rha (4-1) | | 1 | Terauchi *et al.* (1998) |
| Nicotianoside Ia | 21.6 | 863, 845, 683, 665, 597, 579, 575, 557, 539, 537, 521, 519, 451, 435, 433, 417, 413, 411, 399, 395, 393, 381, 377, 359, 341, 327, 309, 289, 273, 271, 267, 255, 249, 237, 231, 231, 215, 213, 203, 201, 189, 177, 175, 163, 163, 161, 149, 147, 147, 145, 135, 129, 127, 121, 119, 111, 109, 95 | $C_{41}H_{66}O_{19}$ | $[M+NH_4]+$ | 880.4537 | 880.4531 | -0.68 | Glc - Ma (6-1) | Glc - Rha (4-1) | 1 | 1 | Heiling *et al.* (2010) |
| Nicotianoside Ib | 21.3 | 683, 665, 557, 537, 519, 435, 417, 413, 411, 399, 395, 377, 359, 327, 309, 289, 271, 249, 231, 215, 213, 189, 177, 175, 163, 161, 149, 147, 145, 121, 109 | $C_{41}H_{66}O_{19}$ | $[M+NH_4]+$ | 880.4537 | 880.4515 | -2.50 | Glc | Glc - Rha (4-1) - Ma (6-1) | 1 | 1 | Heiling *et al.* (2010) |
| Nicotianoside Ic | 22.1 | 683, 537, 435, 417, 395, 309, 289, 271, 249, 231, 215, 149 | $C_{41}H_{66}O_{19}$ | $[M+NH_4]+$ | 880.4537 | 880.4534 | -0.34 | Glc | Glc - Rha (4-1) | 1* | 1 | Heiling *et al.* (2010) |
| Nicotianoside IIa | 22.3 | 417, 395, 335, 271 | $C_{44}H_{68}O_{22}$ | $[M+NH_4]+$ | 966.454 | 966.4534 | -0.62 | Glc - Ma (6-1) | Glc - Rha (4-1) - Ma (6-1) | 2 | 3 | Heiling *et al.* 2016 |
| Nicotianoside IIb | 22.4 | 683, 665, 661, 647, 643, 625, 607, 537, 519, 497, 479, 461, 417, 413, 399, 395, 377, 359, 289, 271, 249, 231, 215, 213, 203, 201, 189, 163 | $C_{44}H_{68}O_{22}$ | $[M+NH_4]+$ | 966.454 | 966.4523 | -1.76 | Glc - Ma (6-1) | Glc - Rha (4-1) - Ma (6-1) | 2 | 1 | Heiling *et al.* (2010) |
| Nicotianoside IIc | 22.9 | 683, 665, 643, 537, 519, 497, 479, 435, 417, 413, 399, 395, 377, 359, 289, 271, 249, 231, 215, 163, 149 | $C_{44}H_{68}O_{22}$ | $[M+NH_4]+$ | 966.454 | 966.4529 | -1.14 | Glc - Ma (6-1) | Glc - Rha (4-1) - Ma (6-1) | 2 | 1 | Heiling *et al.* (2010) |
| Nicotianoside IId | 23.5 | 683, 537, 519, 417, 413, 395, 359, 289, 271, 249 | $C_{44}H_{68}O_{22}$ | $[M+NH_4]+$ | 966.454 | 966.4535 | -0.52 | Glc - Ma (6-1) | Glc - Rha (4-1) - Ma (6-1) | 2 | 3 | Heiling *et al.* 2016 |
| Nicotianoside III | 20.1 | 923, 905, 743, 617, 597, 581, 563, 473, 455, 451, 435, 419, 417, 399, 381, 327, 309, 293, 289, 273, 271, 215, 177, 147, 129 | $C_{44}H_{74}O_{20}$ | $[M+NH_4]+$ | 940.5112 | 940.511 | -0.21 | Glc - Rha (4-1) | Glc - Rha (4-1) | | 1 | Heiling *et al.* (2010) |
| Nicotianoside IV | 20.9 | 829, 683, 665, 563, 557, 541, 537, 455, 435, 417, 413, 399, 395, 377, 359, 309, 293, 289, 271, 249, 231, 147 | $C_{47}H_{76}O_{23}$ | $[M+NH_4]+$ | 1026.5116 | 1026.5095 | -2.05 | Glc - Rha (4-1) | Glc - Rha (4-1) | 1* | 1 | Heiling *et al.* (2010) / Jassbi *et al.* (2011) |
| Nicotianoside IXa | 21.7 | 537, 519, 501, 411, 393, 375, 289, 271, 267, 249, 231, 231, 215, 203, 201, 191, 189k, 177, 175, 163, 161, 149, 147, 145, 135, 133, 127, 123, 121, 119, 109, 107, 105, 95 | $C_{35}H_{56}O_{15}$ | $[M+NH_4]+$ | 734.3957 | 734.3957 | 0.00 | Glc | Glc | 1* | 3 | Heiling *et al.* 2016 |
| Nicotianoside IXb | 21.9 | 537, 519, 451, 433, 411, 393, 289, 271, 267, 249, 231, 231, 215, 189, 177, 175, 163, 161, 149, 145, 121, 119, 109, 105 | $C_{35}H_{56}O_{15}$ | $[M+NH_4]+$ | 734.3957 | 734.3956 | -0.14 | Glc | Glc | 1* | 3 | Heiling *et al.* 2016 |
| Nicotianoside IXc | 22.1 | 537, 519, 433, 411, 289, 271, 249, 231, 231, 215, 203, 201, 189, 177, 175, 163, 161, 149, 135, 121, 119 | $C_{35}H_{56}O_{15}$ | $[M+NH_4]+$ | 734.3957 | 734.3953 | -0.54 | Glc | Glc | 1* | 3 | Heiling *et al.* 2016 |
| Nicotianoside IXd | 22.3 | 451, 433, 429, 411, 289, 271, 249, 231, 231, 215, 203, 201, 191, 189, 177, 175, 163, 161, 149, 145, 135, 121, 119, 109, 105 | $C_{35}H_{56}O_{15}$ | $[M+NH_4]+$ | 734.3957 | 734.395 | -0.95 | Glc | Glc | 1* | 3 | Heiling *et al.* 2016 |
| Nicotianoside V | 21.9 | 829, 683, 665, 643, 563, 559, 541, 537, 523, 519, 505, 497, 435, 417, 399, 395, 377, 359, 293, 289, 271, 249, 231, 215, 213 | $C_{50}H_{78}O_{26}$ | $[M+NH_4]+$ | 1112.5120 | 1112.5103 | -1.53 | Glc - Rha (4-1) | Glc - Rha (4-1) | 1* | 1 | Heiling *et al.* (2010) / Jassbi *et al.* (2011) |
| Nicotianoside VIa | 19.3 | 845, 719, 683, 575, 557, 537, 521, 417, 413, 411, 395, 393, 377, 359, 309, 289, 271, 249, 231, 215 | $C_{47}H_{76}O_{24}$ | $[M+NH_4]+$ | 1042.5065 | 1042.5048 | -1.63 | Glc - Glc (2-1) | Glc - Rha (4-1) | 1* | 1 | Heiling *et al.* (2010) |
| Nicotianoside VIb | 19.5 | 759, 597, 575, 557, 471, 435, 417, 413, 411, 395, 325, 309, 289, 271, 249 | $C_{47}H_{76}O_{24}$ | $[M+NH_4]+$ | 1042.5065 | 1042.5053 | -1.15 | Glc - Glc (2-1) | Glc - Rha (4-1) | 1* | 3 | Heiling *et al.* 2016 |
| Nicotianoside VIc | 19.7 | 827, 681, 539, 417, 411, 393, 271, 215 | $C_{47}H_{76}O_{24}$ | $[M+NH_4]+$ | 1042.5065 | 1042.5063 | -0.19 | Glc - Glc (2-1) | Glc - Rha (4-1) | 1* | 3 | Heiling *et al.* 2016 |
| Nicotianoside VIIa | 19.7 | 643, 607, 515, 497, 479, 461, 417, 273, 271, 249 | $C_{50}H_{78}O_{27}$ | $[M+NH_4]+$ | 1128.5069 | 1128.5066 | -0.27 | Glc - Glc (2-1) | Glc - Rha (4-1) | 2* | 1 | Heiling et al. (2010) |
| Nicotianoside VIIb | 19.9 | 845, 805, 683, 665, 659, 643, 625, 607, 575, 557, 537, 521, 519, 497, 479, 435, 417, 413, 411, 399, 395, 393, 377, 359, 289, 271, 249, 231, 215, 213, 177 | $C_{50}H_{78}O_{27}$ | $[M+NH_4]+$ | 1128.5069 | 1128.5048 | -1.86 | Glc - Glc (2-1) | Glc - Rha (4-1) | 2* | 3 | Heiling *et al.* 2016 |
| Nicotianoside VIIc | 20.2 | 659, 557, 497, 417, 411, 395, 289, 271, 249 | $C_{50}H_{78}O_{27}$ | $[M+NH_4]+$ | 1128.5069 | 1128.5065 | -0.35 | Glc - Glc (2-1) | Glc - Rha (4-1) | 2* | 3 | Heiling *et al.* 2016 |
| Nicotianoside VIIIa | 20.5 | 643, 497, 479, 417, 271 | $C_{53}H_{80}O_{30}$ | $[M+NH_4]+$ | 1214.5073 | 1214.508 | 0.58 | Glc - Glc (2-1) | Glc - Rha (4-1) | 3* | 3 | Heiling *et al.* 2016 |
| Nicotianoside VIIIb | 20.8 | 643, 497, 395, 271 | $C_{53}H_{80}O_{30}$ | $[M+NH_4]+$ | 1214.5073 | 1214.5042 | -2.55 | Glc - Glc (2-1) | Glc - Rha (4-1) | 3* | 3 | Heiling *et al.* 2016 |
| Nicotianoside Xa | 22.7 | 537, 519, 501, 497, 479, 461, 443, 357, 289, 271, 249, 231, 215, 213, 103, 201, 189, 177, 175, 163, 161, 149, 145, 135, 127, 109 | $C_{38}H_{58}O_{18}$ | $[M+NH_4]+$ | 820.3961 | 820.3949 | -1.46 | Glc | Glc | 2* | 3 | Heiling *et al.* 2016 |
| Nicotianoside Xb | 23.2 | 537, 519, 497, 479, 461, 289, 271, 249, 231, 215, 189, 177, 175, 163, 161, 149 | $C_{38}H_{58}O_{18}$ | $[M+NH_4]+$ | 820.3961 | 820.3958 | -0.37 | Glc | Glc | 2* | 3 | Heiling *et al.* 2016 |
| Nicotianoside Xc | 23.5 | 537, 519, 497, 479, 461, 289, 271, 249, 231, 215, 189, 177, 175, 163, 161, 149, 145, 135, 127, 121, 109 | $C_{38}H_{58}O_{18}$ | $[M+NH_4]+$ | 820.3961 | 820.3951 | -1.22 | Glc | Glc | 2* | 3 | Heiling *et al.* 2016 |
| Nicotianoside Xd | 23.8 | 537, 497, 335, 289, 271, 249, 231, 215 | $C_{38}H_{58}O_{18}$ | $[M+NH_4]+$ | 820.3961 | 820.3955 | -0.73 | Glc | Glc | 2* | 3 | Heiling *et al.* 2016 |
| Nicotianoside Xe | 24.0 | 537, 497, 289, 271, 249, 215 | $C_{38}H_{58}O_{18}$ | $[M+NH_4]+$ | 820.3961 | 820.3959 | -0.24 | Glc | Glc | 2* | 3 | Heiling *et al.* 2016 |
| Nicotianoside XIa | 19.3 | 537, 411, 325, 289, 271, 249, 215 | $C_{41}H_{66}O_{20}$ | $[M+NH_4]+$ | 896.4486 | 896.4491 | 0.56 | Glc - Glc (2-1) | Glc | 1* | 3 | Heiling *et al.* 2016 |
| Nicotianoside XIb | 19.4 | 699, 573, 537, 519, 411, 393, 375, 289, 271, 267, 265, 249, 231, 215, 189, 177, 163, 149, 145, 105 | $C_{41}H_{66}O_{20}$ | $[M+NH_4]+$ | 896.4486 | 896.4475 | -1.23 | Glc - Glc (2-1) | Glc | 1* | 3 | Heiling *et al.* 2016 |
| Nicotianoside XIc | 19.6 | 613, 591, 573, 451, 433, 429, 411, 393, 325, 289, 271, 267, 249, 231, 215, 177, 163, 163, 161, 149, 145, 121 | $C_{41}H_{66}O_{20}$ | $[M+NH_4]+$ | 896.4486 | 896.4467 | -2.12 | Glc - Glc (2-1) | Glc | 1* | 3 | Heiling *et al.* 2016 |
| Nicotianoside XId | 19.9 | 451, 429, 411, 325, 289, 271, 249, 163 | $C_{41}H_{66}O_{20}$ | $[M+NH_4]+$ | 896.4486 | 896.4483 | -0.33 | Glc - Glc (2-1) | Glc | 1* | 3 | Heiling *et al.* 2016 |
| Nicotianoside XIIa | 20.0 | 515, 497, 479, 461, 333, 271, 249, 231, 215 | $C_{44}H_{68}O_{23}$ | $[M+NH_4]+$ | 982.4490 | 982.4470 | -2.04 | Glc - Glc (2-1) | Glc | 2* | 3 | Heiling *et al.* 2016 |
| Nicotianoside XIIb | 20.2 | 699, 677, 659, 641, 537, 519, 515, 497, 479, 461, 411, 393, 375, 357, 289, 271, 249, 231, 215, 213, 177, 163, 149, 127, 105 | $C_{44}H_{68}O_{23}$ | $[M+NH_4]+$ | 982.4490 | 982.4473 | -1.73 | Glc - Glc (2-1) | Glc | 2* | 3 | Heiling *et al.* 2016 |
| Nicotianoside XIIc | 20.4 | 659, 519, 497, 411, 289, 271, 249 | $C_{44}H_{68}O_{23}$ | $[M+NH_4]+$ | 982.4490 | 982.4481 | -0.92 | Glc - Glc (2-1) | Glc | 2* | 3 | Heiling *et al.* 2016 |
| Nicotianoside XIIIa | 20.5 | 745, 497, 271 | $C_{47}H_{70}O_{26}$ | $[M+NH_4]+$ | 1068.4494 | 1068.4477 | -1.59 | Glc - Glc (2-1) | Glc | 3* | 3 | Heiling *et al.* 2016 |
| Nicotianoside XIIIb | 20.8 | 785, 745, 727, 519, 515, 497, 479, 461, 333, 271, 249, 215 | $C_{47}H_{70}O_{26}$ | $[M+NH_4]+$ | 1068.4494 | 1068.4463 | -2.90 | Glc - Glc (2-1) | Glc | 3* | 3 | Heiling *et al.* 2016 |
| Nicotianoside XIIIc | 20.9 | 497, 479, 271, 249 | $C_{47}H_{70}O_{26}$ | $[M+NH_4]+$ | 1068.4494 | 1068.4469 | -2.34 | Glc - Glc (2-1) | Glc | 3* | 3 | Heiling *et al.* 2016 |
| Nicotianoside XIIId | 21.0 | 745, 497, 479, 461, 271, 249 | $C_{47}H_{70}O_{26}$ | $[M+NH_4]+$ | 1068.4494 | 1068.4472 | -2.06 | Glc - Glc (2-1) | Glc | 3* | 3 | Heiling *et al.* 2016 |
| Nicotianoside XIIIe | 21.3 | 497, 479, 461, 271, 249 | $C_{47}H_{70}O_{26}$ | $[M+NH_4]+$ | 1068.4494 | 1068.4475 | -1.78 | Glc - Glc (2-1) | Glc | 3* | 3 | Heiling *et al.* 2016 |
| Nicotianoside XIIIf | 21.5 | 497, 271 | $C_{47}H_{70}O_{26}$ | $[M+NH_4]+$ | 1068.4494 | 1068.4473 | -1.97 | Glc - Glc (2-1) | Glc | 3* | 3 | Heiling *et al.* 2016 |
| DTG 648 | 23.3 | 613, 451, 433, 325, 289, 289, 271, 231, 229, 215, 203, 201, 191, 189, 179, 177, 175, 163, 163, 161, 159, 149, 147, 145, 135, 133, 127, 123, 121, 119, 109, 107, 95, 85, 81 | $C_{32}H_{54}O_{12}$ | $[M+NH_4]^+$ | 648.3967 | 648.3954 | -2.00 | 2 Hex# | | | 3 | *not described* |
| DTG 734a | 24.5 | 699, 681, 537, 519, 411, 393, 375, 357, 289, 271, 267, 265, 253, 249, 231, 231, 229, 215, 213, 203, 201, 191, 189, 177, 175, 163, 161, 149, 147, 145, 135, 133, 127, 123, 121, 119, 109, 107, 105, 95, 81 | $C_{35}H_{56}O_{15}$ | $[M+NH_4]^+$ | 734.3949 | 734.3957 | 1.09 | 2 Hex + 1 Ma# | | 1* | 3 | *not described* |
| DTG 734b | 24.7 | 537, 411, 289, 271, 249, 231, 215, 177, 163, 149, 145 | $C_{35}H_{56}O_{15}$ | $[M+NH_4]^+$ | 734.3951 | 734.3957 | 0.82 | 2 Hex + 1 Ma# | | 1* | 3 | *not described* |
| DTG 734c | 24.7 | 411, 289, 271, 249, 231, 215, 189, 177, 149, 145 | $C_{35}H_{56}O_{15}$ | $[M+NH_4]^+$ | 734.3945 | 734.3957 | 1.63 | 2 Hex + 1 Ma# | | 1* | 3 | *not described* |
| DTG 820a | 25.3 | 785, 767, 537, 519, 515, 497, 479, 461, 443, 357, 339, 333, 315, 289, 273, 271, 265, 249, 231, 215, 213, 203, 201, 189, 177, 175, 163, 161, 149, 147, 145, 127, 121, 119, 109 | $C_{38}H_{58}O_{18}$ | $[M+NH_4]^+$ | 820.3944 | 820.3961 | 2.07 | 2 Hex + 2 Ma# | | 2* | 3 | *not described* |
| DTG 820b | 25.7 | 785, 537, 519, 497, 479, 461, 443, 357, 289, 271, 265, 249, 231, 215, 213, 177, 163, 161, 149, 147, 145, 127, 121, 109, 105 | $C_{38}H_{58}O_{18}$ | $[M+NH_4]^+$ | 820.3954 | 820.3961 | 0.85 | 2 Hex + 2 Ma# | | 2* | 3 | *not described* |
| DTG 820c | 26 | 497, 289, 271, 249, 231, 215, 145 | $C_{38}H_{58}O_{18}$ | $[M+NH_4]^+$ | 820.3943 | 820.3961 | 2.19 | 2 Hex + 2 Ma# | | 2* | 3 | *not described* |
| DTG 820d | 24.9 | 497, 461, 271, 249, 231 | $C_{38}H_{58}O_{18}$ | $[M+NH_4]^+$ | 820.3963 | 820.3961 | -0.24 | 2 Hex + 2 Ma# | | 2* | 3 | *not described* |
| DTG 486 | 27 | 451, 289, 271, 215, 191, 177, 163, 149, 147, 121, 119 | $C_{26}H_{44}O_7$ | $[M+NH_4]^+$ | 486.3434 | 486.3425 | -1.85 | 1 Hex# | | | 3 | *not described* |
| DTG 718a | 27.1 | 683, 537, 519, 435, 417, 399, 395, 377, 359, 341, 315, 289, 271, 265, 255, 249, 231, 215, 213, 203, 201, 177, 175, 163, 161, 149, 147, 145, 129, 121, 119, 109, 85 | $C_{35}H_{56}O_{14}$ | $[M+NH_4]^+$ | 718.4017 | 718.4008 | -1.25 | 1 Hex + 1 DHex + 1 Ma# | | 1* | 3 | *not described* |
| DTG 718b | 27.4 | 435, 413, 395, 377, 359, 315, 289, 271, 255, 249, 231, 231, 213, 149 | $C_{35}H_{56}O_{14}$ | $[M+NH_4]^+$ | 718.4012 | 718.4008 | -0.56 | 1 Hex + 1 DHex + 1 Ma# | | 1* | 3 | *not described* |
| DTG 572a | 28.1 | 537, 519, 289, 271, 249, 231, 215, 203, 189, 177, 175, 163, 161, 149, 145, 135, 127, 123, 121, 109, 95 | $C_{29}H_{46}O_{10}$ | $[M+NH_4]^+$ | 572.343 | 572.3429 | -0.17 | 1 Hex + 1 Ma# | | 1* | 3 | *not described* |
| RGHGL | 25.9 | 417, 309, 289, 273, 271, 215, 163, 149, 147, 129 | $C_{32}H_{54}O_{11}$ | $[M+NH_4]^+$ | 632.4009 | 632.4004 | -0.79 | OH | Glc - Rha (4-1) | | 1 | Poreddy *et al.* (2015) |
| DTG 632b | 26 | 417, 327, 309, 289, 273, 271, 147, 129 | $C_{32}H_{54}O_{11}$ | $[M+NH_4]^+$ | 632.4005 | 632.4004 | -0.16 | 1 Hex + 1 DHex# | | | 3 | *not described* |

Putative ID, putative identification of metabolite; RT (min), Retention time of Dionex Ultimate 3000 System coupled to a ESI-QTOF-MS (Impact II), in min; Mol Form, molecular formula of the metabolite; Adduct, Adduct of the metabolite in positive mode; Theo. Mass, theoretical monoisotopic mass calculated for the adduct; *m/z*, measured mass to charge ratio; Mean delta (ppm), deviation between the average of found accurate mass and real accurate mass, in ppm ; Structure, Structure of the metabolite: $R_1$ - residue at the C-17 of the HGL, $R_2$ - residue at the C-3 of the HGL, Hex = Hexose, DHex = Deoxyhexose, Pent = Pentose, Ma = malonylgroup, # position of the sugar decoration at the HGL is unknown, * position of the malonylgroup is unknown
