## Supplemental Table 8 for "Specific decorations of 17-hydroxygeranyllinalool diterpene glycosides solve the autotoxicity problem of chemical defense in *Nicotiana attenuata*"

Supplemental Table 8a: Percentages HGL-DTGs in different tissues of *N. attenuata* (all quantities are in percent)

| | Tissue type | | | | | | | | |
|---|---|---|---|---|---|---|---|---|---|
| Compound group | Stem leaf | Buds | Sepals | Ovarium + Nectarium | Style + Stigma | Anthers + Filament | Corolla tube + limb | Unripe Seed Capsules | Seeds |
| Intermediate HGL-DTGs | 0.1 | 0.2 | 0.2 | 0.1 | 0.2 | 0.1 | 0.1 | 0.2 | 20.6 |
| Lyciumoside I and malonylated forms | 3.9 | 2.1 | 0.3 | 3.4 | 0.2 | 0.0 | 0.0 | 0.1 | 5.6 |
| Lyciumoside II and malonylated forms | 2.6 | 5.1 | 10.4 | 12.0 | 1.2 | 0.6 | 0.8 | 38.3 | 19.1 |
| Lyciumoside IV and malonylated forms | 74.9 | 49.0 | 16.9 | 44.2 | 68.7 | 36.3 | 41.9 | 15.9 | 19.1 |
| Nicotianoside III and malonylated forms | 6.8 | 5.2 | 1.4 | 2.6 | 2.9 | 7.7 | 4.2 | 0.4 | 5.6 |
| Attenoside and malonylated forms | 11.6 | 37.4 | 69.7 | 37.2 | 26.6 | 48.6 | 52.3 | 44.9 | 18.9 |
| DTG 956/DTG 1042/DTG 1118/DTG 1204 | 0.2 | 1.0 | 1.0 | 0.5 | 0.3 | 6.6 | 0.6 | 0.2 | 7.5 |
| Rhamnosylated HGL-DTGS | 93.5 | 92.6 | 89.0 | 84.5 | 98.3 | 99.2 | 99.0 | 61.3 | 51.0 |
| Non-rhamnosylated HGL-DTGS | 6.5 | 7.2 | 10.7 | 15.4 | 1.5 | 0.6 | 0.9 | 38.4 | 24.7 |
| Intermediate HGL-DTGs | 0.1 | 0.2 | 0.2 | 0.1 | 0.2 | 0.1 | 0.1 | 0.2 | 20.6 |

Supplemental Table 8b: Percentages HGL-DTGs in different tissues of *N. attenuata* impaired in *UGT91T1* expression (all quantities are in percent)

| | Tissue type | | | | | | | | |
|---|---|---|---|---|---|---|---|---|---|
| Compound group | Stem leaf | Buds | Sepals | Ovarium + Nectarium | Style + Stigma | Anthers + Filament | Corolla tube + limb | Unripe Seed Capsules | Seeds |
| Intermediate HGL-DTGs | 0.1 | 0.2 | 0.0 | 0.1 | 0.0 | 0.0 | 0.0 | 0.1 | 6.9 |
| Lyciumoside I and malonylated forms | 35.3 | 32.1 | 7.4 | 33.6 | 53.5 | 6.8 | 9.2 | 7.2 | 3.6 |
| Lyciumoside II and malonylated forms | 53.3 | 52.9 | 86.6 | 62.0 | 42.7 | 68.9 | 82.6 | 91.2 | 19.1 |
| DTG 972/ DTG 1058 | 0.4 | 0.6 | 0.5 | 0.2 | 0.2 | 10.3 | 1.1 | 0.4 | 0.3 |
| Lyciumoside IV and malonylated forms | 7.2 | 6.1 | 0.4 | 2.4 | 2.7 | 4.1 | 1.9 | 0.3 | 41.9 |
| Nicotianoside III and malonylated forms | 0.0 | 0.0 | 0.0 | 0.0 | 0.0 | 0.0 | 0.0 | 0.0 | 0.3 |
| Attenoside and malonylated forms | 3.7 | 8.1 | 5.1 | 1.7 | 0.9 | 8.9 | 5.2 | 0.8 | 26.5 |
| DTG 956/DTG 1042/DTG 1118/DTG 1204 | 0.0 | 0.1 | 0.0 | 0.0 | 0.0 | 1.0 | 0.0 | 0.0 | 1.5 |
| Rhamnosylated HGL-DTGS | 11.0 | 14.2 | 5.5 | 4.1 | 3.6 | 14.0 | 7.1 | 1.1 | 70.2 |
| Non-rhamnosylated HGL-DTGS | 89.0 | 85.6 | 94.4 | 95.8 | 96.4 | 86.0 | 92.8 | 98.8 | 23.0 |
| Intermediate HGL-DTGs | 0.1 | 0.2 | 0.0 | 0.1 | 0.0 | 0.0 | 0.0 | 0.1 | 6.9 |
