## Supplemental Table 9 for "Specific decorations of 17-hydroxygeranyllinalool diterpene glycosides solve the autotoxicity problem of chemical defense in *Nicotiana attenuata*"

Supplemental Table 9: Primers

Virus-induced gene-silencing (VIGS) in *N. attenuata*

| Name | Gene | Sequence : (5' to 3') |
|---|---|---|
| 00062X | UGT91T1 | GCGGCGCTCGAGCCTTCCAACATAGCACCACTAG |
| 00062B | UGT91T1 | GCGGCGGGATCCGTTTTGGGGGCCCGAAGAAGAC |
| 00211X | UGT74P3 | GCGGCGCTCGAGGTATCATTTGGAAGTTTGGCCAG |
| 00211B | UGT74P3 | GCGGCGGGATCCGCATTAGTTGGTTGATCTGCCC |
| 05859X | UGT74P5 | GCGGCGCTCGAGGGAAGTTTGGCCAGTCTAGGAG |
| 05859B | UGT74P5 | GCGGCGGGATCCGTTGGTTGATCTGTCCACTGAG |
| GLS1-39 | GLS | GCGGCGGTCGACGAATTGGAAGAACTCAAGAGGTGGTCG |
| GLS2-38 | GLS | GCGGCGGGATCCGCGTGTCGAAGGTGGTACAACTTAGC |

Virus-induced gene-silencing (VIGS) in *N. obtusifolia*

| Name | Gene | Sequence : (5' to 3') |
|---|---|---|
| No_000119Sal | UGT74P6 | GCGGCGGTCGACGAAGCATTGAGTTTGGGAGTGC |
| No_000119B | UGT74P6 | GCGGCGGGATCCTGATACTCAGTCTAATGCATTAC |
| No_000582X | UGT74P4 | GCGGCGTGACTCGAGTCTAACCAAGCCAC |
| No_000582BS | UGT74P4 | GCGGCGGGATCCCCCGGGGTCGACGTCGAGTTCCATCCACAATGAG |
| No_003126X | UGT91T1-like | GCGGCGCTCGAGCCTTCTAACTTAGCACCACTAG |
| No_003126B | UGT91T1-like | GCGGCGGGATCCGTTTTGGGGACCCGAAGAAGACCCG |
| pTV000582 _000119: | | |
| Vektor: pTV_000582 x BamHI x SalI (5858 bp) | | |
| Fragment: pTV_000119 x BamHI x SalI (312 bp). | | |

qPCR Primers

| Name | Gene | Sequence : (5' to 3') |
|---|---|---|
| GT2 F | UGT74P5 | GATGTGTTAGAGGAGGTGGT |
| GT2 R | UGT74P5 | TGATGAGACTTGAGCCATATTC |
| GT1V1 F | UGT74P3 | ATGAGACAATACCAGTAGAAGGA |
| GT1V1 R | UGT74P3 | CGACAACTTTCACAGGATATTCA |
| RT1 F | UGT91T1 | AGAACTCTTGCTTGTCAACC |
| RT1 R | UGT91T1 | TTACATTCACTTCCGAAACCC |
| GGPPS F | GGPPS | GATGATCCACACTATGTCCCTC |
| GGPPS R | GGPPS | CTCGCCGTAGACTTTATGGT |
| GLS F | GLS | TCTGGCCCTATGTTTGAGAG |
| GLS R | GLS | CACCCACATTCCATTTCTTGAG |

Recombinant expression

| Name | Gene | Sequence : (5' to 3') |
|---|---|---|
| For | Gateway | GGG GAC AAG TTT GTA CAA AAA AGC AGG CTT C atg …. |
| Rev | Gateway | GGG GAC CAC TTT GTA CAA GAA AGC TGG GTC tc/ta …. |
| RT1_for | UGT91T1 | GGGGACAAGTTTGTACAAAAAAGCAGGCTTC atgaagaaagacagtatccacg |
| RT1_rev | UGT91T1 | GGGGACCACTTTGTACAAGAAAGCTGGGTC tcaaaccttcatctcagcattatttg |
| NobtRT1_for | NoUGT91T1-like | GGGGACAAGTTTGTACAAAAAAGCAGGCTTC atgaagaaagacagtattcacg |
| NobtRT1_rev | NoUGT91T1-like | GGGGACCACTTTGTACAAGAAAGCTGGGTC tcaaaccttctactcaccgttacttc |
| GT1_for | UGT74P3 | GGGGACAAGTTTGTACAAAAAAGCAGGCTTC atggaagaaatcaccagccaaaagaca |
| GT1_rev | UGT74P3 | GGGGACCACTTTGTACAAGAAAGCTGGGTC TCAGTTTAGTGCATTACAAGCTTG |
| GT2_for | UGT74P5 | GGGGACAAGTTTGTACAAAAAAGCAGGCTTC ATGGAAGAAATAACCACCAAATCTC |
| GT2_rev | UGT74P5 | GGGGACCACTTTGTACAAGAAAGCTGGGTC TTACAAGTTTGAAAGAAATTCTTC |
| NobtGT2_for | NoUGT74P6 | GGGGACAAGTTTGTACAAAAAAGCAGGCTTC ATGGAAGAAATAACCACCAAATCTC |
| NobtGT2_rev | NoUGT74P6 | GGGGACCACTTTGTACAAGAAAGCTGGGTC TTACAAGTTTGAAAGAAATTCTTC |
| NobtGT1_for | NoUGT74P4 | GGGGACAAGTTTGTACAAAAAAGCAGGCTTC ATGGAAGAAATCACCAGCCAAAAGAC |
| NobtGT2_for | NoUGT74P4 | GGGGACCACTTTGTACAAGAAAGCTGGGTC TCAGTTTAGTGCATTACAAGCTTG |

Stable lines in *N. attenuata*

| Name | Gene | Sequence : (5' to 3') | Alignment | Fragment length |
|---|---|---|---|---|
| 00062X | UGT91T1 | GCGGCGCTCGAGCCTTCCAACATAGCACCACTAG | 154-175 | 306 |
| 00062B | UGT91T1 | GCGGCGGGATCCGTTTTGGGGGCCCGAAGAAGAC | 460-439 | |
| 00211X | UGT74P3 | GCGGCGCTCGAGGTATCATTTGGAAGTTTGGCCAG | 802-824 | 310 |
| 00211B | UGT74P3 | GCGGCGGGATCCGCATTAGTTGGTTGATCTGCCC | 1112-1091 | |
| 05859X | UGT74P5 | GCGGCGCTCGAGGGAAGTTTGGCCAGTCTAGGAG | 811-832 | 295 |
| 05859B | UGT74P5 | GCGGCGGGATCCGTTGGTTGATCTGTCCACTGAG | 1106-1085 | |
| No_000119Sal | NoUGT74P6 | GCGGCGGTCGACGAAGCATTGAGTTTGGGAGTGC | | 322 |
| No_000119B | NoUGT74P6 | GCGGCGGGATCCTGATACTCAGTCTAATGCATTAC | | |
| No_000582X | UGT74P4 | GCGGCGTGACTCGAGTCTAACCAAGCCAC | | 322 |
| No_000582BS | UGT74P4 | GCGGCGGGATCCCCCGGGGTCGACGTCGAGTTCCATCCACAATGAG | | |
| No_003126X | NoUGT91T1-like | GCGGCGCTCGAGCCTTCTAACTTAGCACCACTAG | 154-175 | 282 |
| No_003126B | NoUGT91T1-like | GCGGCGGGATCCGTTTTGGGGACCCGAAGAAGACCCG | 460-436 | |
| pTV000582 _000119: | | | | |
| Vektor: pTV_000582 x BamHI x SalI (5858 bp) | | | | |
| Fragment: pTV_000119 x BamHI x SalI (312 bp). | | | | |
| GLS1-39 | GLS | GCGGCGGTCGACGAATTGGAAGAACTCAAGAGGTGGTCG | 1499-1525 | 340 |
| GLS2-38 | GLS | GCGGCGGGATCCGCGTGTCGAAGGTGGTACAACTTAGC | 1839-1814 | |
