## Supplemental Figures for "Specific decorations of 17-hydroxygeranyllinalool diterpene glycosides solve the autotoxicity problem of chemical defense in *Nicotiana attenuata*"

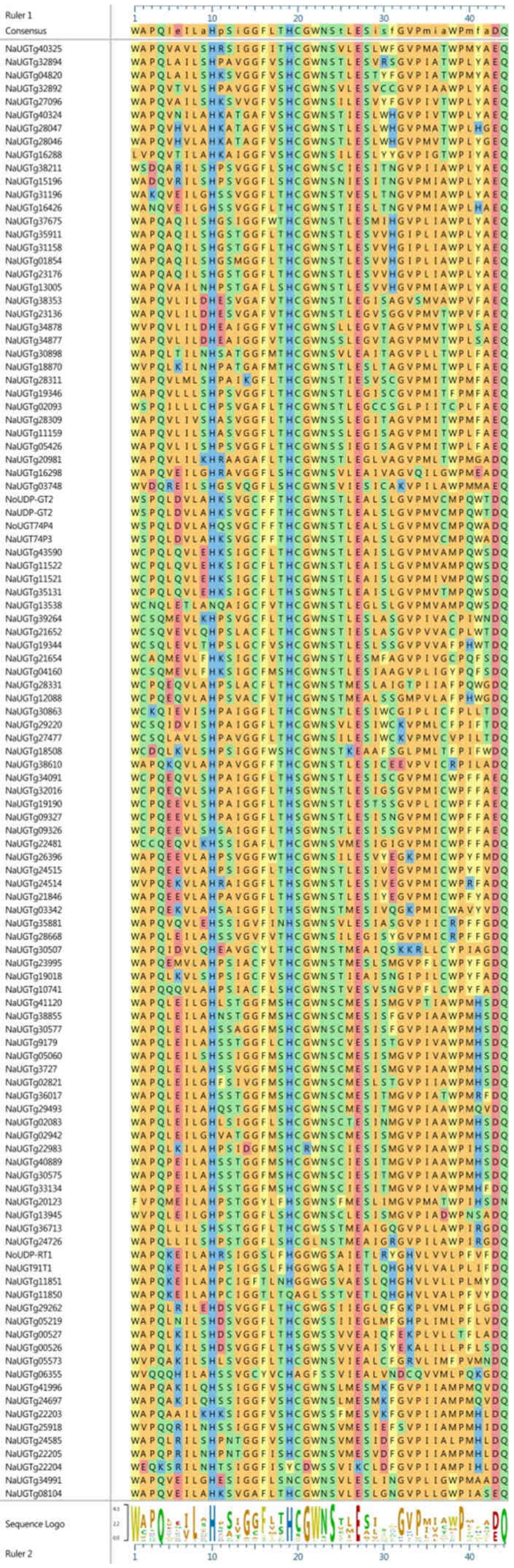
Ruler 1
Consensus
WAPQIeILaHpSiGGFLTHCGWNStLESisfGVPmiaWPmfaDQ
Sequence Logo
Ruler 2

**Supplemental Figure 1: Alignment of the UGT C-terminal consensus sequence of 112 family 1 glycosyltransferases from *N. attenuata* and *N. obtusifolia*.**

To identify members of the UGT family, the 44 amino acids conserved sequence of the PSPG motif was verified using HMMER and aligned using MUSCLE. The consensus sequence is shown in the upper part of the figure and the percentage of similar residues is displayed by the height of the letters in the sequence logo in the lower part of the figure.

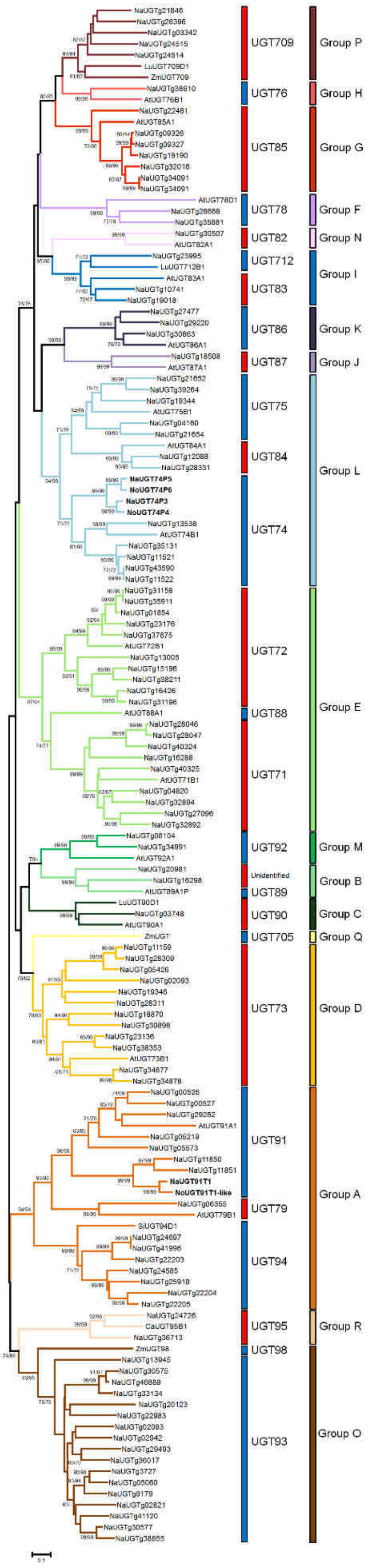


**Supplemental Figure 2: Phylogenetic analysis of the *N. attenuata* UGT superfamily shows 16 major groups**

The phylogenetic tree was obtained by aligning 138 full length amino acid sequences coding for UGTs of the superfamily 1. The phylogenetic relationship was inferred by using either Neighbor-Joining or Maximum Likelihood based on the JTT matrix-based model (both bootstrap = 1000). Bootstrap values over 60% are indicated above the nodes, with the number on the left for the Neighbor-Joining and right for Maximum-Likelihood. 19 *Arabidopsis thaliana*, one *Cicer arietinum*, three *Linum usitatissimum*, one *Sesamum indicum* and three *Zea mays* sequences from each UGT subgroup were included as references in the analysis. Subgroups were created based on sequence similarity (45% similarity for major group, 60% similarity for subgroups) to the reference UGTs and are indicated next to the phylogenetic tree. Sequence similarity to group O was established using ZOG1 and ZOX1 from *Phaseolus vulgaris*. Evolutionary analyses were conducted in MEGA5. All positions containing gaps and missing data were eliminated. NaUGT74P3, NoUGT74P4, NaUGT91T1, NoUGT91T1-like, NaUGT74P5 and NoUGT74P6 analyzed in the present study are in bold letters.

For the construction of the UDP-Glycosyltransferase tree in *N. attenuata*, we used the following Genbank accessions as markers for *Arabidopsis thaliana* (AtUGT71B1 – AB025634; AtUGT72B1 – AC023628; AtUGT73B1 - AT4G34138; AtUGT74B1 - AT1G24100; AtUGT75B1 - AT1G05560; AtUGT76B1 - AT3G11340; AtUGT78D1 - AT1G30530; AtUGT79B1 - AT5G54060; AtUGT82A1 - AT3G22250; AtUGT83A1 - AT3G02100; AtUGT84A1 - AT4G15480; AtUGT85A1 - AT1G22400; AtUGT86A1 - AT2G36970; AtUGT87A1 - AT2G30150; AtUGT88A1 - AT3G16520; AtUGT89A1P – AC006085 (40400-41711); AtUGT90A1 – AC005167 (24146-26230); AtUGT91A1 - AT2G22590; AtUGT92A1 - AT5G12890); *Cicer arietinum* (CaUGT95B1 - gi|533214762); *Linum usitatissimum* (LuUGT709D1 - JN088380; LuUGT712B1 - JN088353; LuUGT90D1 - JN088402); *Sesamum indicum* (SiUGT94D1 - AB333799); *Zea mays* (ZmUGT709 - NP_001148991.1; ZmUGT - NP_001130895.2; ZmUGT98 - NP_001141165.1)

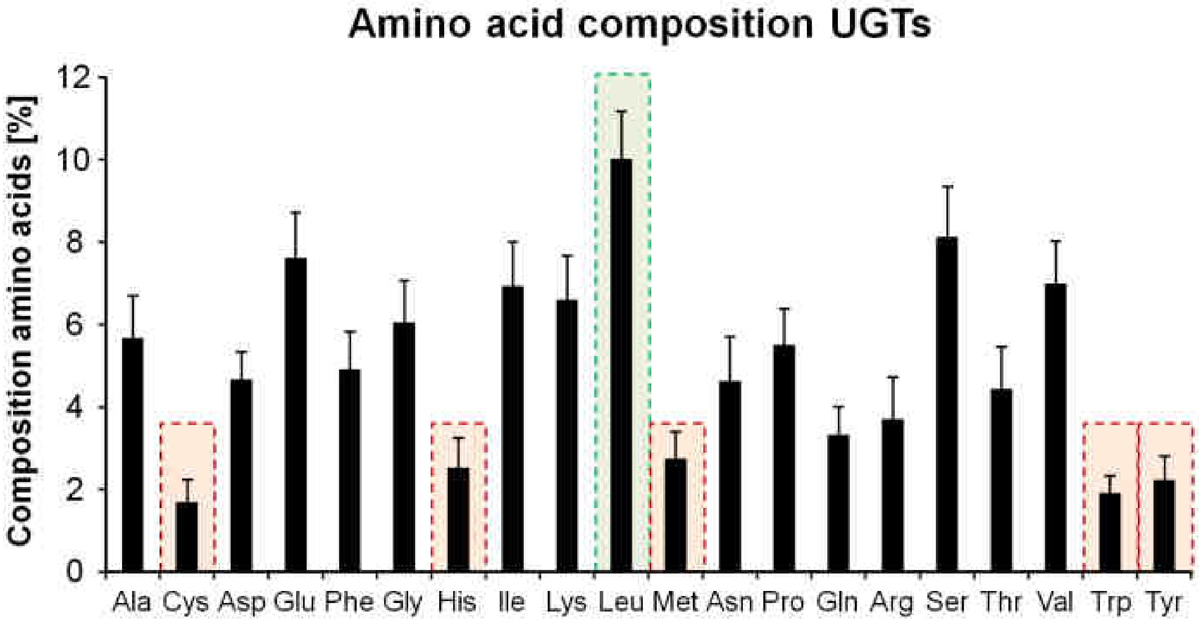


**Supplemental Figure 3: Amino acid composition of all identified UGTs of the superfamily 1 in *N. attenuata***

Shown is the amino acid composition in percentage [%] for all identified UGTs of the superfamily 1 in *N. attenuata*.

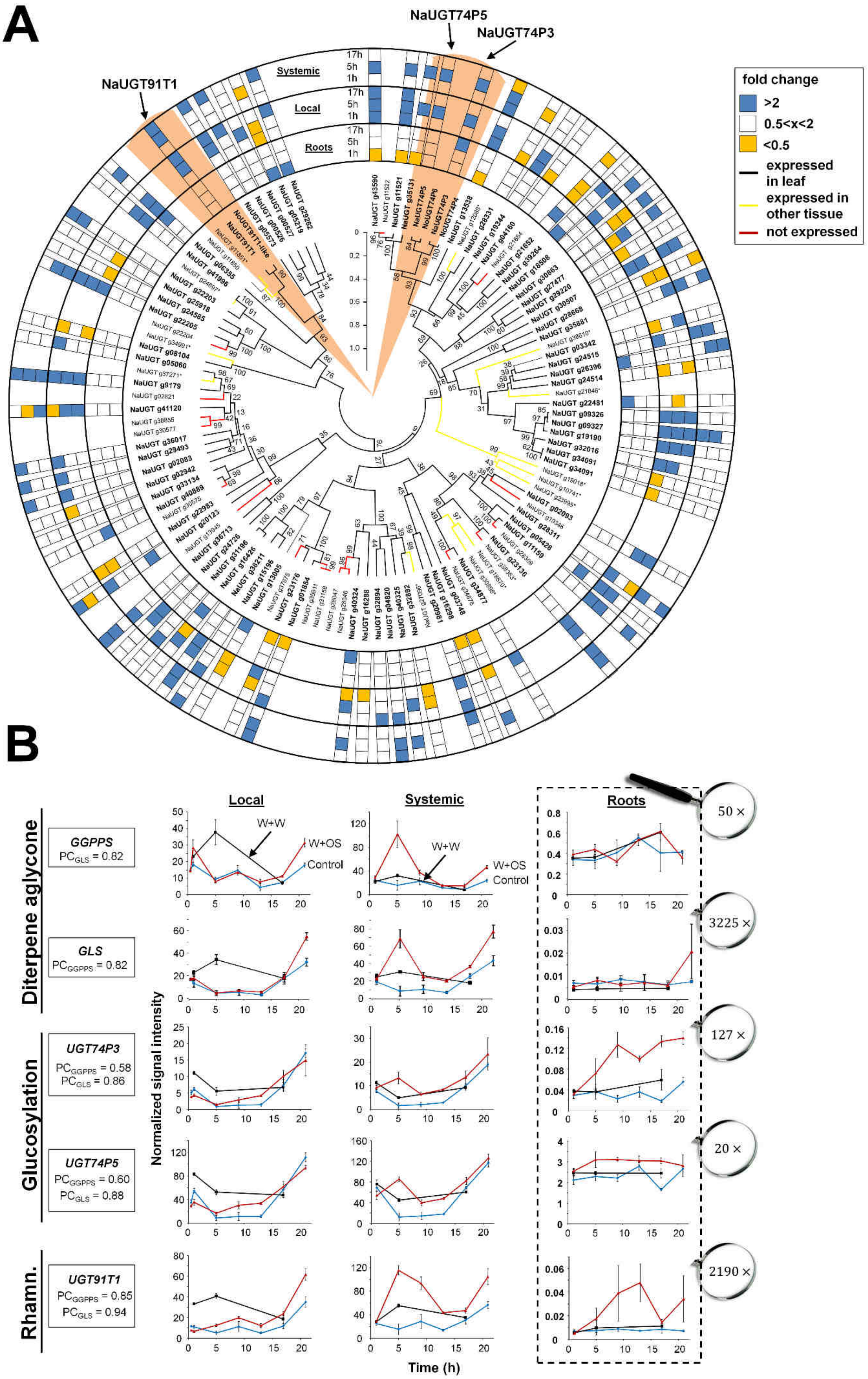
A
NaUGT74P5
NaUGT74P3
NaUGT91T1
Systemic
Local
Roots
17h
5h
1h
fold change
>2
0.5<x<2
<0.5
expressed in leaf
expressed in other tissue
not expressed
B
Local
Systemic
Roots
Diterpene aglycone
Glucosylation
Rhamn.
GGPPS
PC$_{GLS}$ = 0.82
GLS
PC$_{GGPPS}$ = 0.82
UGT74P3
PC$_{GGPPS}$ = 0.58
PC$_{GLS}$ = 0.86
UGT74P5
PC$_{GGPPS}$ = 0.60
PC$_{GLS}$ = 0.88
UGT91T1
PC$_{GGPPS}$ = 0.85
PC$_{GLS}$ = 0.94
W+W
W+OS
Control
Normalized signal intensity
Time (h)
50 ×
3225 ×
127 ×
20 ×
2190 ×

**Supplemental Figure 4: Phylogenetic relationships and herbivory-induced tissue-specific expression of 110 predicted UDP-glycosyltransferases (UGT)**

A) UGT phylogenetic tree generated from the alignment of 107 UGTs from *N. attenuata* and three UGTs from *N. obtusifolia* inferred from the Maximum Likelihood method based on the JTT matrix based model (bootstrap = 1000) (Jones et al. 1982). UGTs were identified based on the presence of the PSPG-Box motif. Shown are the changes in transcript abundance (fold change W+OS vs control: <0.5 and >2) after 1h, 5h and 17h (N=3 biological replicates per time point and per treatment group) resulting from puncture wounds being immediately treated with oral secretions (W+OS) of *M. sexta* larvae in local treated leaves, orthostichous systemic leaves and roots. Expression of 77 UGTs was detected in these tissues, 12 UGTs were detected in other tissue types from an RNAseq tissue atlas and 18 UGTs were not detected in any of the profiled tissues. Light orange sectors highlight the three UGTs characterized in this study for their involvement in HGL-DTG biosynthesis in *N. attenuata* (black). B) Kinetic analysis of W+OS elicited transcript levels of *GGPPS*, *GLS*, *UGT74P3*, *UGT74P5* and *UGT91T1* in locally elicited, systemic leaves and root tissues. Normalized transcript levels were obtained from a previously published full transcriptome microarray experiment (Kim *et al.* 2011). To identify candidate UGTs responsible for HGL-DTG biosynthesis, tissue-level Pearson correlations (PC) were calculated between candidate transcript abundances and those of known (GGPPS, Jassbi *et al.* 2008; GLS, Falara *et al.* 2014) genes in the pathway. *UGT74P3*, *UGT74P5* and *UGT91T1* returned high PC scores with *GLS* in systemic tissues, as did *UGT91T1* with *GGPPS* and *GLS*. Values within the magnifier represent the average magnification factor between root and shoot tissue.

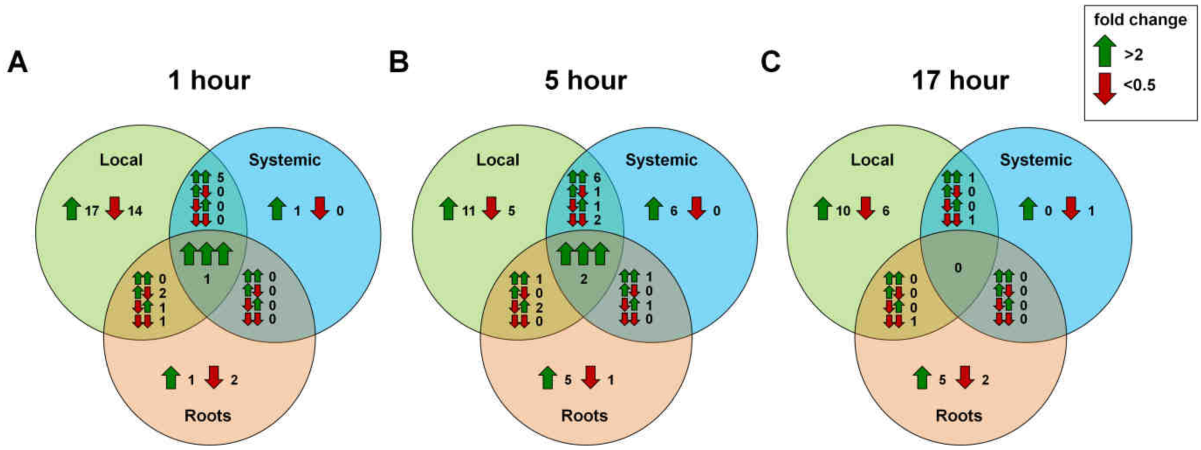


**Supplemental Figure 5: Transcriptomic variation of UGTs after treatment with OS in *N. attenuata***

Venn-diagrams in panel A), B) and C) display the up- and down-regulated UGTs at 1, 5 and 17h after treatment with W+OS in local and systemic leaf tissue as well as in root tissue in *N. attenuata* (N=3). Arrows represent an increase or decrease in the fold change of the expression of UGTs of W+OS-treated vs. Control tissue.

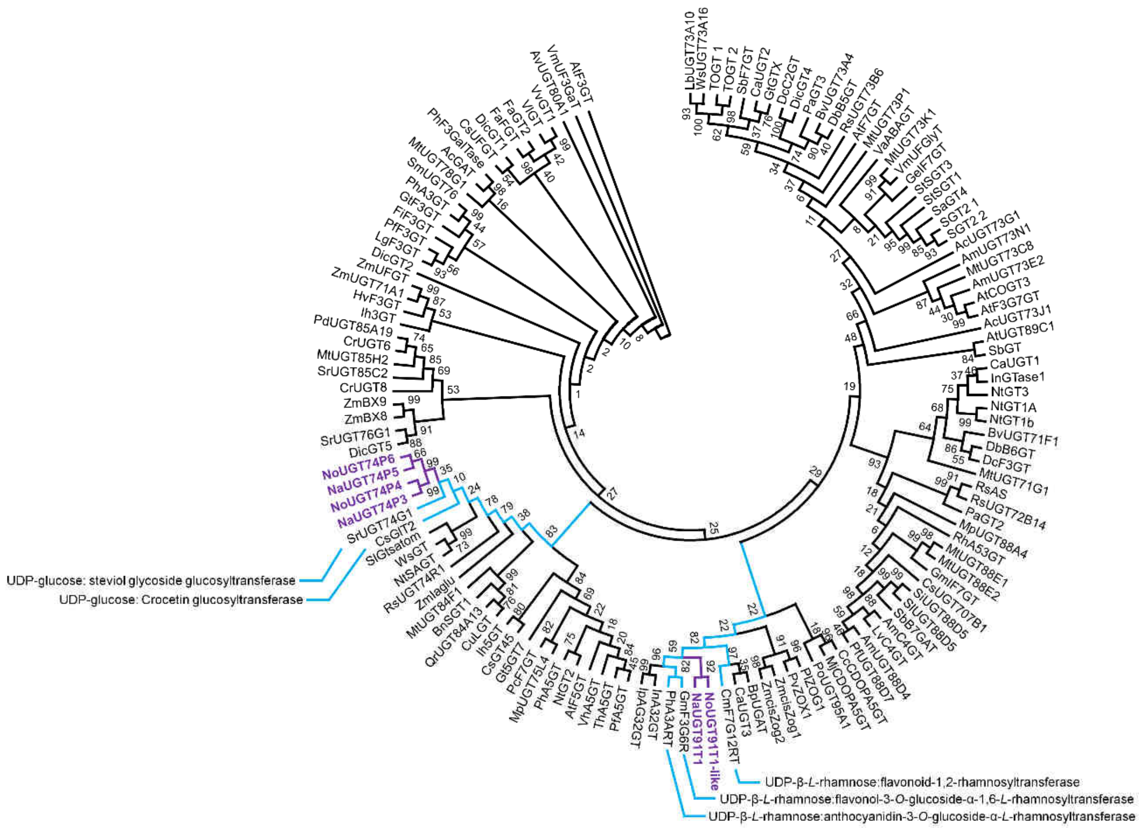


**Supplemental Figure 6: Phylogenetic tree analysis for the UDP-glycosyltransferases used for stable and transient silencing.**

The tree was obtained by aligning characterized glycosyltransferases of the GT superfamily 1 (GT 1, 129 GT amino acid sequences) and inferring their phylogenetic relationship using the Maximum Likelihood method (bootstrap = 1000) based on the JTT matrix-based model. Evolutionary analyses were conducted in MEGA5. NaUGT74P3, NoUGT74P4, NaUGT91T1, NoUGT91T1-like, NaUGT74P5 and NoUGT74P6 analyzed in the present study are highlighted with purple branches. Blue branches indicate closely related UGTs with similar function.

For the construction of the UDP-glycosyltransferase tree in different species, we used the following Genbank accessions for *Allium cepa* (UGT73G1, AAP88406.1; UGT73J1, AAP88407.1), *Antirrhinum majus* (AmC4GT, BAE48239; UGT73E2 (Amugt36), BAG16513.1; UGT73N1 (Amugt38), BAG16514.1; UGT88D4, BAG31945), *Arabidopsis thaliana* (AtF3G7GT, Q9ZQ95; AtF3GT, AAM91339; AtF5GT, AAM91686; AtF7GT, AAL90934; AtUGT89C1, AAP31923; AtDOGT1, NP_181218; AtUGT75D1, AAB58497.1), *Aralia cordata* (AcGAT, BAD06514), *Avena sativa* (AvUGT80A1, CAB06081), *Bellis perennis* (UGT94B1 (BpUGAT), BAD77944), *Beta vulgaris* (BvUGT71F1, AAS94330; BvUGT73A4, AAS94329.1), *Brassica napus* (UGT84A9 (BnSGT1), AF287143_1), *Catharanthus roseus* (CaUGT3, BAH80312; CaUGT1, BAD29721; CaUGT2, BAD29722; UGT85A2a (CrUGT6), BAK55749; UGT709C2 (CrUGT8),BAO01109), *Celosia cristata* (CcCDOPA5GT, BAD91804), *Citrus maxima* (CmF7G12RT, AAL06646), *Citrus sinensis* (CsUFGT, AAS00612), *Citrus unshiu* (CuLGT, BAA93039), *Crocosmia x crocosmiiflora (*CcUGT77B2,

MG938542); *Crocus sativus* (CsGT45, ACM66950.1; CsUGT707B1, CCG85331; Glt2 (UGTCs2), AAP94878.1), *Dianthus caryophyllus* (DcF3GT, BAD52004; DicGT1, BAD52003; DicGT2, BAD52005; DicGT4 (DcC2GT), BAD52006; DicGT5, BAD52007), *Dorotheanthus bellidiformis* (DbB5GT, CAB56231; DbB6GT, AAL57240), *Forsythia x intermedia* (FiF3GT, AAD21086), *Fragaria x ananassa* (FaFGT, AAU12367; FaGT2, AAU09443), *Gentiana triflora* (Gt5GT7, BAG32255; GtF3GT, BAA12737; GtGTX, BAC54092), *Glycine max* (GmF3G6R, BAN91401; GmIF7GT, BAF64416), *Glycyrrhiza echinata* (GeIF7GT, BAC78438), *Hordeum vulgare subsp. vulgar* (HvF3GT, CAA33729), *Ipomoea nil* (In3GGT(InA32GT), BAD95885; InGTase1, BAF75917), *Ipomoea purpurea* (Ip3GGT(IpA32GT), BAD95882), *Iris x hollandica* (Ih3GT, BAD83701; Ih5GT, BAD06874), *Lamium galeobdolon* (LgF3GT, AEB61487), *Linaria vulgaris* (LvC4GT, BAE48240), *Lycium barbarum* (Ugt73a10, BAG80536), *Maclura pomifera* (MpUGT75L4, ABL85474; MpUGT88A4, ABL85471), *Medicago trunculata* (MtUGT73C8, ABI94020; MtUGT73K1, AAW56091; MtUGT73P1, ABI94026; MtUGT78G1, ABI94025; MtUGT84F1, ABI94023; MtUGT85H2, ABI94024.1; MtUGT88E1, ABI94021; MtUGT88E2, ABI94025; MtUGT71G1, AAW56092), *Mirabilis jalapa* (CDOPA5GT, BAD91804), *Nicotiana tabacum* (NTGT1A, BAB60720.1; NTGT1b, BAB60721.1; NtGT2, BAB88935; NtGT3, BAB88934; NtSAGT, AAF61647; TOGT 1, AAK28303; TOGT 2, AAK28304), *Perilla frutescens* (PfA5GT, BAA36421; PfF3GT, BAA19659; PfUGT88D7 (F7GAT), BAG31948), *Petunia x hybrida* (PhA3ART, CAA50376; PhA3GT, BAA89008; PhA5GT, BAA89009; PhF3GalTase, AF165148_1), *Phaseolus lunatus* (PlZOG1, AAD04166); *Phaseolus vulgaris* (PvZOX1, AF116858_1), *Phytolacca americana* (PaGT2, BAG71125; PaGT3, BAG71127), *Pilosella officinarum* (PoUGT95A1, ACB56927), *Prunus dulcis* (PdUGT85A19, ABV68925), *Pyrus communis* (PcF7GT, AAY27090), *Quercus robur* (QrUGT84A13, AHA54051), *Rauwolfia serpentine* (RsAS, CAC35167), *Rhodiola sachalinensis* (RsUGT73B6, AAS55083; RsUGT74R1, ABP49574; RsUGT72B14, ACD87062), *Rosa hybrida* (RhA53GT, BAD99560), *Scutellaria baicalensis* (SbB7GAT, BAD99560; SbF7GT, BAA83484), *Scutellaria laeteviolacea var. yakusimensis* (SlUGT88D5, BAG31946), *Sesamum indicum* (SiUGT88D6, BAG31947), *Solanum aculeatissimum* (SaGT4, BAD89042.1), *Solanum berthaultii* (SbGT, AAB62270.1), *Solanum lycopersicum* (SlGtsatom, CAI62049.1), *Solanum melongena* (SmUGT76, CAA54558.1), *Solanum tuberosum* (Sgt2.1, ABB29873.1; Sgt2.2, ABB29874.1; StSgt1, AAB48444.1; StSgt3, ABB84472.1), *Stevia rebaudiana* (SrUGT74G1, AY345982; SrUGT76G1, AY345974; SrUGT85C2, AY345978), *Torenia hybrida* (ThA5GT, BAC54093), *Verbena hybrida* (VhA5GT, BAA36423), *Vigna angularis* (VaABAGT, BAB83692), Vigna mungo (VmUF3GaT, BAA36972; VmUFGlyT, BAA36410), *Vitis labrusca* (VlGT, ABR24135), *Vitis vinifera* (VvGT1, AAB81682), *Withania somnifera* (WsPGT, FJ560880; WsUGT73A16, FJ654696/ACO44747.1), *Zea mays* (ZmBX8, AF331854_1; ZmBX9, CAX02221; ZmcisZog1, AAK53551; ZmcisZog2, AAL92460; ZmIaglu, AAA59054;

ZmUFGT, CAA30760; ZmUGT71A1, CAA31856).

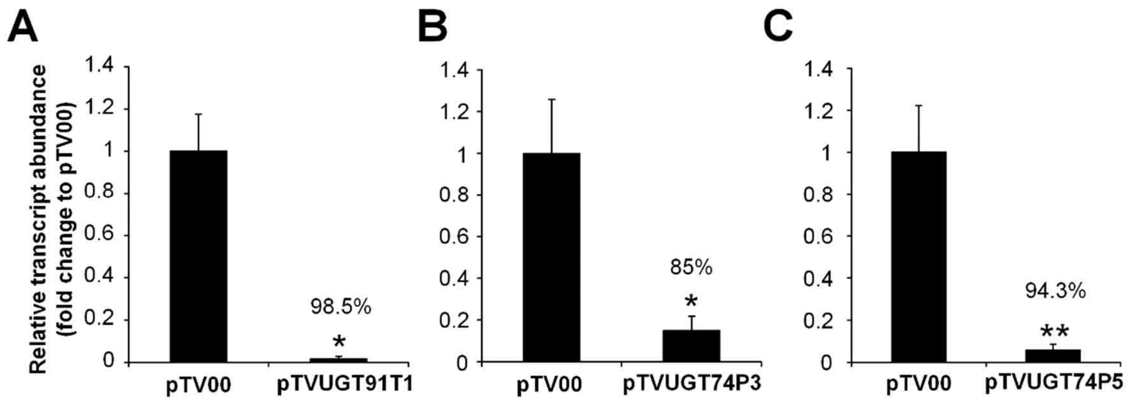


**Supplemental Figure 7: Silencing efficiency for the three transiently-silenced 17-HGL-DTG biosynthetic UGTs in pTVUGT91T1, pTVUGT74P3 and pTVUGT74P5.**

Relative transcript abundance (fold change to pTV00 empty vector *N. attenuata* plants after normalization to the transcript abundance of *Elongation Factor 1α* – Na*ELF1α*) of **A)** *UGT91T1*, **B)** *UGT74P3* and **C)** *UGT74P5* in leaves of transiently-silenced *N. attenuata* plants (N=4). Asterisks indicate significant differences between empty vector control (pTV00) and transiently-silenced lines (*t*-test, *P ≤ 0.05, ** P < 0.01).

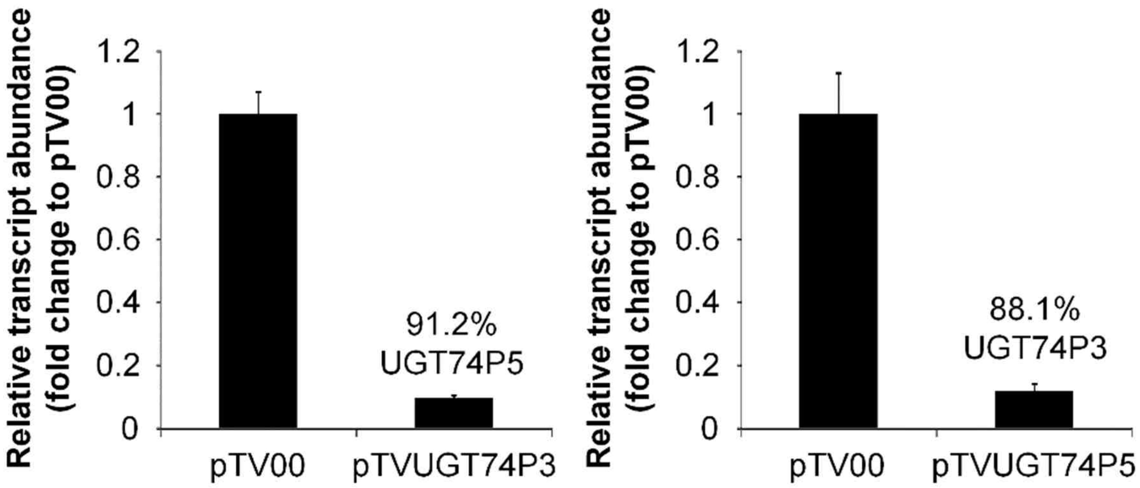


**Supplemental Figure 8: Co-silencing efficiency of UGT74P3 and UGT74P5 in pTVUGT74P3 and pTVUGT74P5.**

Relative transcript abundance (fold change to pTV00 empty vector *N. attenuata* plants after normalization to the transcript abundance of *Elongation Factor 1α* – Na*ELF1α*) of **A)** *UGT74P5* in pTVUGT74P3, **B)** *UGT74P3* in pTVUGT74P5 - in buds of transiently-silenced *N. attenuata* plants (N=7). Asterisks indicate significant differences between empty vector control (pTV00) and transiently-silenced lines (*t*-test, *** $P < 0.001$).

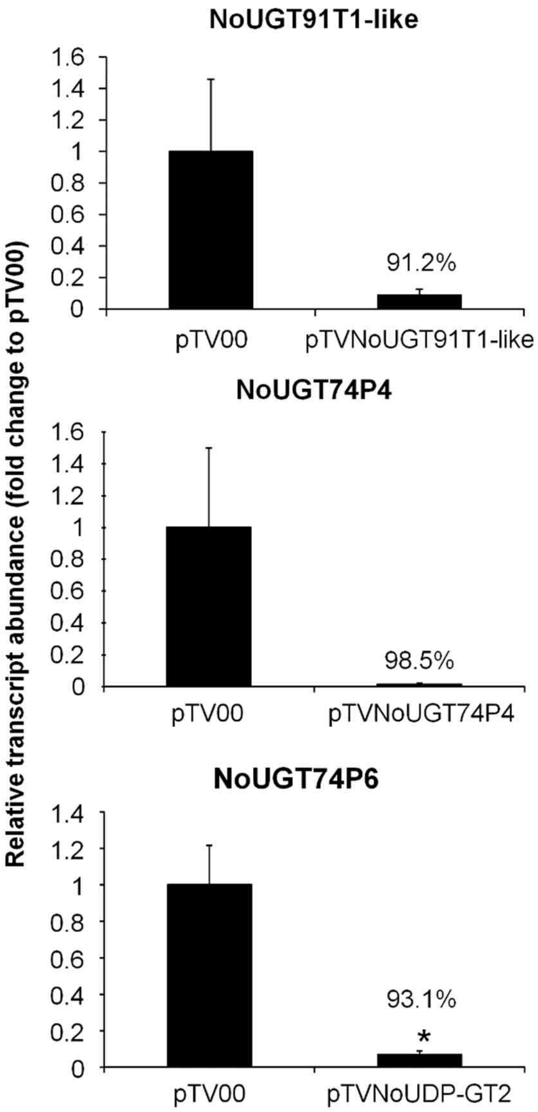


**Supplemental Figure 9: Silencing efficiency for the three transiently-silenced 17-HGL-DTG biosynthetic UGTs in pTVUGT91T1-like, pTVUGT74P3 and pTVUGT74P6 in *N. obtusifolia*.**

Relative transcript abundance (fold change to pTV00 empty vector *N. obtusifolia* plants after normalization to the transcript abundance of *Elongation Factor 1α* – No*ELF1α*) of **A)** No*UGT91T1-like*, **B)** No*UGT74P4* and **C)** No*UGT74P6* in leaves of 37-days-old elongated transiently-silenced *N. obtusifolia* plants (N=4). Asterisks indicate significant differences between empty vector control (pTV00) and transiently-silenced lines (*t*-test, *P ≤ 0.05).

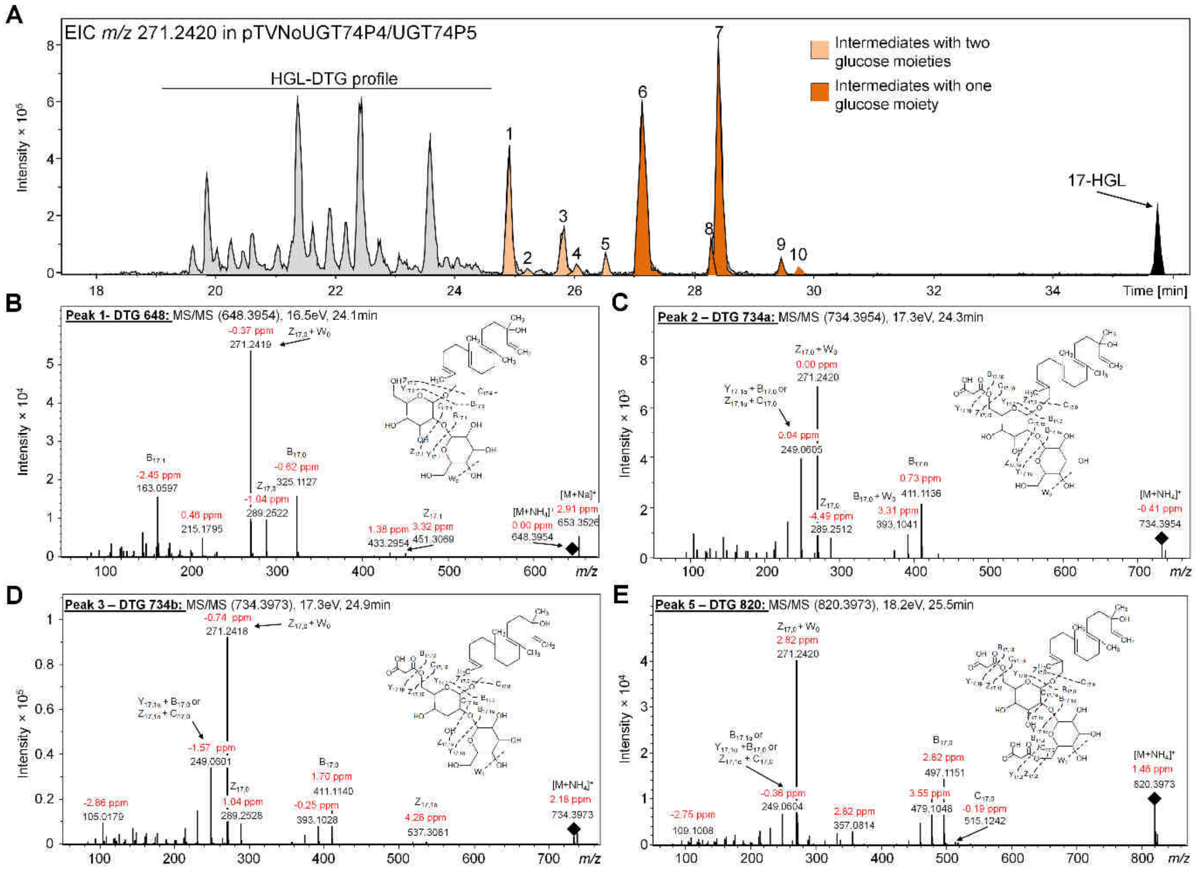

**Supplemental Figure 10a: Mass spectrometric characterization and annotation of novel HGL-DTGs in transiently-silenced *N. obtusifolia* plants impaired in No*UGT74P4* and No*UGT74P6* expression.**

A) Shown is the EIC trace *m/z* 271.2420 for the 17-HGL aglycone fragment representing the metabolic alteration of the HGL-DTG profile in transiently-silenced *N. obtusifolia* plants inoculated with *A. tumefaciens* harboring the vector pTVNoUGT74P4/UGT74P6. Additional novel HGL-DTGs are highlighted in light orange for intermediates containing at least two hexoses (presumably glucose) and in darker orange for intermediates containing only one hexose moiety. Furthermore, the associated MS/MS spectra for the highlighted Peak 1 (B), Peak 2 (C), Peak 3 (D) and Peak 5 (E) are displayed. A putative structure is assigned to all intermediate HGL-DTGs and the fragmentation scheme is explained via the predicted neutral losses from the molecular ion $[M+H]^+$.

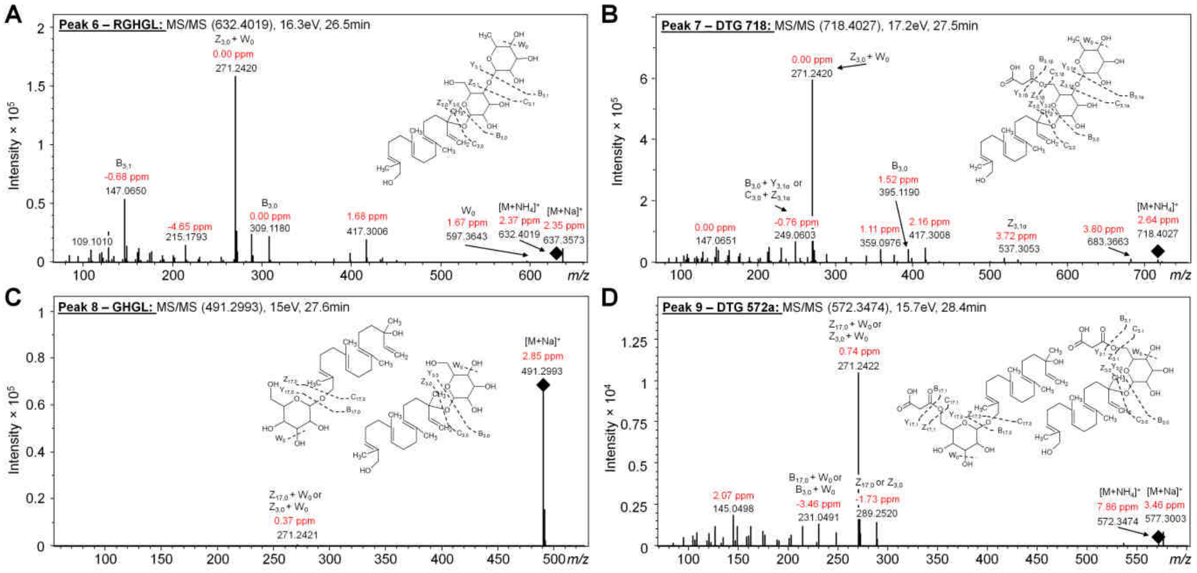


### Supplemental Figure 10b: Mass spectrometric characterization and annotation of novel HGL-DTGs in transiently-silenced *N. obtusifolia* plants impaired in No*UGT74P4* and No*UGT74P6* expression.

MS/MS spectra for the highlighted Peak 6 (A), Peak 7 (B), Peak 8 (C) and Peak 9 (D) are displayed. A putative structure is assigned to all intermediate HGL-DTGs and the fragmentation scheme is explained via the predicted neutral losses from the molecular ion $[M+H]^+$.

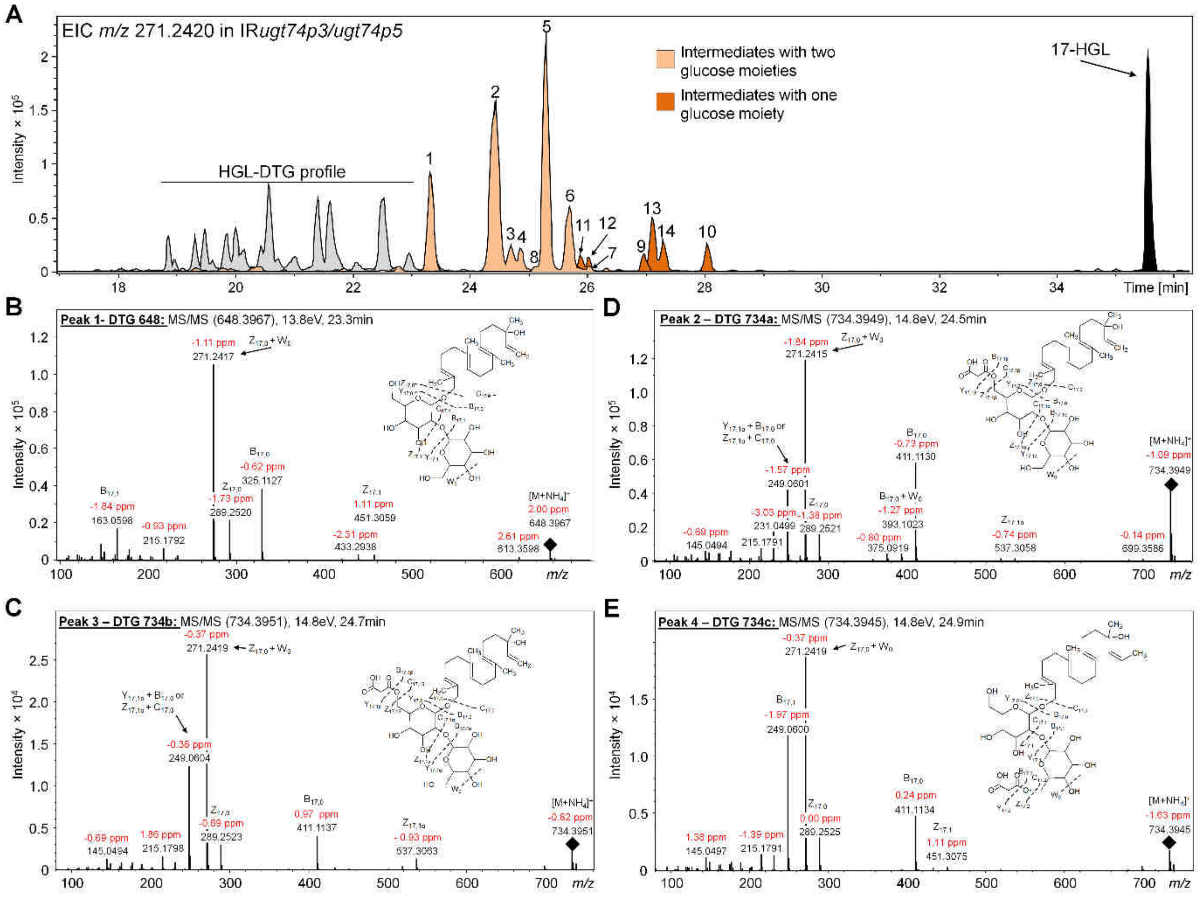


**Supplemental Figure 11a: Characterization and annotation of novel HGL-DTGs via MS/MS in stably-silenced *N. attenuata* plants impaired in *UGT74P3* and *UGT74P5* expression.**

A) Shown is the EIC trace *m/z* 271.2420 for the 17-HGL aglycone fragment representing the metabolic alteration of the HGL-DTG profile in stably-silenced *N. attenuata* plants impaired in *UGT74P3* and *UGT74P5* expression. Additional novel HGL-DTGs are highlighted in light orange for intermediates containing at least two hexoses (presumably glucose) and in darker orange for intermediates containing only one hexose moiety. Furthermore, the associated MS/MS spectra for the highlighted Peak 1 (B), Peak 2 (D), Peak 3 (C) and Peak 4 (E) are displayed. A putative structure is assigned to all intermediate HGL-DTGs and the fragmentation scheme is explained via the predicted neutral losses from the molecular ion $[M+H]^+$.

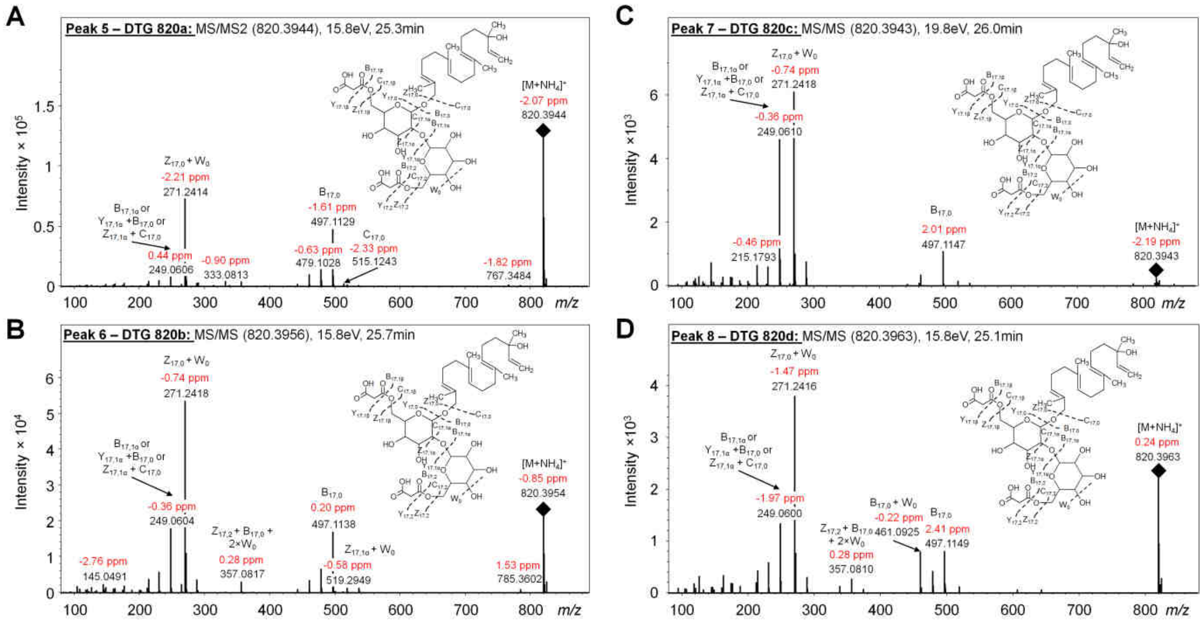


**Supplemental Figure 11b: Characterization and annotation of novel HGL-DTGs via MS/MS in stably-silenced *N. attenuata* plants impaired in *UGT74P3* and *UGT74P5* expression.**

MS/MS spectra for the highlighted Peak 5 (A), Peak 6 (B), Peak 7 (C) and Peak 8 (D) are displayed. A putative structure is assigned to all intermediate HGL-DTGs and the fragmentation scheme is explained via the predicted neutral losses from the molecular ion $[M+H]^+$.

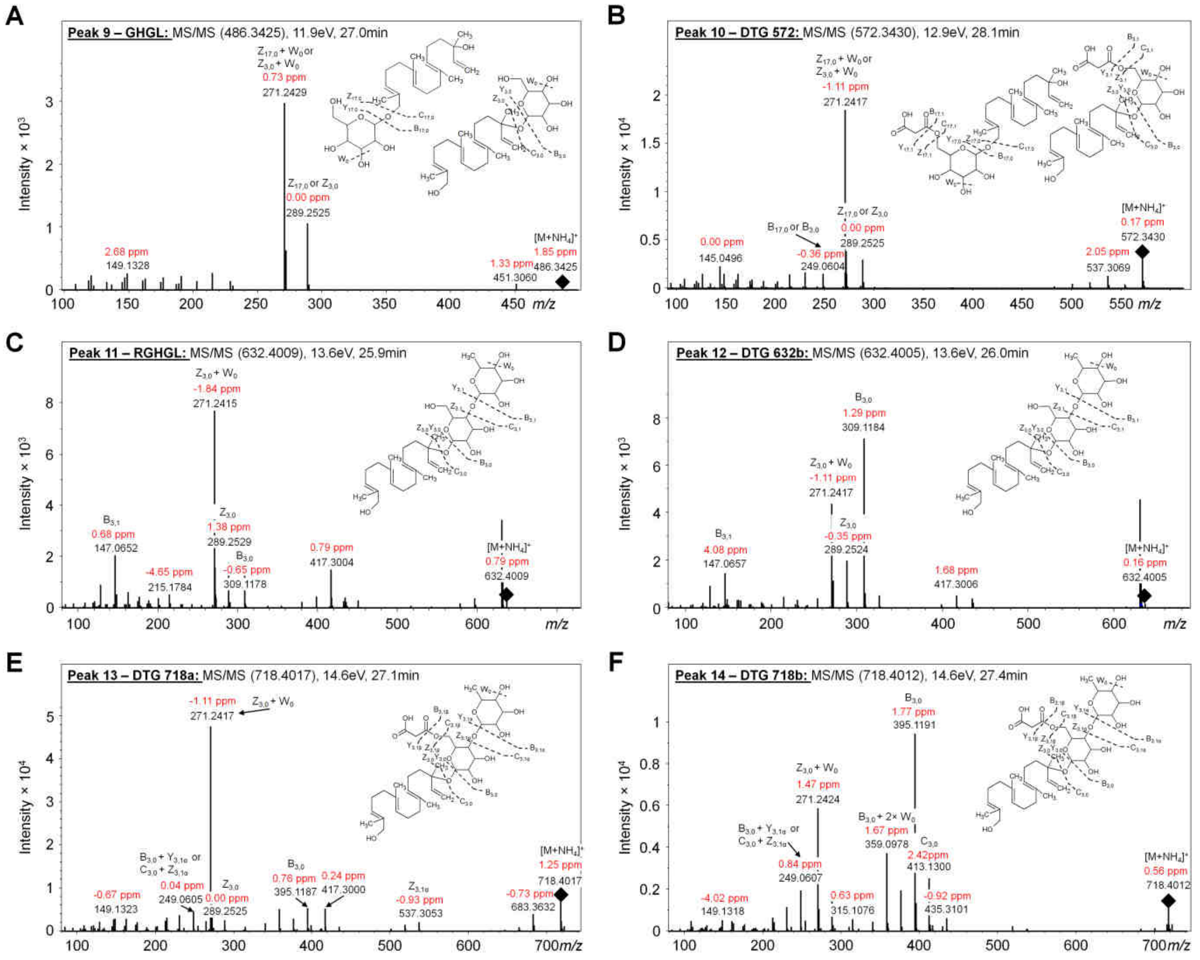


**Supplemental Figure 11c: Characterization and annotation of novel HGL-DTGs via MS/MS in transiently-silenced *N. attenuata* plants impaired in *UGT74P3* and *UGT74P5* expression.**

MS/MS spectra for the highlighted Peak 9 (A), Peak 10 (B), Peak 11 (C), Peak 12 (D), Peak 13 (E) and Peak 14 (F) are displayed. A putative structure is assigned to all intermediate HGL-DTGs and the fragmentation scheme is explained via the predicted neutral losses from the molecular ion $[M+H]^+$.

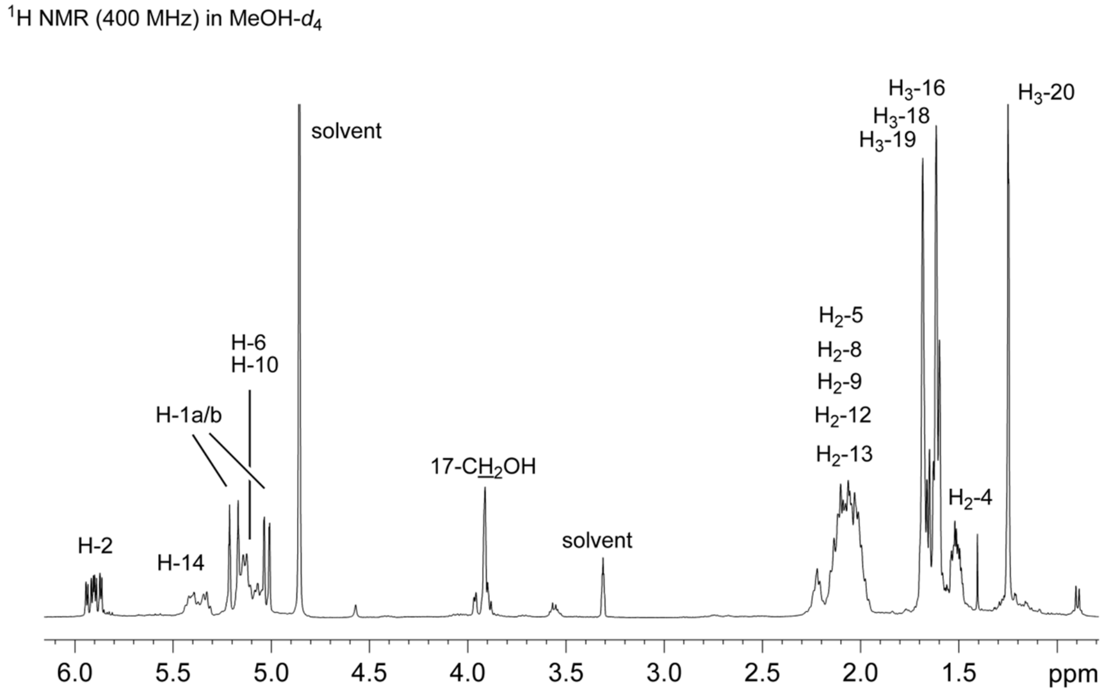


**Supplemental Figure 12:** $^{1}$H NMR spectrum of synthetic 17-hydroxygeranyllinalool (17-HGL, HPC24 Standards)

17-HGL was measured at 400 MHz in MeOH-*d*$_4$ (300 K). Signals were assigned using data from DEPT 135, $^{1}$H-$^{1}$H COSY, HSQC and HMBC spectra.

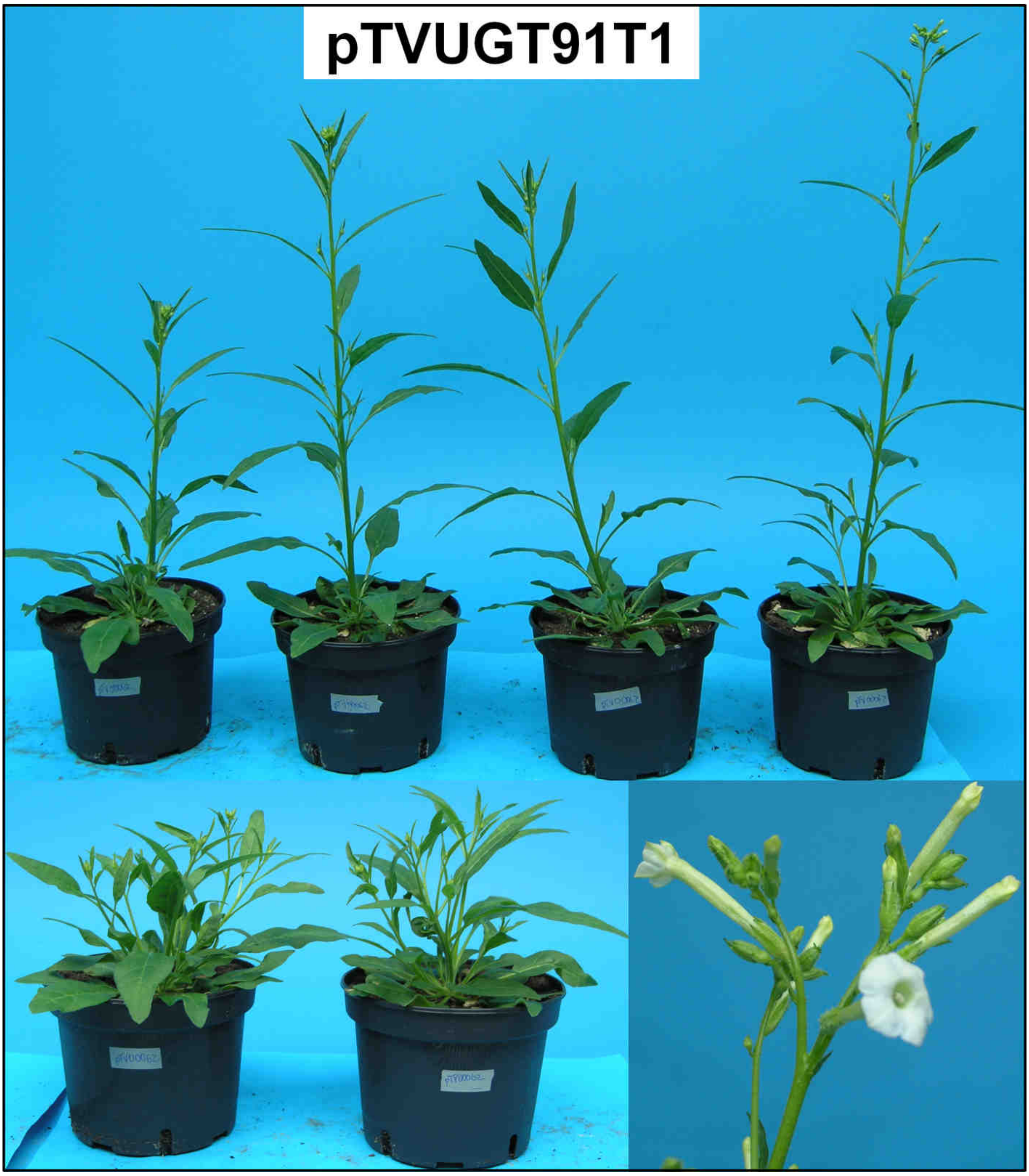


**Supplemental Figure 13a: Morphological characterization of *N. attenuata* plants transiently-silenced in *UGT91T1* expression**

Shown are 37-day-old elongated *N. attenuata* plants transiently-silenced via VIGS in *UGT91T1* expression.

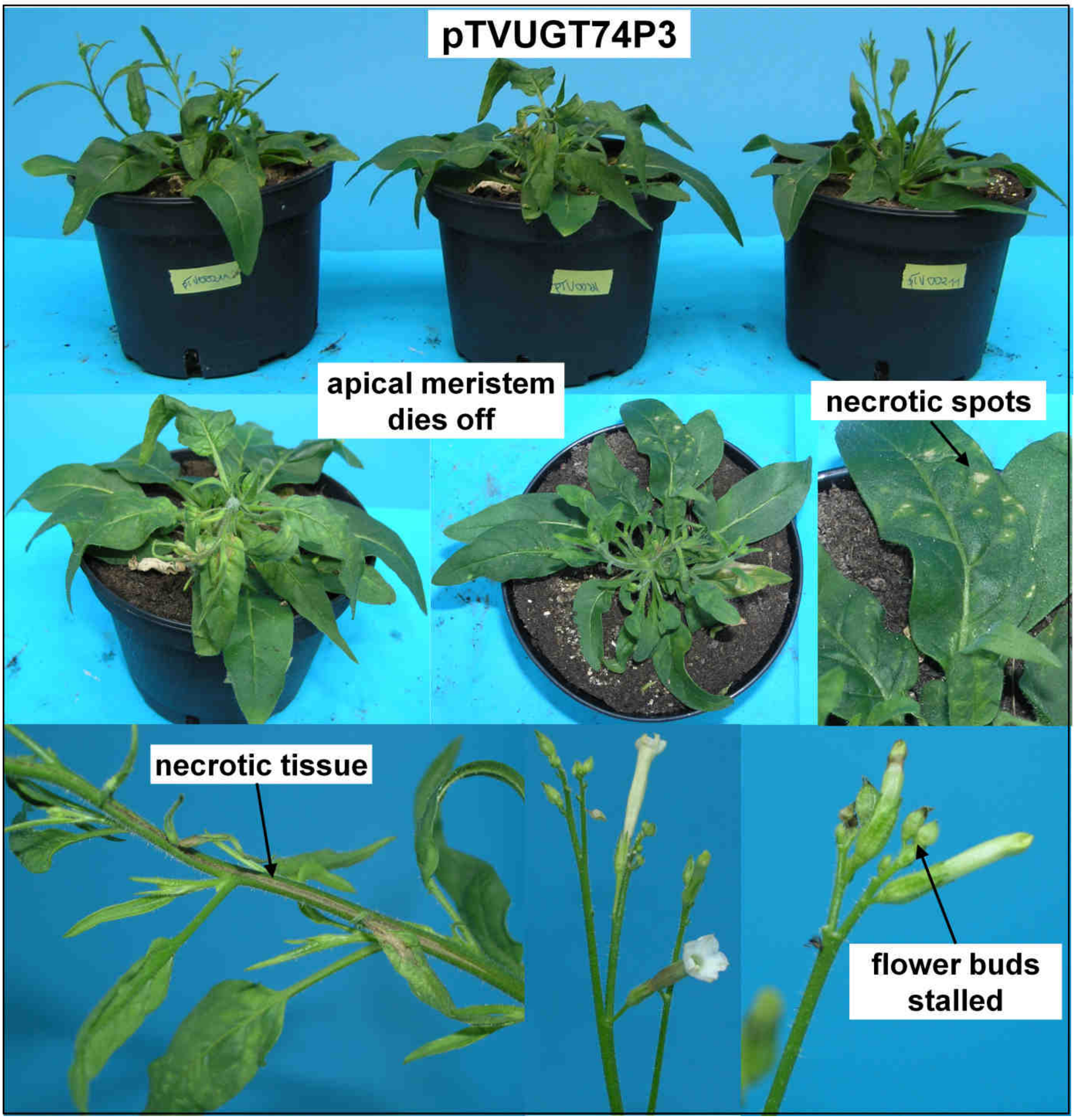


**Supplemental Figure 13b: Morphological characterization of *N. attenuata* plants transiently-silenced in *UGT74P3* expression**

Shown are 37-day-old elongated *N. attenuata* plants transiently-silenced via VIGS in *UGT74P3* expression. Morphological alterations that ranged from the presence of necrotic spots and tissues to dead apical meristems and a high percentage of stalled flower buds.

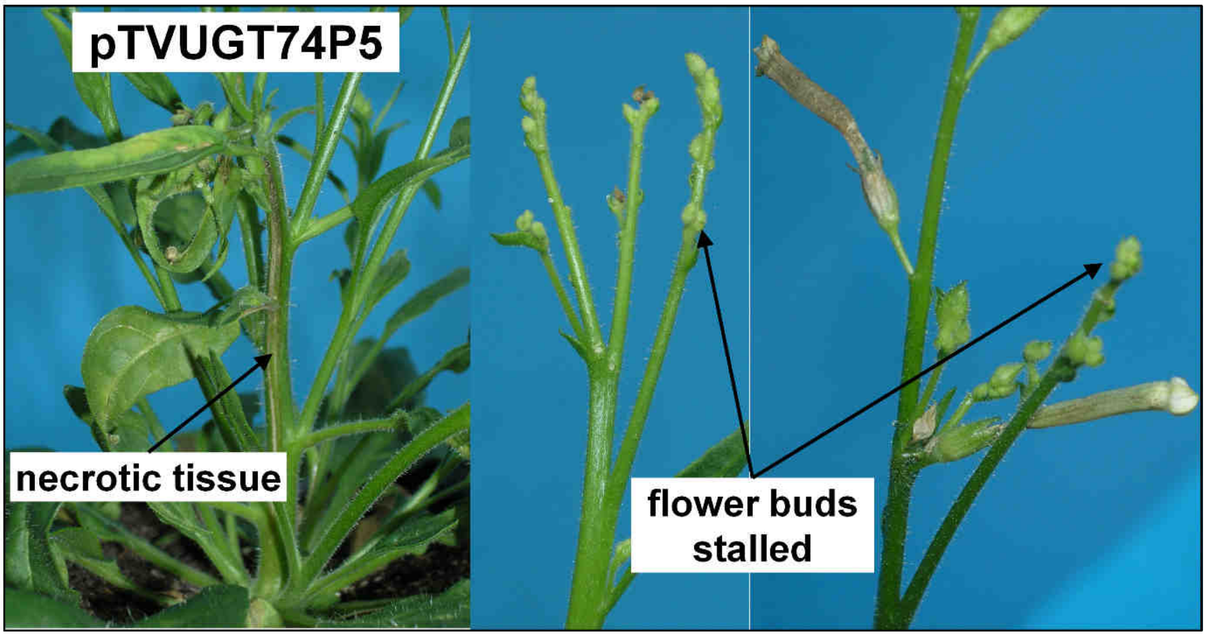


**Supplemental Figure 13c: Morphological characterization of *N. attenuata* plants transiently-silenced in *UGT74P5* expression**

Shown are 37-day-old elongated *N. attenuata* plants transiently-silenced via VIGS in *UGT74P5* expression. Morphological alterations like necrotic tissues and a high percentage of stalled flower buds were observed.

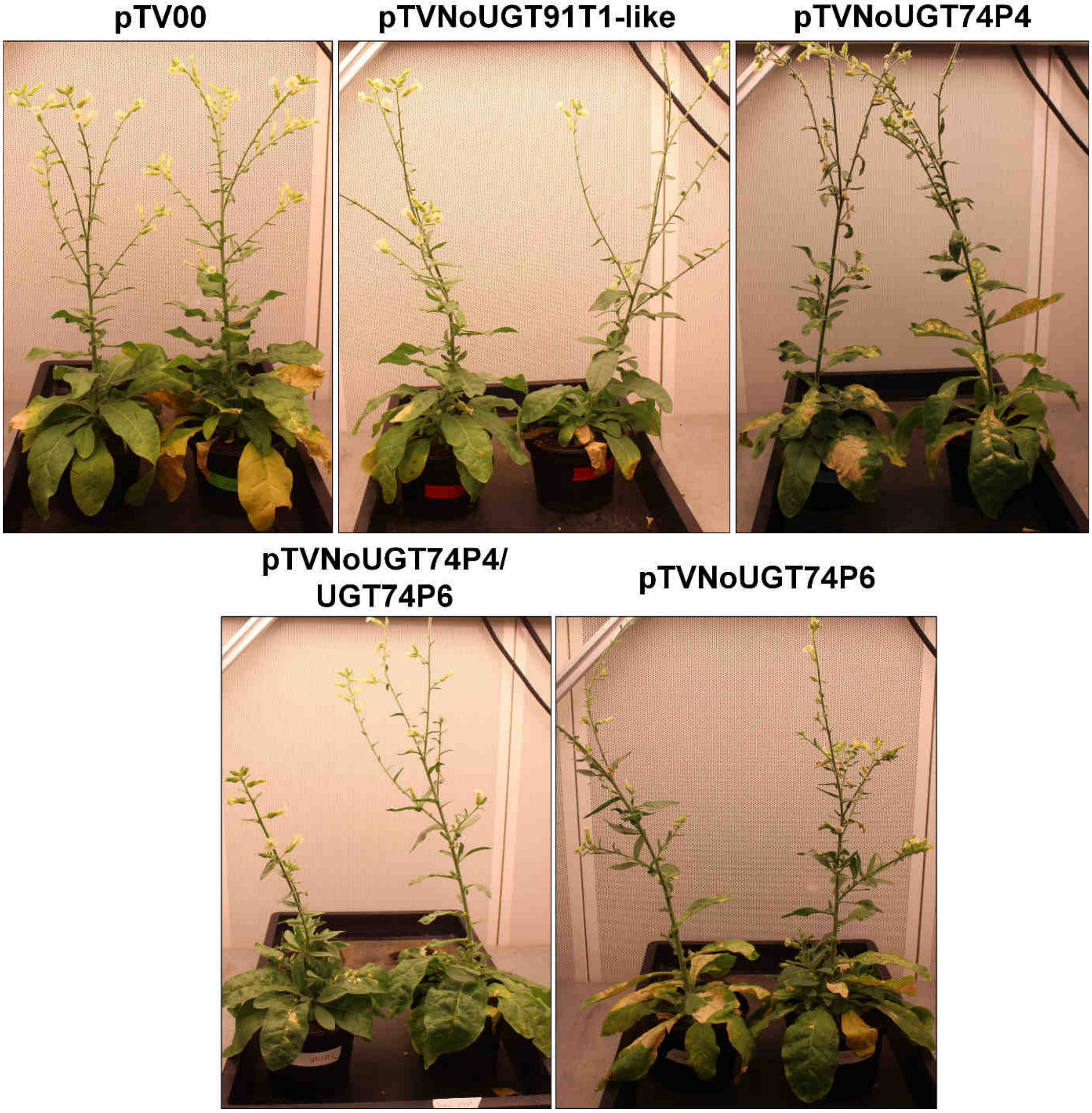


**Supplemental Figure 14: Morphological characterization of *N. obtusifolia* plants transiently-silenced in No*UGT91T1-like*, No*UGT74P4*, No*UGT74P6* and No*UGT74P4*/*UGT74P6* expression**

Shown are 60-day-old flowering *N. obtusifolia* plants transiently-silenced via VIGS in No*UGT91T1-like*, No*UGT74P4*, No*UGT74P6* and No*UGT74P4*/*UGT74P6* expression. Withered flowers were removed. Abundant necrotic spots were only detected in pTVNoUGT74P4 and pTVNoUGT74P6. In contrast to *N. attenuata,* no apical meristem necrosis could be detected.

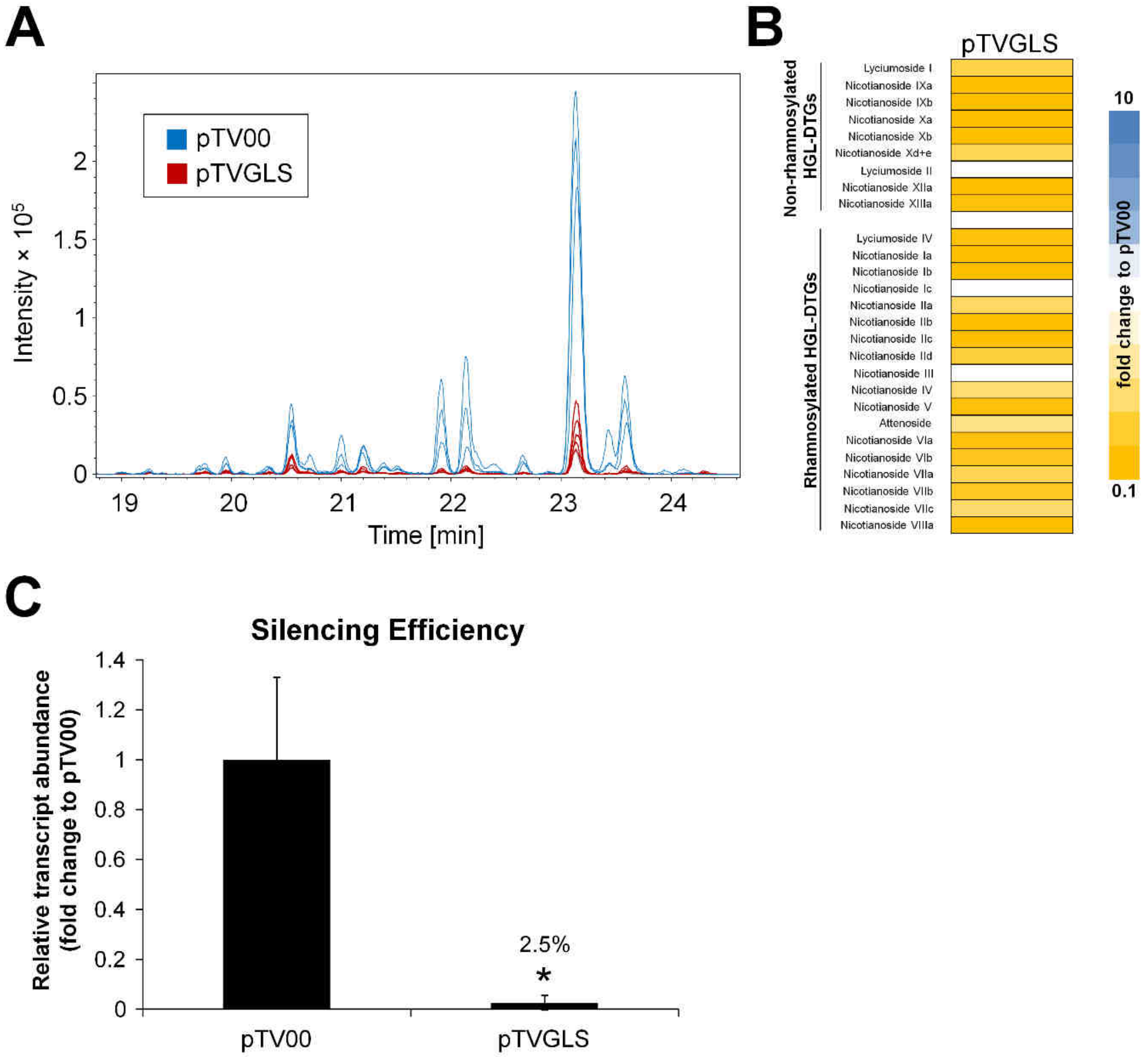


**Supplemental Figure 15: Metabolite profiling and morphological characterization of *N. attenuata* plants transiently-silenced via virus-induced gene silencing (VIGS) of geranyllinalool synthase (GLS)**

A) Shown is the EIC trace for the HGL-DTG aglycone fragment (*m/z* 271.2420) in 37 days old elongated transiently-silenced *N. attenuata* plants impaired in *GLS* expression as well as empty vector control plants (pTV00). B) Heatmap visualization of deregulations of the leaf HGL-DTG profile of pTVGLS transformed plants (N=4). The color gradient visualizes fold changes in individual HGL-DTGs for the GLS-silenced plants compared to the average in the pTV00 empty vector plants. Further details on the abundance of HGL-DTGs can be found in Supplemental Data 2 - GLS. C) Relative transcript abundance (fold change to *N. attenuata Elongation Factor 1α* – Na*ELF1α*) of Na*GLS* in leaves of transiently-silenced *N. attenuata* plants (N=4). Asterisks indicate significant differences between pTV00 empty vector control and pTVGLS (*t*-test, *P ≤ 0.05).

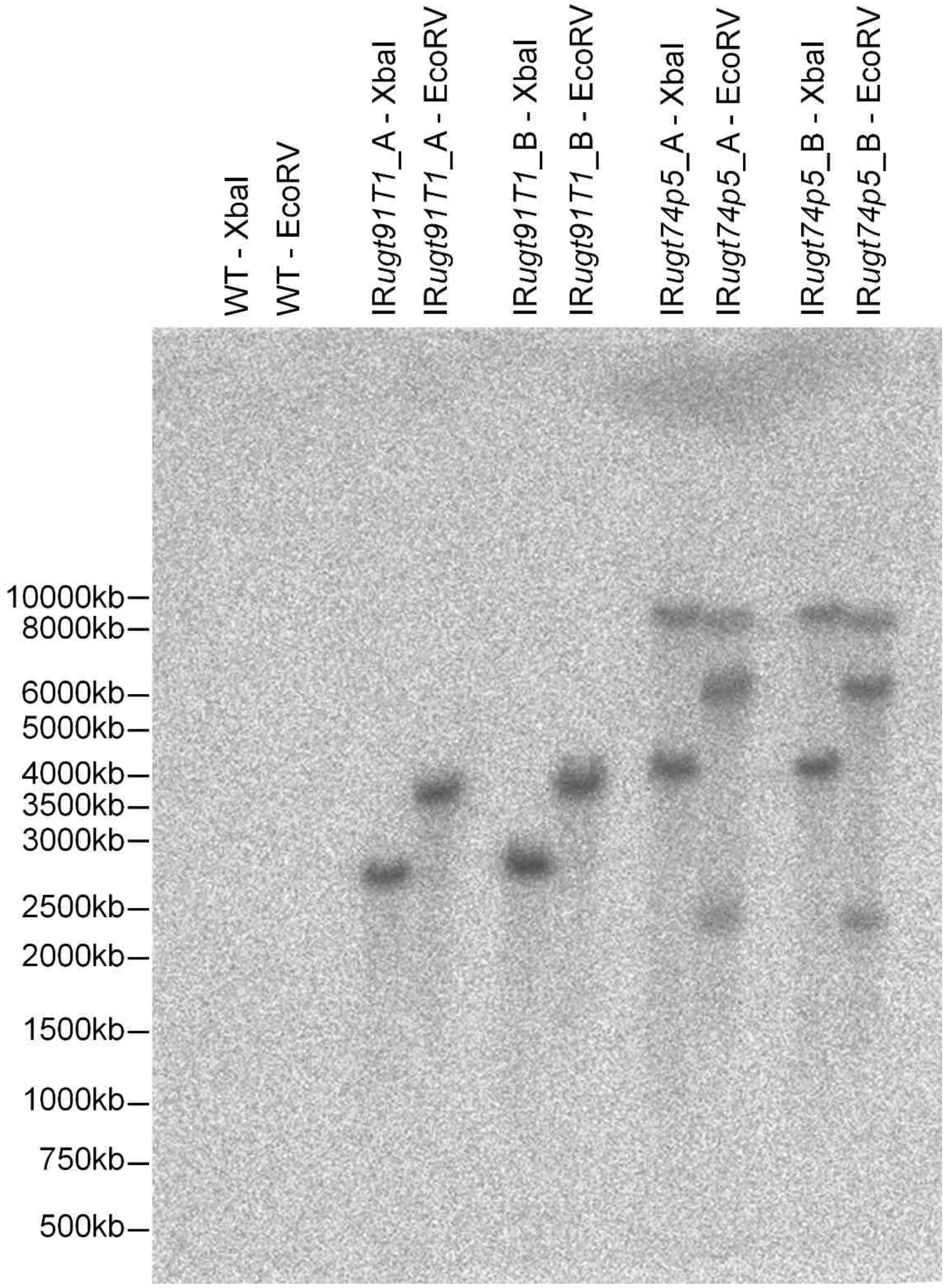


**Supplemental Figure 16: Southern Blot**

Examination of the insertion events of both IR*ugt91t1* and both IR*ugt74p5* transformed lines using XbaI and EcoRV as restriction enzymes.

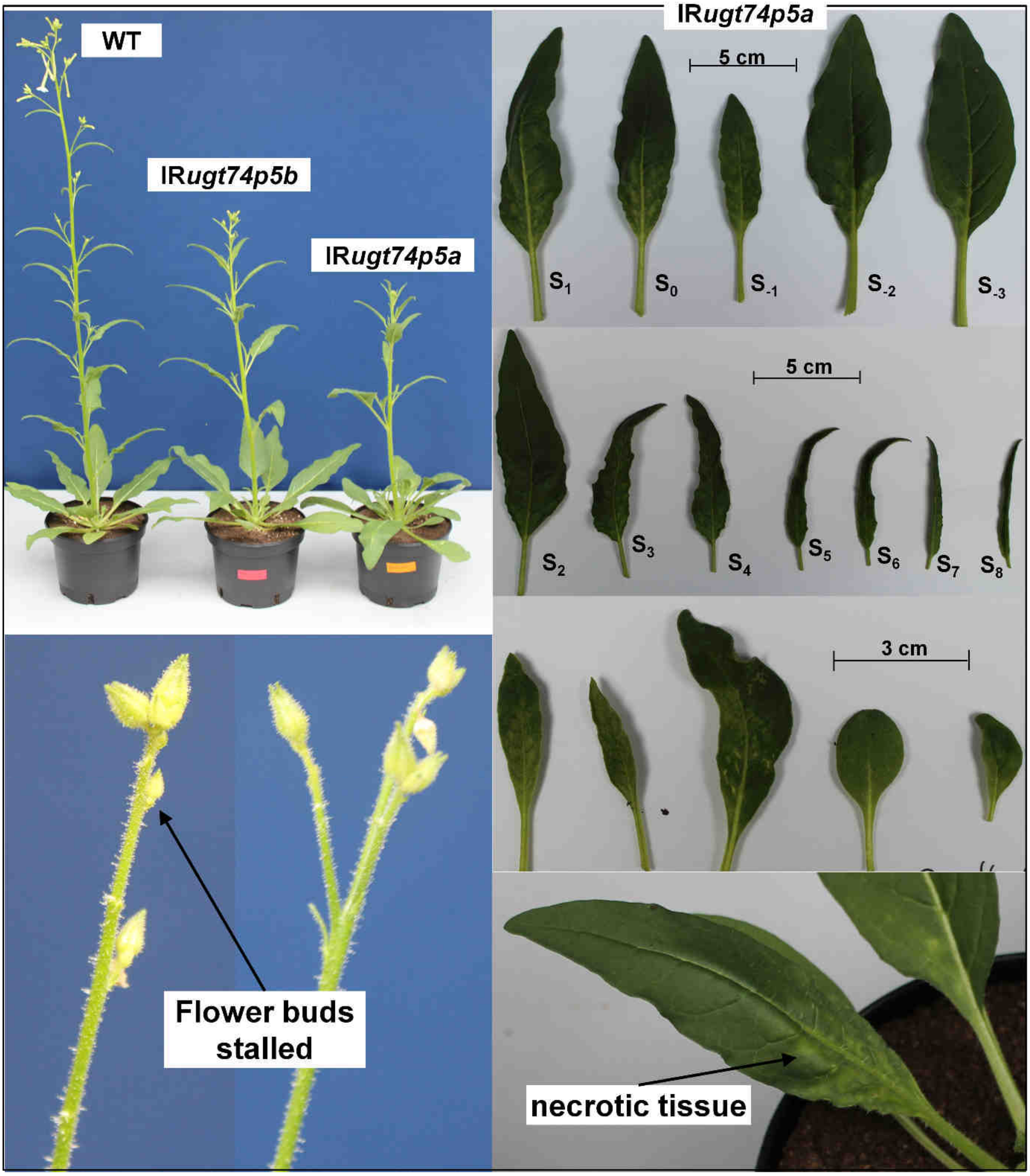


**Supplemental Figure 17a: Morphological characterization of the stable transformed *IRugt74p5* Line A**

Shown are 43-day-old flowering *N. attenuata* plants impaired in *UGT74P5* expression compared to wild type. Morphological alterations of IR*ugt74p5a* ranged from curly deformed rosette leaves with necrotic spots to deformed thin stem leaves. Additionally, several “dwarfish” or succulent round shaped leaves and chlorotic stalled flower buds were observed.

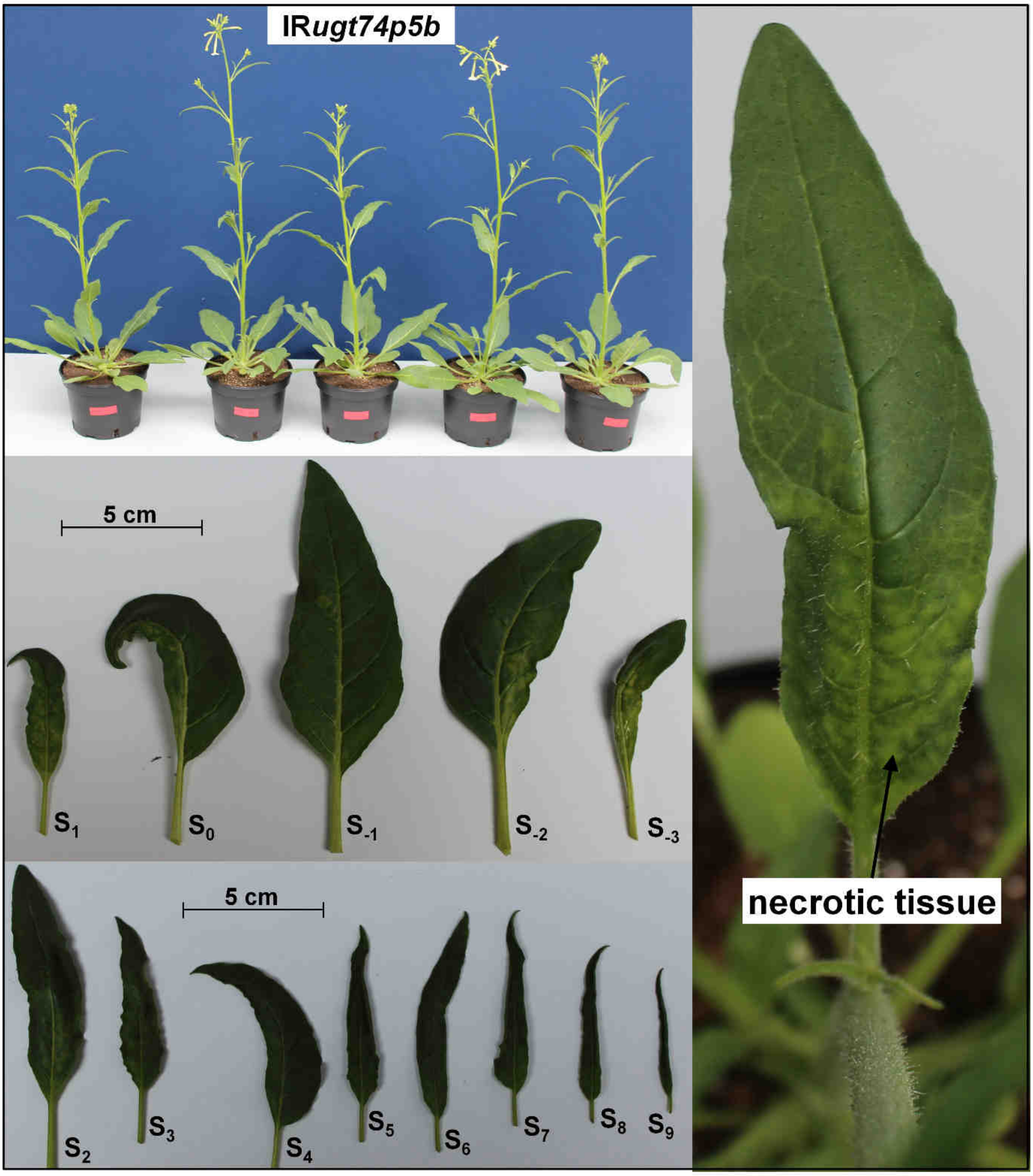


**Supplemental Figure 17b: Morphological characterization of the stable transformed *IRugt74p5* Line B**

Shown are 43-day-old flowering *N. attenuata* plants impaired in *UGT74P5* expression (Line B). Morphological alterations of IR*ugt74p5b* ranged from curly deformed thin and necrotic rosette leaves to deformed thin rippled stem leaves.

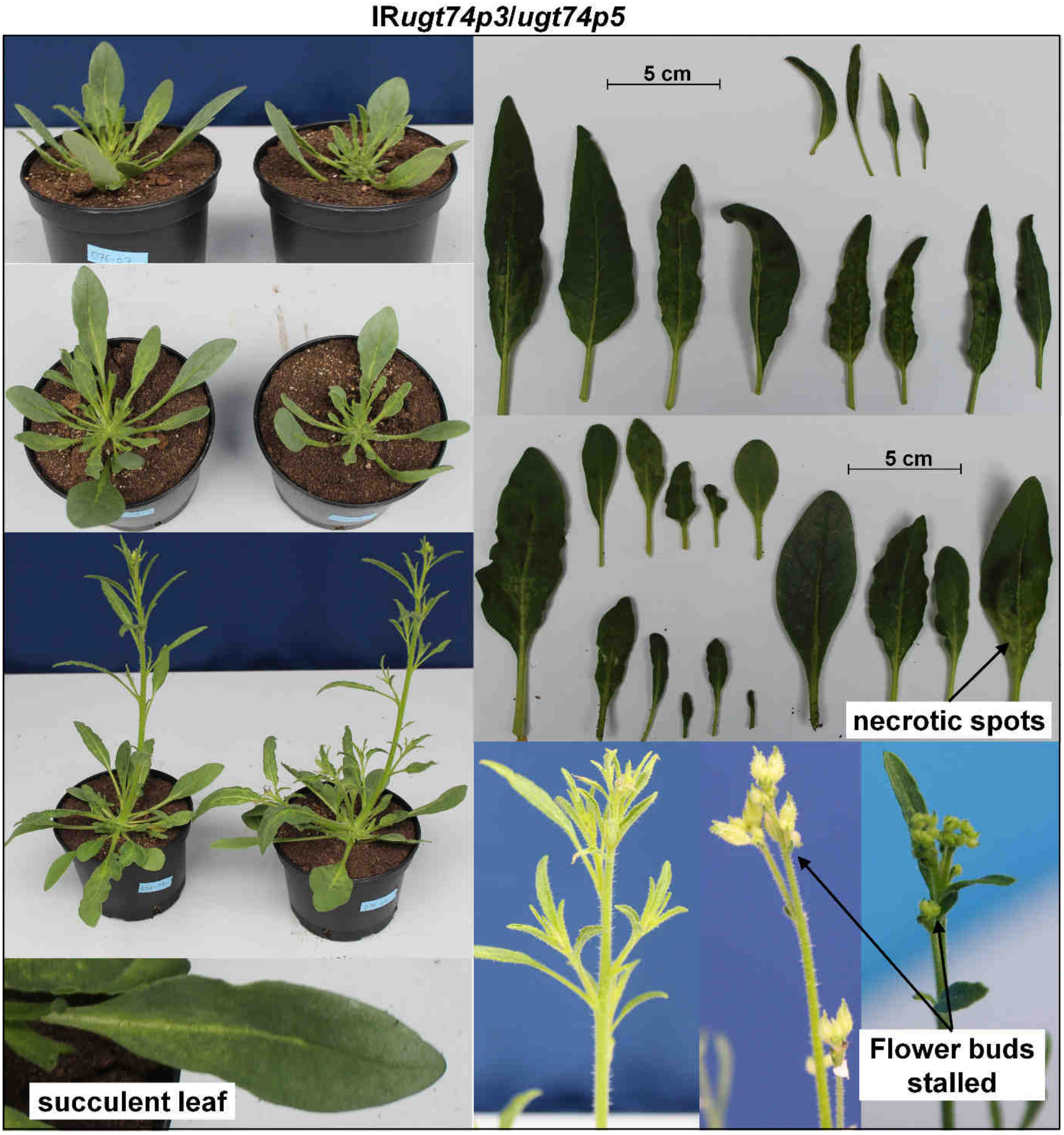


**Supplemental Figure 17c: Morphological characterization of the stable transformed *IRugt74p3/ugt74p5*.**

Shown are 43-day-old flowering *N. attenuata* plants impaired in UGT74P3 and *UGT74P5* expression. Morphological alterations of IR*ugt74p3/ugt74p5* ranged from curly, thin and deformed rosette leaves with necrotic spots to "dwarfish" or succulent round-shaped stem leaves with a high grade of deformation. Furthermore, higher branching grades as well as stalled chlorotic flower buds were observed. Additionally several impaired plants displayed a "dwarf-like" phenotype with small deformed and succulent leaves.

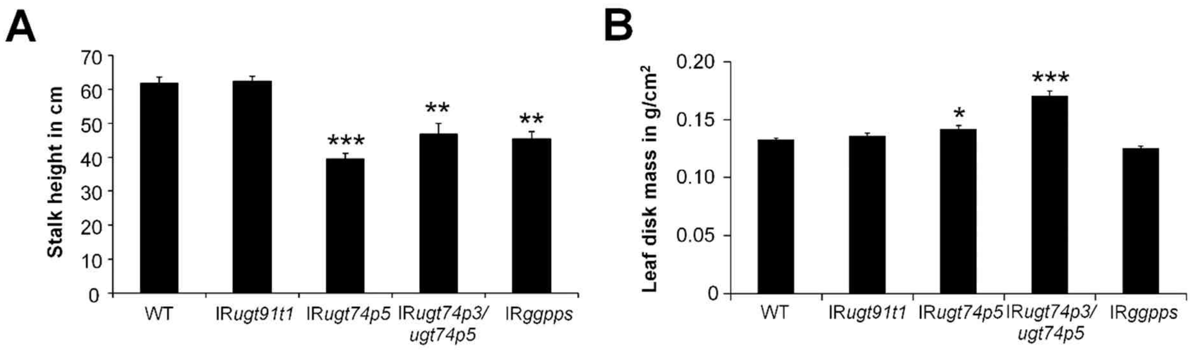


**Supplemental Figure 18: Characterization of growth parameters in IR*ugt91t1*, IR*ugt74p5*, IR*ugt74p3/ugt74p5* and IR*ggpps***

Displayed is the stalk height (A) and leaf disk mass (B) of 40-day-old elongated *N. attenuata* plants stably transformed with IR*ugt91t1* Line A, IR*ugt74p5* Line B, IR*ugt74p3/ugt74p5*, IR*ggpps* at the transition to the flowering stage (N=15). Asterisks indicate significant differences between wild type control and stably-silenced lines (*P≤ 0.05, ** P < 0.01, *** P < 0.001).

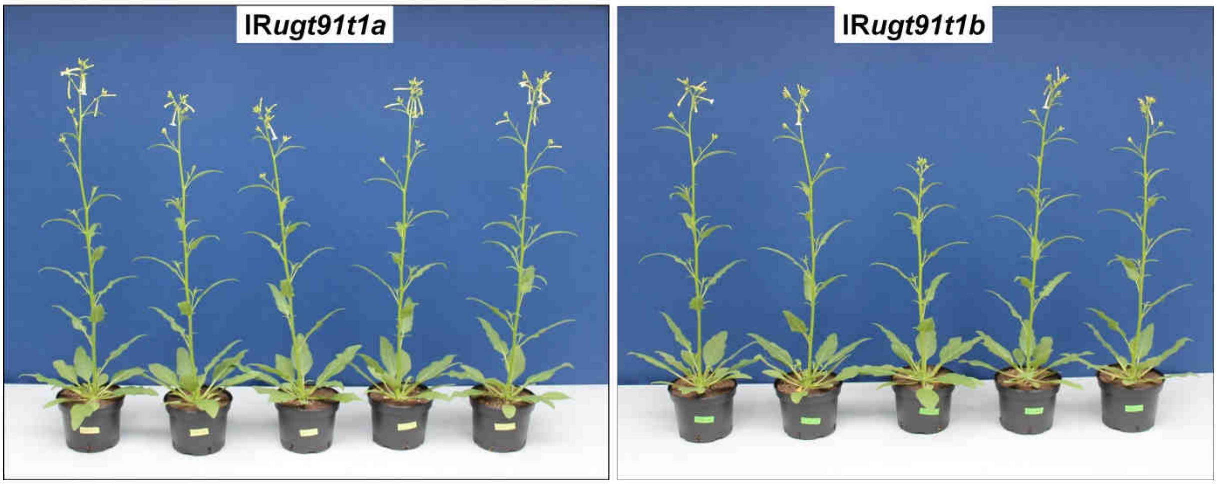


**Supplemental Figure 19: Morphological characterization of IR*ugt91t1***

Shown are 43-day-old *N. attenuata* plants of both stably silenced IR*ugt91t1* lines.

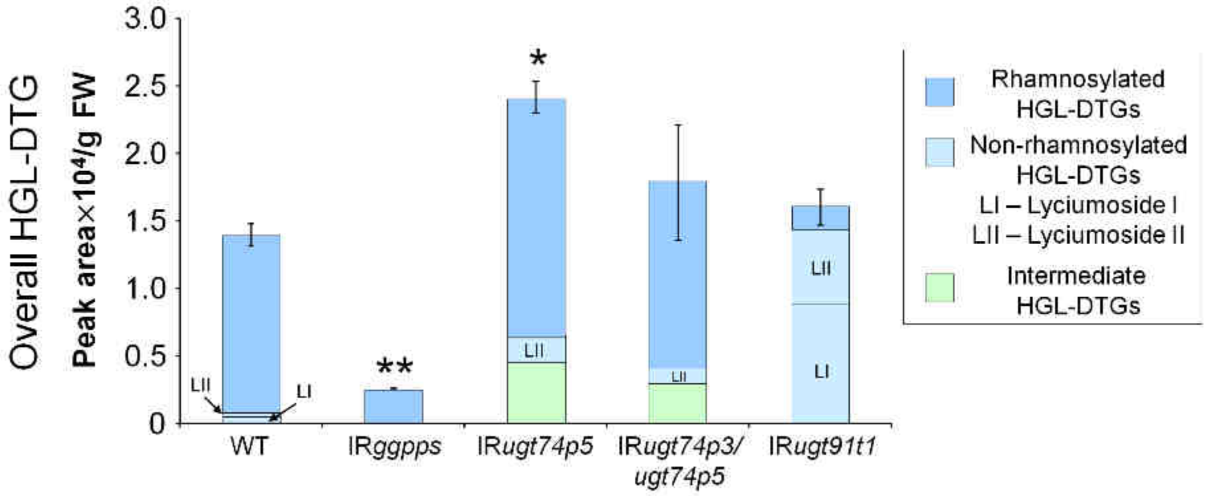


**Supplemental Figure 20: Overall abundance of HGL-DTGs**

Shown is the overall area/g FW of rhamnosylated, non-rhamnosylated and intermediate HGL-DTGs in the different stable lines (N=5). LI and LII represent the most abundant non-rhamnosylated HGL-DTGs lyciumoside I and lyciumoside II and their malonylated forms in the different lines. Asterisks indicate significant differences between WT and stably-silenced lines (*P ≤ 0.05, ** P < 0.01).

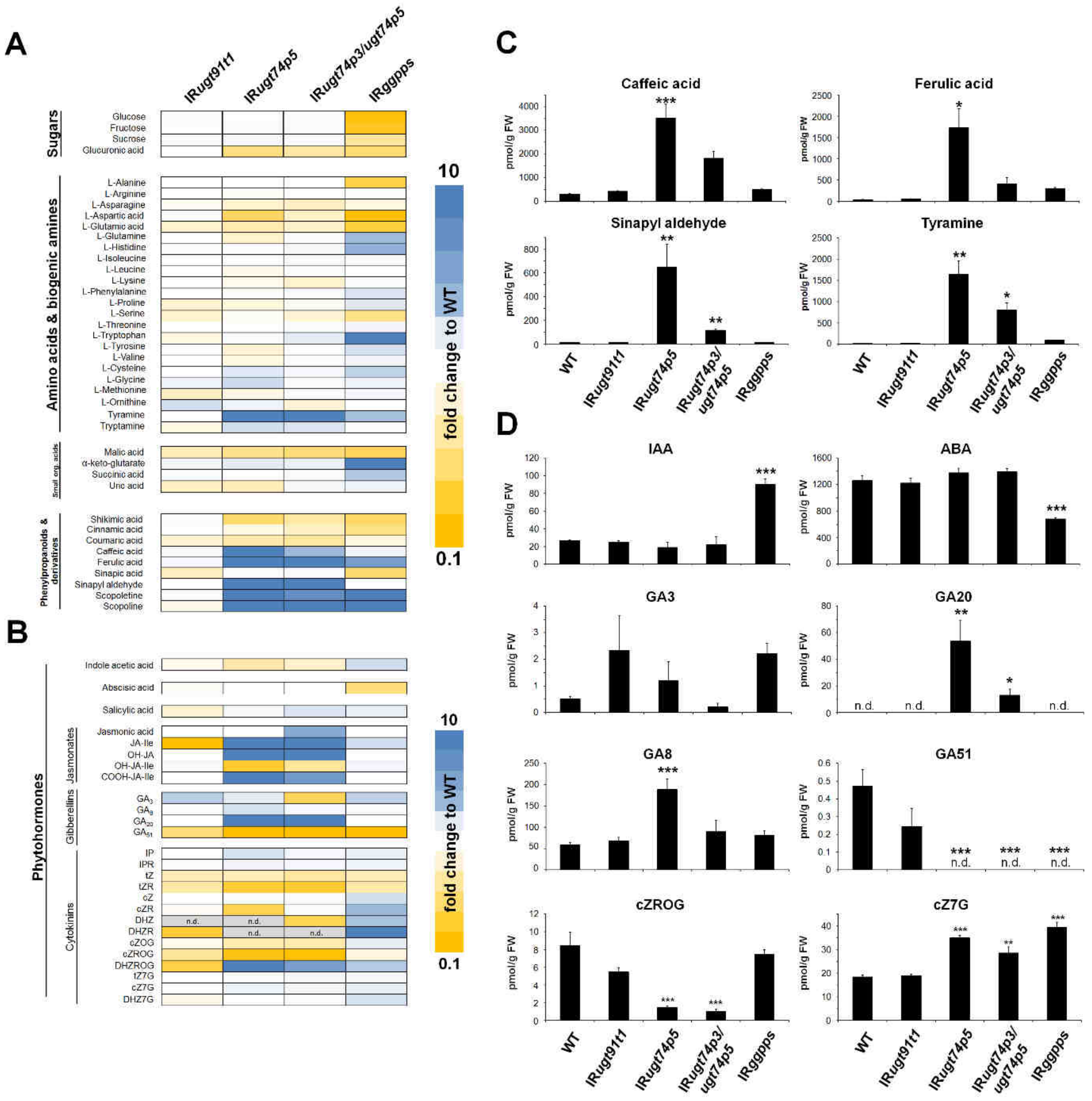


**Supplemental Figure 21: Disrupting HGL-DTG glycosylation reorganizes general, specialized and hormonal metabolic pathways**

Heatmap visualization of deregulations in the leaves' A) + C) general/specialized and B) + D) hormonal metabolic profiles of IR*ugt91t1*, IR*ugt74p5*, IR*ugt74p3/ugt74p5* and IR*ggpps* plants (N=5). Color gradients visualize fold changes in individual metabolites for each of the stable lines compared to the average in the WT plants. Asterisks indicate significant differences between WT control and stable transformants (*P ≤ 0.05, ** P < 0.01, *** P < 0.001).

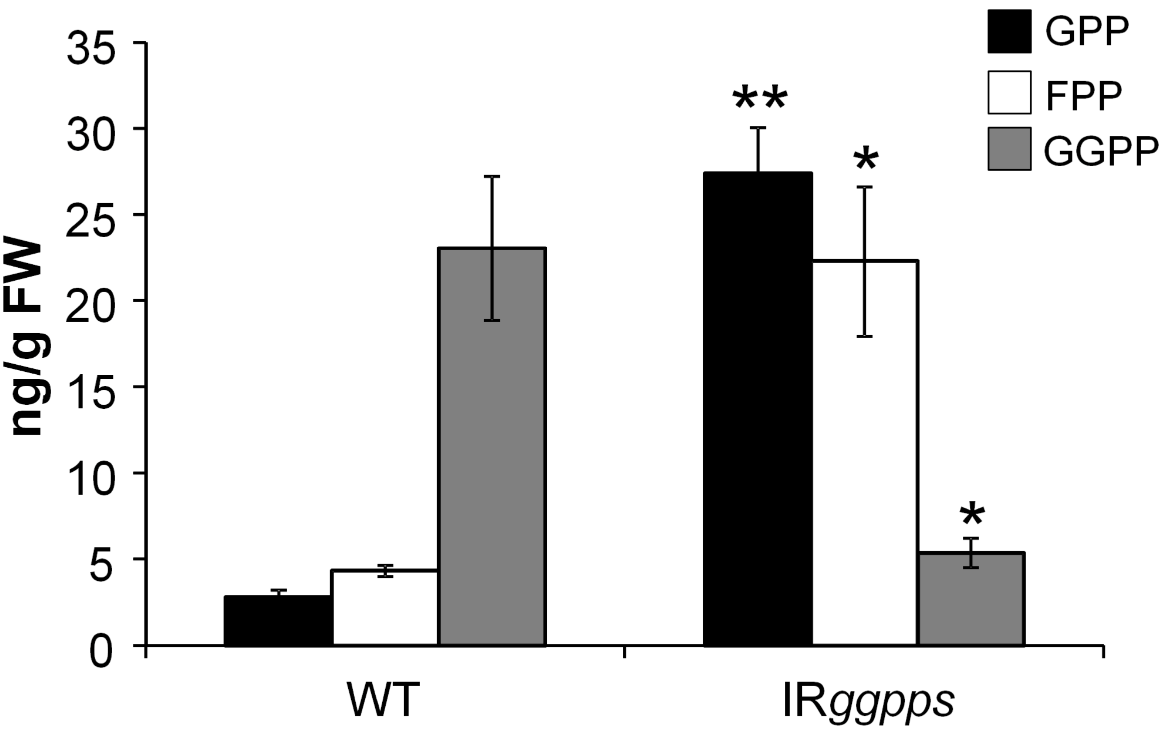


**Supplemental Figure 22: Characterization of free prenyldiphosphates in IR*ggpps* and WT**

Shown is the amount of the free prenyldiphosphates, GDP, FDP and GGDP, in leaf tissue of IR*ggpps* and wild type *N. attenuata* plants (N=4). Asterisks indicate significant differences between WT and IR*ggpps* (*t*-test, *P ≤ 0.05, ** P < 0.01).

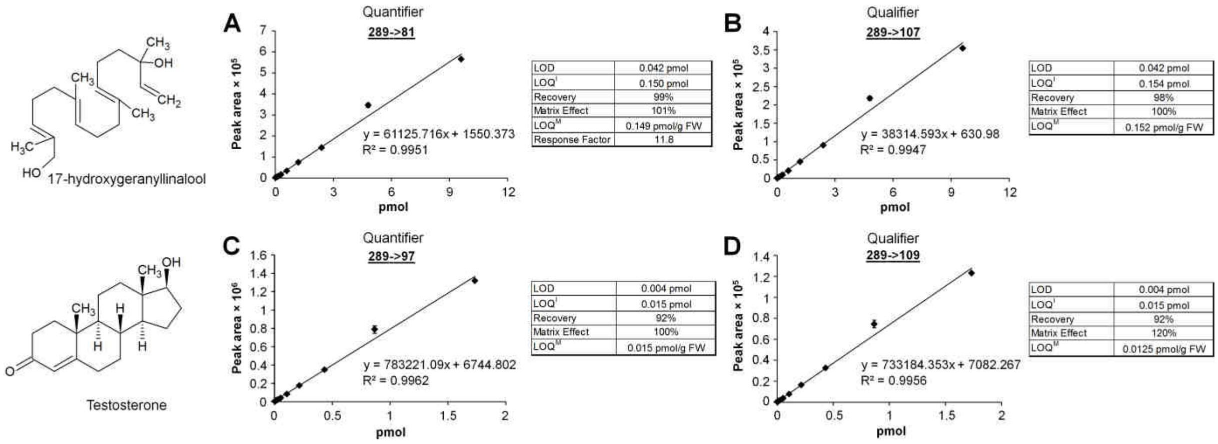


**Supplemental Figure 23: Quantitative 17-HGL method**

Shown are the standard curves for the quantifiers (A and C) and qualifiers (B and D) for 17-HGL as well as the testosterone internal standard. Method parameters like limit of detection (LOD), limit of quantification (LOQ), recovery, matrix effect, quantification limit of the instrument ($LOQ^M$) and the response factor are presented.

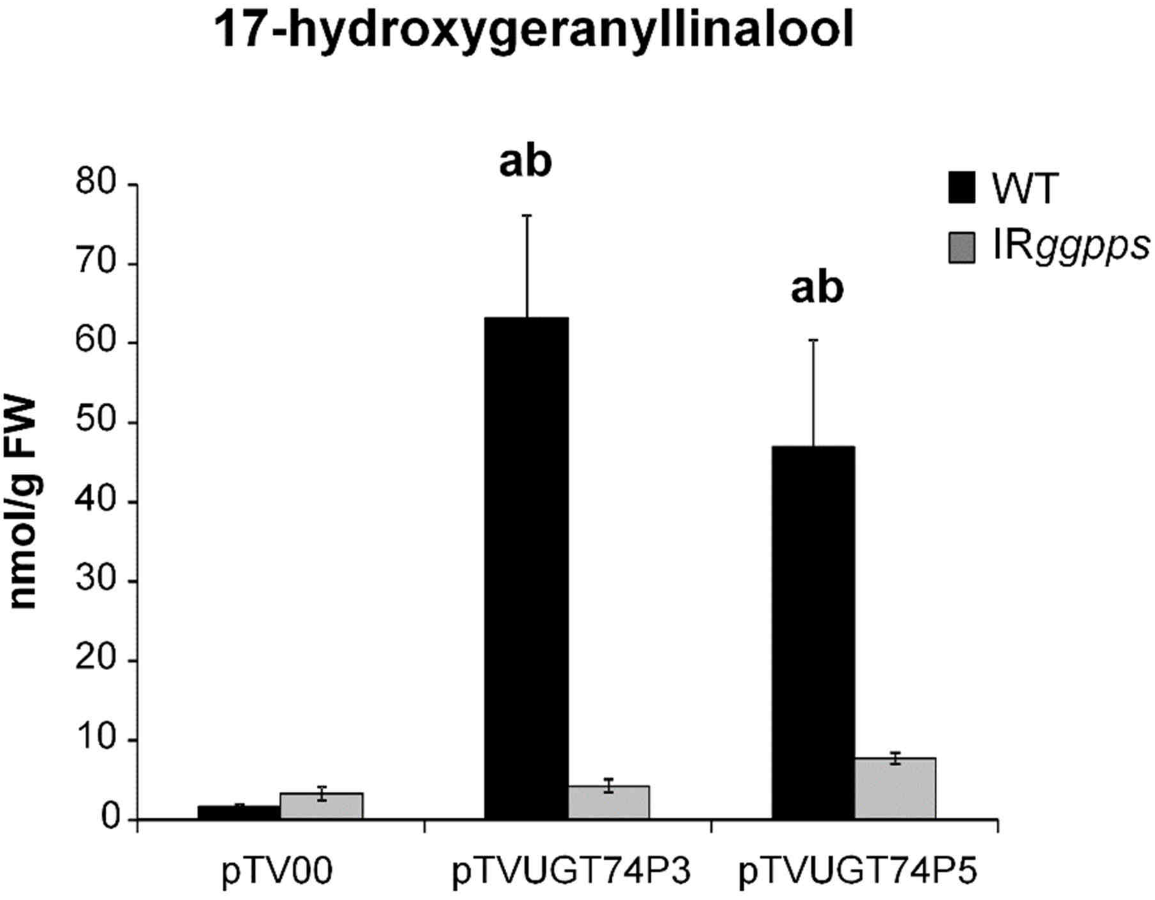


**Supplemental Figure 24: 17-HGL concentration in IR*ggpps* and WT plants transiently transformed with pTV00, pTVUGT74P3 and pTVUGT74P5**

Shown is the amount of the 17-HGL in leaf tissue of WT and IR*ggpps* plants transiently transformed with pTV00, pTVUGT74P3 and pTVUGT74P5 (N=5). Letters indicate significant differences from the pTV00 empty vector control (a) and between IR*ggpps* and wild type (b) ($P \leq 0.05$).

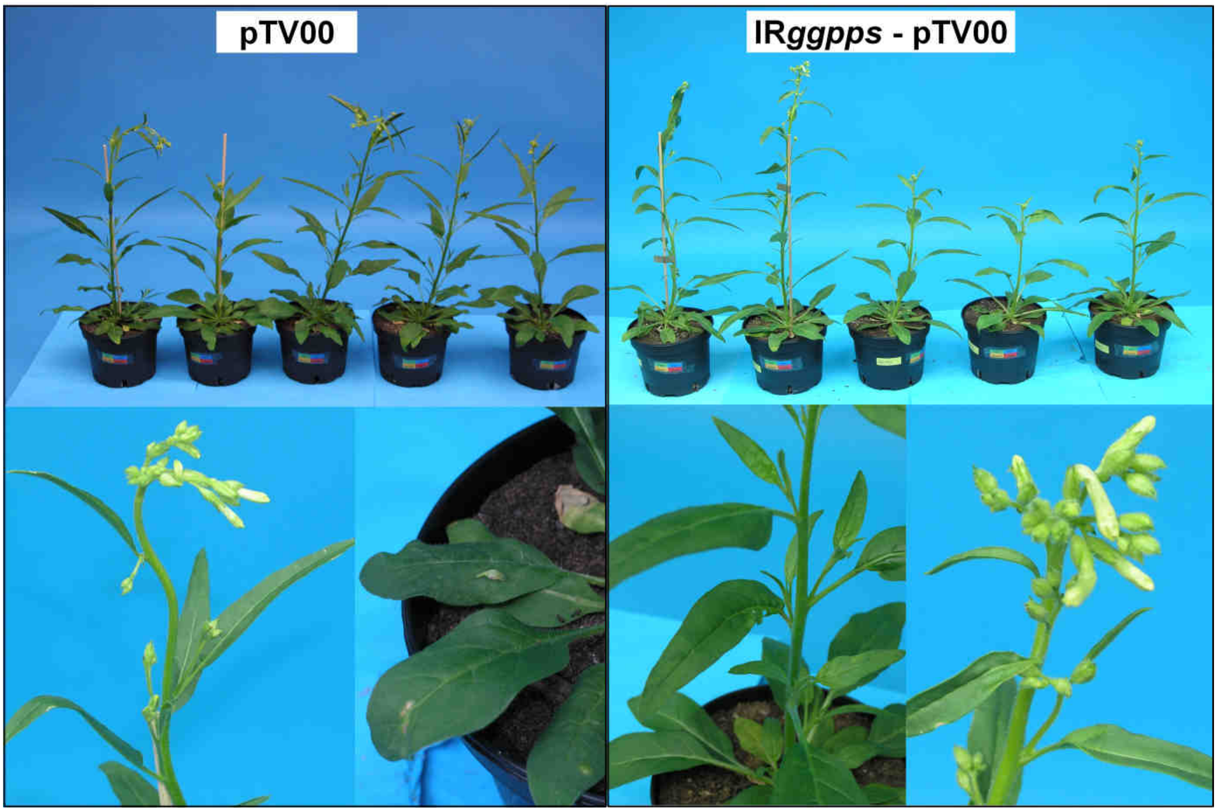


**Supplemental Figure 25: Characterization of the morphological phenotype of WT and IR*ggpps* plants transiently transformed with pTV00.**

Shown are 42-day-old *N. attenuata* plants impaired in *GGPPS* expression and WT. Both are transiently transformed with the empty vector control and display typical morphological alterations like curly leaves due to the VIGS procedure.

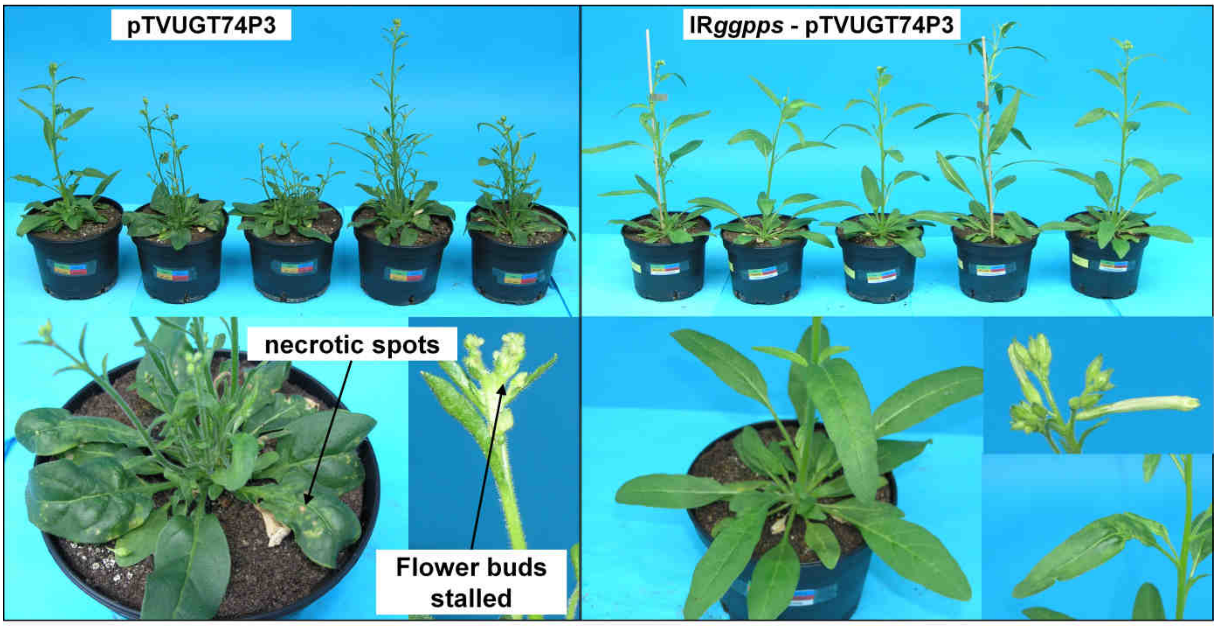


**Supplemental Figure 26: Characterization of the morphological phenotype of WT and IR*ggpps* plants transiently transformed with pTVUGT74P3.**

Shown are 42-day-old *N. attenuata* plants impaired in *GGPPS* expression and WT. Both are transiently-transformed with the pTVUGT74P3 vector harboring a glucosyltransferase responsible for the synthesis of HGL-DTGs. WT VIGS plants display severe morphological alterations ranging from necrotic spots, lower stem height, enhanced branching and stalled or aborted flower buds. IR*ggpps* plants transiently transformed with pTVUGT74P3 only show slightly curly leaves.

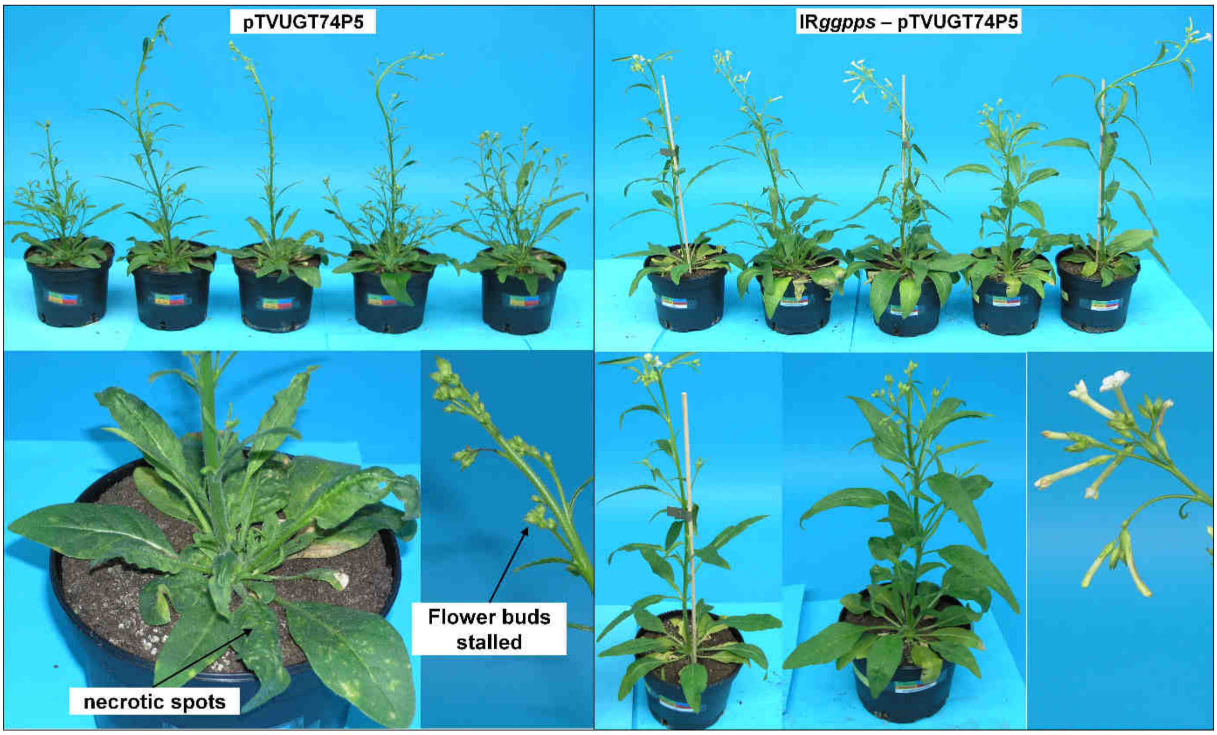


**Supplemental Figure 27: Characterization of the morphological phenotype of WT and IR*ggpps* plants transiently transformed with pTVUGT74P5.**

Shown are 42-day-old *N. attenuata* plants impaired in *GGPPS* expression and WT. Both are transiently-transformed with the pTVUGT74P5 vector harboring a glucosyltransferase responsible for the synthesis of HGL-DTGs. WT VIGS plants display severe morphological alterations ranging from necrotic spots, enhanced branching and stalled or aborted flower buds. IR*ggpps* VIGS plants using pTVUGT74P5 only show slightly curly leaves.
